# The DNA mimic protein BCAS0292 is involved in the regulation of virulence of *Burkholderia cenocepacia*

**DOI:** 10.1101/2020.08.06.226779

**Authors:** Ruth Dennehy, Simon Dignam, Sarah McCormack, Maria Romano, Yueran Hou, Laura Ardill, Matthew X. Whelan, Zuzanna Drulis-Kawa, Tadhg Ó Cróinín, Miguel A. Valvano, Rita Berisio, Siobhán McClean

**Affiliations:** Centre of Microbial Host Interactions, Institute of Technology Tallaght, Old Blessington Road, Dublin 24; School of Biomolecular and Biomedical Science, University College Dublin, Belfield, Dublin 4; Institute of Biostructures and Bioimaging, National Research Council, Via Mezzocannone 16, I-80134 Naples, Italy; Institute of Genetics and Microbiology, University of Wroclaw, 51-148 Wroclaw, Poland; Wellcome-Wolfson Institute for Experimental Medicine, Queen’s University Belfast, Belfast, BT9 7BL, United Kingdom

## Abstract

Adaptation of opportunistic pathogens to their host environment requires reprogramming of a vast array of genes to facilitate survival in the host. *Burkholderia cenocepacia*, a Gram-negative bacterium that colonizes environmental niches, is exquisitely adaptable to the hypoxic environment of the cystic fibrosis lung and survives in macrophages. *B. cenocepacia* possesses a large genome encoding multiple virulence systems, stress response proteins and a large locus that responds to low oxygen. We previously identified BCAS0292, an acidic protein encoded on replicon 3. Deletion of the BCAS0292 gene resulted in altered abundance of >1000 proteins; 46 proteins became undetectable while 556 proteins showed ≥1.5-fold reduced abundance, suggesting BCAS0292 is a global regulator. Moreover, the ΔBCAS0292 mutant showed a range of pleiotropic effects: virulence, host-cell attachment and motility were reduced, antibiotic susceptibility was altered and biofilm formation enhanced. Its growth and survival were impaired in 6% oxygen. Structural analysis revealed BCAS0292 presents a dimeric β-structure with a negative electrostatic surface. Further, the ΔBCAS0292 mutant displayed altered DNA supercoiling, implicated in global regulation of gene expression. We propose that BCAS0292 acts as a DNA-mimic, altering DNA topology and regulating the expression of multiple genes, thereby enabling the adaptation of *B. cenocepacia* to highly diverse environments.

## Introduction

Pathogenic bacteria experience a broad range of stress conditions within their hosts, including changes in temperature, oxygen levels, pH, nutrient limitation and antimicrobial molecules. Bacteria respond and adapt to these stresses by reprogramming the expression of large numbers of genes. Environmental opportunistic pathogens, such as *Pseudomonas aeruginosa*, *Acinetobacter baumannii*, *Mycobacterium marinum, Bacillus subtilis, Vibrio vulnificus* and members of genus Burkholderia preferentially colonise environmental niches, including soil and water courses and their success as human pathogens relies on their plasticity to adapt to varying conditions. Antimicrobial resistance (AMR) ranks among the most important threats to human health. Among the most problematic AMR bacteria, the so-called ESKAPE pathogens, such as *P. aeruginosa*, are opportunistic pathogens that are highly adaptable in their ability to colonise a diverse range of niches including the human host ((De Oliveira *et al.*, 2020). Insights into the adaptability of these successful opportunistic pathogens will enable the development of alternatives to antimicrobial therapies.

One example of an exquisitely adaptable pathogen which colonises very diverse ecological niches ranging from contaminated soils, water sources and pharmaceutical plants to the human lung is the Gram-negative genus Burkholderia. These bacteria possess large genomes (~8Mb) and have enormous genetic potential for adaptation within the host. Within this genus, *Burkholderia cepacia* complex (Bcc) is a group of closely related bacteria which causes opportunistic chronic infections in people with CF (Mahenthiralingam *et al.*, 2008). The complex currently comprises 24 species (Peeters *et al.*, 2013, Bach *et al.*, 2017) which are highly resistant to different classes of antibiotics (Zhou *et al.*, 2007). Some members of the complex, notably *B. cenocepacia* and *B. multivorans* have been shown to survive in macrophages and to attach to human epithelial cells (Saini *et al.*, 1999, Cullen *et al.*, 2017, Schmerk and Valvano, 2013, Valvano, 2015, Caraher *et al.*, 2007). Further, *B. cenocepacia* can persist in 6% oxygen by means of a co-regulated 50-gene cluster of genes, designated the low-oxygen activated (*lxa*) locus (Sass *et al.*, 2013) which we have shown to be upregulated during chronic infection in response to the hypoxic environment of the CF lung (Cullen *et al.*, 2018).

Previously, we identified an immunogenic protein encoded by the BCAS0292 gene in *B. cenocepacia* using serum from Bcc-colonised people with cystic fibrosis (CF) (Shinoy *et al.*, 2013). This gene encodes a hypothetical protein and is located within a virulence gene cluster on replicon 3 of the *B. cenocepacia* genome (Figure 1), which also includes *aidA* (BCAS0293). The *aidA* gene, a paralog of BCAS0292, encodes a protein that is required for virulence in *C. elegans* (Huber *et al.*, 2004). Both BCAS0292 and AidA proteins have been classified as PixA proteins, a poorly understood group of inclusion body proteins associated with cells in stationary phase (Winsor *et al.*, 2008, Goetsch *et al.*, 2006). Both BCAS0292 and *aidA* are the most highly quorum sensing regulated genes in *cepR* and *cepRcciIR* mutants (O’Grady *et al.*, 2009). Both genes are also involved in adaptation to the host in a rat chronic infection model (O’Grady and Sokol, 2011). Further, transcripts of both genes are upregulated under low oxygen conditions, and BCAS0292 is also the most upregulated gene in stationary phase, suggesting it may be involved in the stress response of *B. cenocepacia* during anoxic and low nutrient conditions (Sass *et al.*, 2013). Overall, these studies suggest a role for the BCAS0292 protein in the stress response of *B. cenocepacia*; however, its specific function remained unknown. In this work, we constructed a ΔBCAS0292 deletion mutant strain to investigate its role in *B. cenocepacia* pathogenesis. We demonstrate that loss of BCAS0292 alters the abundance of more than a thousand proteins, suggesting the protein is involved in global regulation of gene expression. The deletion results in a range of pleiotropic effects in the mutant strain. We also provide evidence indicating the protein has surface characteristics comparable to known DNA mimic proteins and propose that BCAS0292 functions as a DNA mimic by altering the state of DNA supercoiling.

**Figure 1.**
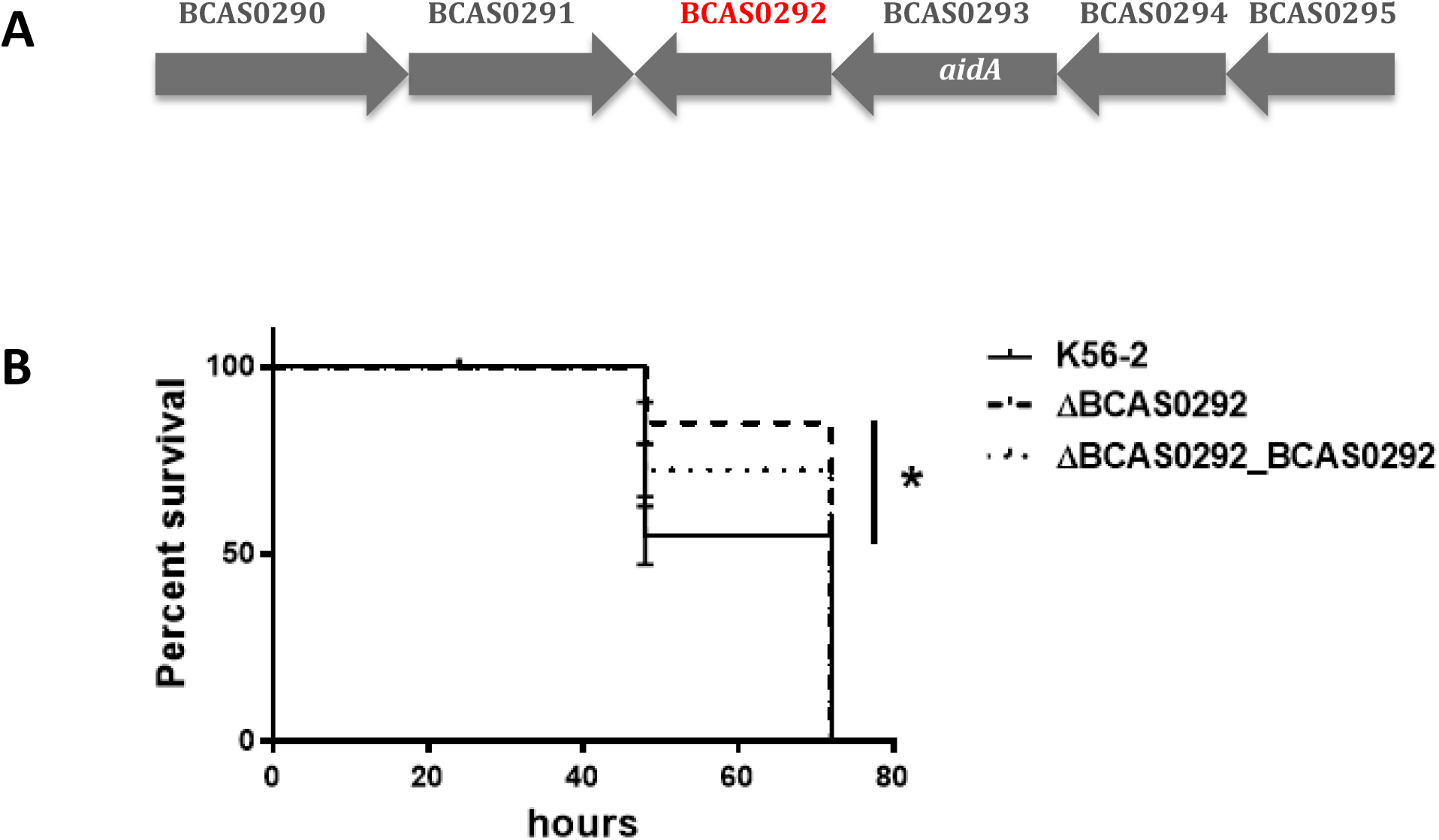
**(A) BCAS0292 gene cluster.** Genetic organisation of the gene cluster encoding the BCAS0292 and AidA proteins in *B. cenocepacia* J2315 and K56-2. The direction of transcription of each gene is indicated by the grey arrows. BCAS gene designations are according to annotation of the *B. cenocepacia* J2315 genome (Holden *et al.*, 2009). (B) **Effect of BCAS0292 gene deletion on virulence.** Kaplan Meier plot showing *G. mellonella* survival following exposure 2 CFU of K56-2 WT, ΔBCAS0292 mutant or Δ0292_0292 complement strain. Data collated from three independent experiments. * Statistical difference between K56-2 WT and Δ*BCAS0292;* p = 0.0036

## Results

BCAS0292 belongs to the PixA protein family. These inclusion body proteins, first identified in *Xenorhabdus nematophila*, are typically between 173 and 191 amino acids in length (Goetsch *et al.*, 2006). They are elevated in cells transitioning to stationary phase (Goetsch *et al.*, 2006), but are poorly understood. BCAS0292 has a calculated molecular weight of 19.2 kDa and a calculated pI of 4.81 and has 34 homologs in several sequenced *B. cenocepacia* strains, and in *B. anthina*, *B. cepacia*, *B pyrrocinia* and other *Burkholderia* species (Winsor *et al.*, 2008); however, a BlastP search showed that it is not found outside *Burkholderia* (Altschul *et al.*, 1997). The BCAS0292 gene is encoded in a two-gene operon with *aidA* (O’Grady *et al.*, 2009) (Figure 1A). AidA is also classified as a PixA protein while it also shares sequence similarity with PixB of *X. nematophila* (Lucas *et al.*, 2018). The *aidA* gene is more widely distributed across *Burkholderia* species with 417 orthologs identified to date (Winsor *et al.*, 2008). AidA also shares a high level of identity with PixB proteins across Gammaproteobacteria, Alphaproteobacteria, Acinobacteria and species (Lucas *et al.*, 2018). While BCAS0292 is immunoreactive, indicating that it is expressed during human infection, we did not detect AidA in our immunoproteomic experiments (Shinoy *et al.*, 2013). To investigate the potential function of BCAS0292 in infection, we constructed a deletion mutant of BCAS0292 in K56-2, a *B. cenocepacia* strain that is amenable to genetic manipulation and a complemented strain (referred to as ΔBCAS0292 and Δ0292_0292, respectively) and compared their phenotypes to the wild type (WT) strain.

### Virulence of K56-2 WT and ΔBCAS0292 strains in *Galleria mellonella* infection model

To examine any potential role in pathogenicity, the virulence of the K56-2 ΔBCAS0292 mutant strain and the K56-2 WT strain were compared using the *G. mellonella* wax moth larvae infection model. K56-2 WT is highly virulent in this model, which is demonstrated by the very low mean LD_50_ value of 1.7 CFU at 48 h. Deletion of the BCAS0292 gene resulted in a moderate reduction in virulence (Figure 1b, p = 0.036) which was partially rescued in the Δ0292_0292 complemented strain, suggesting that BCAS0292 contributes to *B. cenocepacia* virulence in this model.

### Potential role in host-cell interactions

We have previously shown that other immunogenic proteins, including peptidoglycan associated lipoprotein are involved in attachment to host epithelial cells (McClean *et al.*, 2016, Dennehy *et al.*, 2017). The mutant strain ΔBCAS0292 displayed 2.2-fold reduction in attachment to CFBE41o^−^ cells relative to the WT strain (p = 0.0073) (Figure 2a). Adhesion was rescued to WT levels in the Δ0292_0292 complemented strain, suggesting a role for the BCAS0292 protein in host cell attachment (Figure 2a). BCAS0292 is a predicted to localize at the inner membrane (PSort prediction tool: http://psort1.hgc.jp/form.html; certainty 0.117). Therefore, to examine if it was directly involved in host cell attachment, the BCAS0292 gene was cloned into *E. coli* BL21 cells, expression induced with 1 mM IPTG and cellular attachment quantified using confocal immunofluorescent microscopy (Figure 2b). *E. coli* binds poorly to lung epithelial cells with roughly 2 cells attached per 100 epithelial cells (Figure 2c). *E. coli* BL21_BCAS0292 cells showed no increase in attachment to CFBE41o^−^ cells, relative to control BL21 cells, with a mean value of 1.5 bacterial cells per 100 epithelial cells. Expression of BCAS0292 protein under these experimental conditions was confirmed by SDS-PAGE electrophoresis (Figure 2d). The absence of an effect on host cell attachment of *E. coli* expressing BCAS0292 together with the reduction in host cell attachment of the ΔBCAS0292 *B. cenocepacia* mutant indicates that the protein exerts an indirect role in bacterial adhesion to lung epithelial cells.

**Figure 2:**
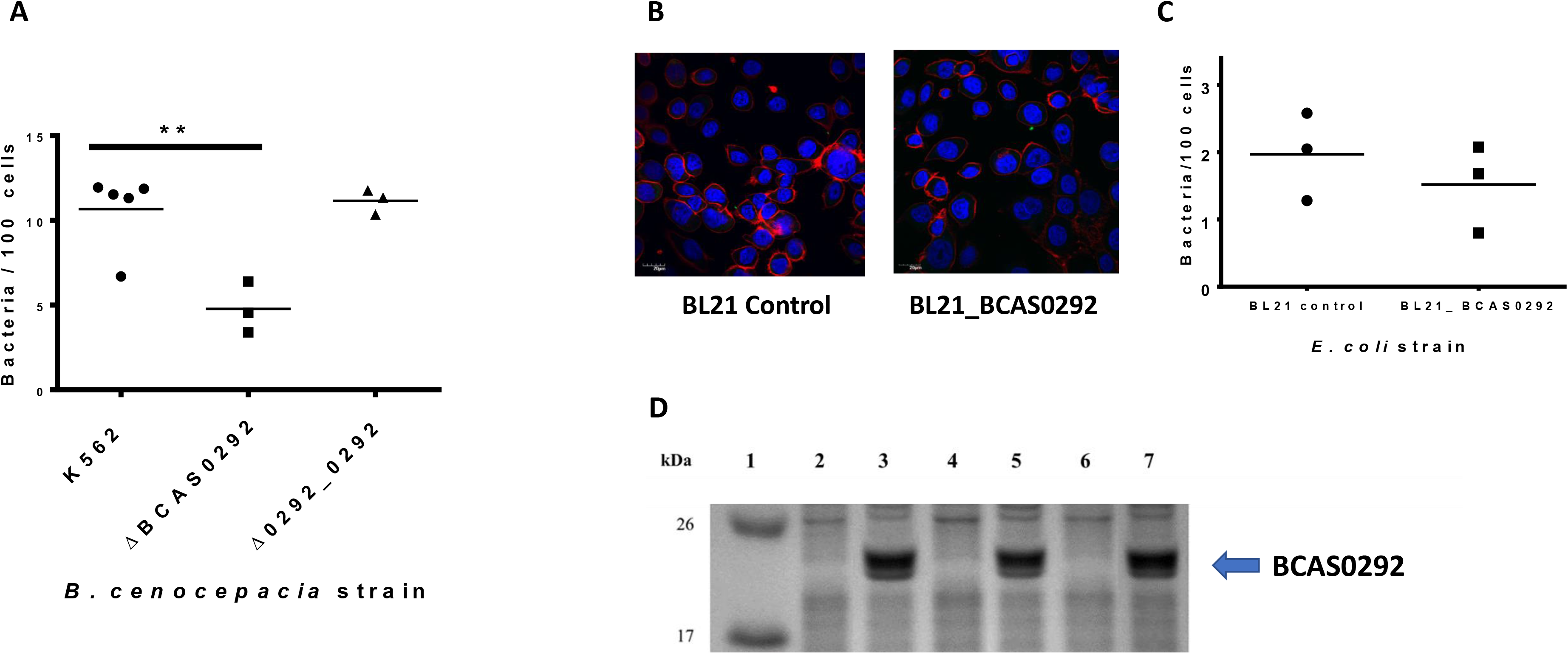
Role of BCAS0292 in epithelial cell attachment. **A)** Attachment of K56-2 WT, ΔBCAS0292 mutant and Δ0292_0292 complemented strain to CFBE41o^−^ cells at an MOI 50:1. Data represents the mean number of bacteria/ 100 cells for each strain with 20 fields of view counted per data point in three independent experiments, ** p =0.0073. B) Confocal microscopy images representing the attachment of *E. coli* BL21 control and *E. coli* BL21_BCAS0292 strains to CFBE41o-cells. Bacteria were labelled using a primary anti-*E. coli* FITC-conjugated antibody (green). Nuclei of CFBE41o-cells were counterstained with DAPI (blue) and actin stained with phalloidin conjugated with Alexa fluor 568. C) Attachment *E. coli* BL21 control and *E. coli* BL21_BCAS0292 to CFBE41o-cells, using an MOI 50:1. Data represents the mean number of bacteria/ 100 cells for each strain, with 20 fields of view counted per data point in three independent experiments. D) Confirmation of expression of BCAS0292, following induction in *E. coli* BL21 cells with 1mM IPTG by Coomassie Blue stained SDS PAGE gel (conditions used for Figure 1C). Lane 1: MW marker; Lane 2: *E. coli* BL21 expression control; Lane 3: *E. coli* BL21_BCAS0292 after incubation with IPTG (1mM) for 2 h.

### Analysis of the susceptibility of K56-2 WT and ΔBCAS0292 to polymyxin B and meropenem

BCAS0292 expression was previously shown to be increased in response to antibiotics but the encoded protein did not have a direct role in resistance (Sass *et al.*, 2011). To further examine antimicrobial susceptibility, the sensitivity of the ΔBCAS0292 mutant to meropenem and polymyxin B (not previously examined) was measured to determine if the absence of this protein affected the susceptibility of the mutant to either of these antibiotics. The ΔBCAS0292 mutant was 8-fold more sensitive to polymyxin B compared to the WT and complemented strains with an MIC of 12 μg/ ml for ΔBCAS0292 compared to 96 μg/ ml for the WT and complemented strains (Figure 3a). Polymyxin B targets LPS components (Mares *et al.*, 2009) and destabilises the outer membrane; suggesting that the loss of BCAS0292 may play a role in the maintenance of membrane integrity. In contrast, ΔBCAS0292 was more resistant to the β-lactam meropenem in comparison to the WT and complemented strains with an MIC of 4 μg/ ml for ΔBCAS0292 compared to 1.5 μg/ ml for the WT and complemented strains (Figure 3b).

**Figure 3:**
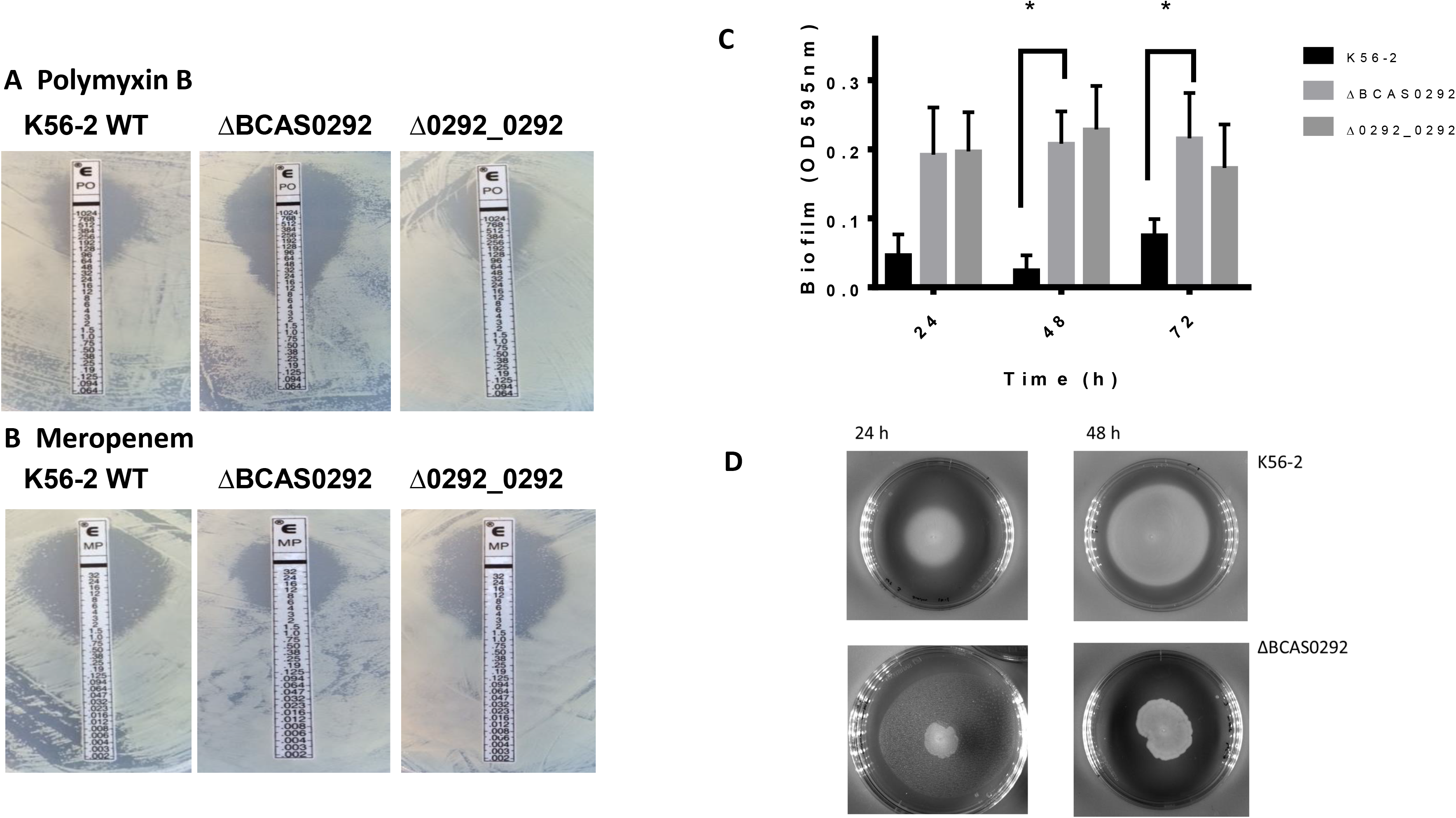
Effect of BCAS0292 on antibiotic sensitivity. Images represent the susceptibility of WT, mutant and complemented strains to polymyxin B (A), (0.064-1,024 μg/ ml) or meropenem (B), (0.002-32 μg/ ml) using E-test strips. The data shown are representative of three independent experiments. (C). Bar chart representing biofilm formation of K56-2 WT, ΔBCAS0292 and Δ0292_0292 after 24, 48 and 72 h. Error bars represent the standard deviation from three independent experiments. *p < 0.0001 represents a significant difference between ΔBCAS0292 and K56-2 WT strain. D) Swimming motility of K56-2 WT and ΔBCAS0292 mutant cells 24 and 48 h after inoculation of agar plates. Images are representative of three independent experiments.

### Effect on Biofilm formation and motility

Biofilm formation is regulated by the cepIR quorum sensing system, which has been shown to regulate BCAS0292 expression (O’Grady and Sokol, 2011). Therefore, we examined whether deletion of the BCAS0292 gene had any effect on biofilm formation. ΔBCAS0292 mutant cells showed significant increased biofilm formation (p < 0.0001) at all time points, which was partially compensated in the complemented strain at 72 h (Figure 3c), suggesting that the BCAS0292 protein repressed biofilm formation. Motility was also impaired in the mutant strain (Figure 3d). The K56-2 WT showed a mean swimming diameter of 3.55 ± 0.12 cm and 6.22 ± 0.12 cm after 24 and 48 h, respectively. Whereas the ΔBCAS0292 mutant strain showed average swimming diameters of 1.83 ± 0.22 and 2.72 ± 0.23 cm after 24 and 48 h, respectively. This phenotype was not restored in the complement strain.

### Whole proteome analysis of WT and ΔBCAS0292 strains

The lack of a direct effect on host cell attachment, combined with the apparent repression of biofilm formation; behaviour in response to antibiotics; and impact on motility suggested that the BCAS0292 protein has a regulatory effect. To test this hypothesis, we compared the proteome profiles of the ΔBCAS0292 mutant with WT under stationary phase conditions. Stationary-phase cultures were used for this experiments since BCAS0292 expression is maximal in stationary phase (Sass *et al.*, 2013). We identified 1,933 proteins showing different abundance between the strains, of which 23 proteins were unique to the ΔBCAS0292 mutant while 46 were unique to K56-2 WT cells (Table 2). Unique proteins are those which are detected in all replicates of one strain but are undetectable in all four replicates of the comparator. A volcano plot highlights that there were more proteins showing reduced abundance and that the fold-changes were more extensive than those showing increased abundance (Figure 4a). We identified 391 proteins with significantly increased abundance by ≥ 1.5-fold in ΔBCAS0292 while 556 proteins (59 % of the total altered proteins) showed significantly reduced abundance by ≥ 1.5-fold. These results suggest BCAS0292 is a global positive regulator of protein expression (Figure 4a). Among the proteins showing reduced abundance in the ΔBCAS0292 mutant were proteins involved in metabolism (36%), virulence (7%), stress responses (6%), and uncharacterized proteins (35%) (Figure 4b). The fold reduction in protein abundance was substantial: 92 proteins showed reduced abundance (ranging from 3- to 276-fold) in the mutant, and 272 proteins were reduced by over 2-fold (Table 3). In contrast, among the 391 proteins with significantly increased abundance by ≥ 1.5-fold in the ΔBCAS0292 mutant relative to the K56-2 WT strain, 58 were increased by ≥ 3.0 fold, with the greatest fold increase in abundance being only 9.8-fold (Supplemental Table S3). When proteins showing increased abundance of 3 to 9.8-fold were grouped according to their function (GO annotation) 67% had roles in metabolic processes, 19% in translation and 7% had roles in the stress response (Supplementary Table S3).

**Figure 4:**
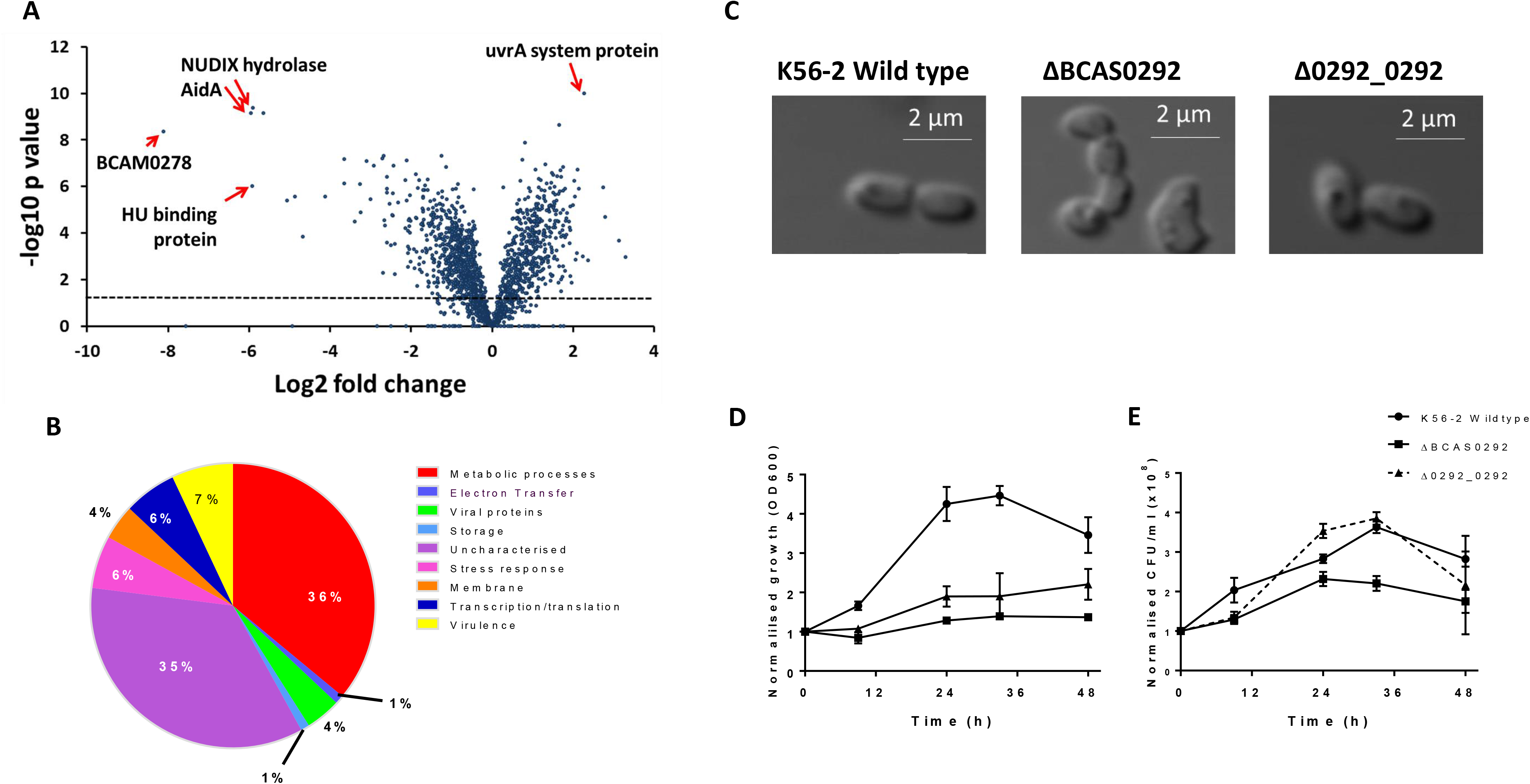
Effect of deletion of the BCAS0292 gene on the proteome and phenotype of *B. cenocepacia* K56-2. A) Fold change in protein abundance plotted as a volcano chart of log2 fold change in protein abundance in the ΔBCAS0292 mutant strain relative to WT strain. Points on the negative side of the X axis represent a reduction in protein abundance. The proteins with the greatest changes in abundance are highlighted with red arrows. Dashed line represents the 1.5-fold change in abundance. B) Pie chart representing the functional GO categories of the proteins that showed abundance reduced by ≥ 1.5 fold in the ΔBCAS0292 mutant strain relative to the K56-2 WT strain. C) Effect of deletion of BCAS0292 on cell morphology as determined by phase contrast microscopy. D) Growth and survival of K56-2 strain, ΔBCAS0292 mutant and 0292_0292 complement strain in 6% oxygen. Cells were cultured in a hypoxia chamber at 6% oxygen for up to 48 h. Growth was determined by OD600nm. Survival was determined by serially diluting aliquots of cultures that had been incubated at 6% oxygen in a controlled hypoxia chamber, plating at time intervals and culturing for 48h in a normoxic environment.

**Table 1:** Total number of differentially expressed proteins between K56-2 WT and Δ*BCAS0292* strains.

| Protein groups | Number of differentially expressed proteins |
| --- | --- |
| Total number of proteins identified | 1,933 |
| Total number of proteins altered by $\geq 1.5$ or absent in all replicates of a either strain | 1,016 |
| Proteins unique to K56-2 WT | 46 |
| Proteins unique to $\Delta BCAS0292$ | 23 |
| Proteins significantly downregulated (by $\geq 1.5$-fold) in the $\Delta BCAS0292$ strain | 556 |
| Proteins significantly upregulated (by $\geq 1.5$-fold) in the $\Delta BCAS0292$ strain | 391 |

**Table 2:**
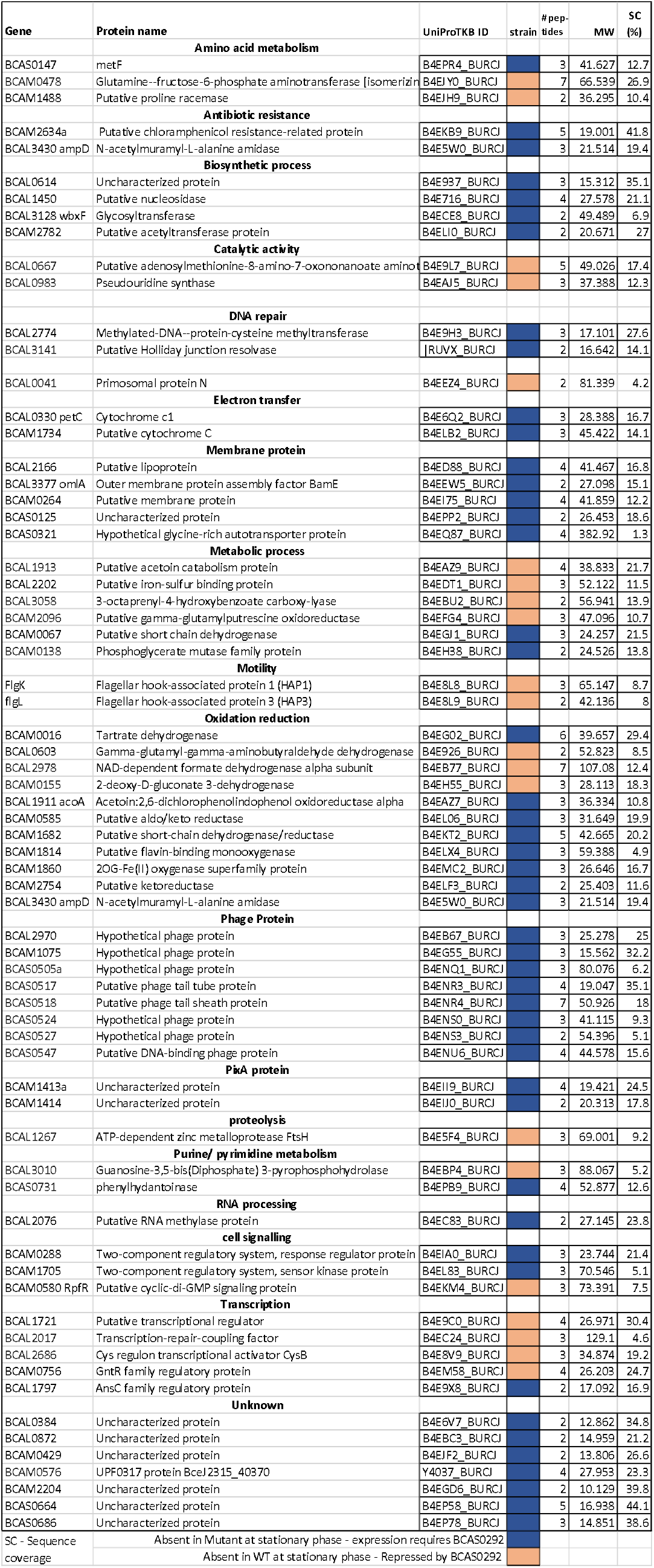
Proteins that were unique to either the ΔBCAS0292 or K56-2 WT strain, i.e. proteins were absent or undetectable in the mutant strain relative to WT or in the WT strain relative to the mutant strain.

**Table 3:**
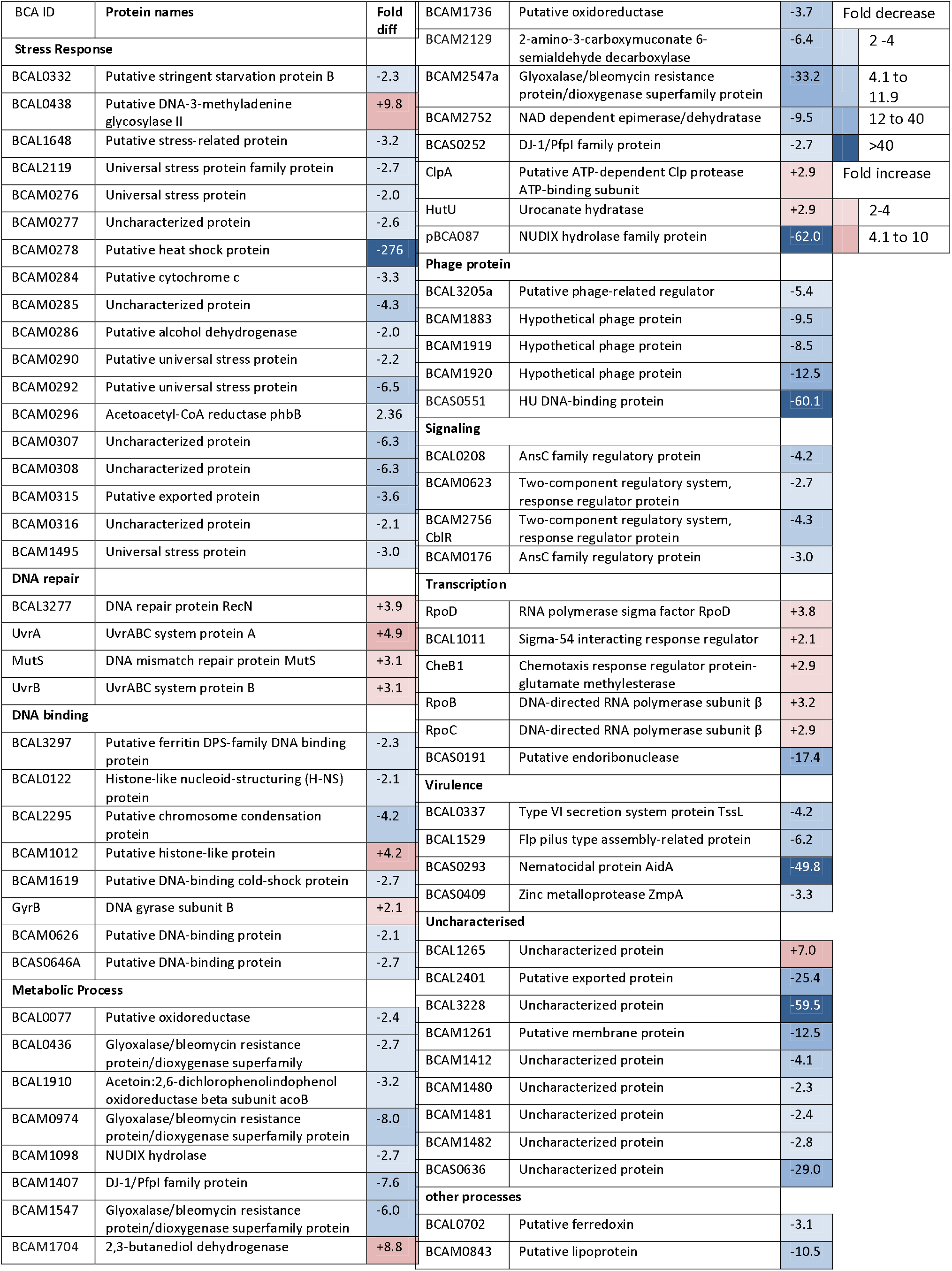
Examples of proteins altered by >2 fold ΔBCAS0292 mutant relative to K56-2 WT cells.

#### Proteins unique to either strain

Proteins that are unique to the ΔBCAS0292 mutant proteome are likely to be proteins that are repressed by BCAS0292. These proteins include RpfR, a receptor for the autoinducer BDSF and two flagellar hook-associated proteins FlgK and FlgL (Table 2). The majority of proteins that are absent in the mutant (and consequently may rely on the presence of BCAS0292 for expression) are proteins with catalytic activity (67%). In addition, 8 (17%) are classified as phage proteins. There are 10 proteins classified as oxidation/reduction proteins, of which 7 are unique to the WT strain. Four out of five transcriptional regulation proteins identified were unique to the ΔBCAS0292 mutant proteome, while the other was unique to the WT strain proteome.

#### Proteins involved in metabolic functions

Proteins with roles in various metabolic processes represented 36 % of the total number of proteins reduced in abundance (Figure 4b). Among these proteins are a plasmid encoded NUDIX hydrolase family protein (pBCA087) reduced by 62-fold (Table 3). This was one of five NUDIX proteins identified that showed reduced abundance in the mutant (Supplemental Table S3). In addition two Glyoxalase/bleomycin resistance protein/dioxygenase superfamily proteins (BCAM2547a, BCAM0974) were reduced by 33.2-fold and 8-fold in ΔBCAS0292, and 6 other proteins in this superfamily were also reduced by 6 fold to 1.74-fold (Table 3 and Supplementary Table S2). The majority of proteins unique to the WT strain also had functions in metabolic processes (37%) (Table 2). Most of these proteins function as enzymes which act as catalases, hydrolases, dehydrogenases, oxidoreductases or transferases, indicating that there is a global change in metabolism of these cells in the absence or presence of BCAS0292. Among the proteins involved in metabolic processes that showed increased abundance were 111 proteins with amino acid metabolism functions (i.e. 25% of total proteins with increased abundance, in comparison with only 42 (6.4%) of proteins with this function showing reduced abundance (Table 3). Other proteins upregulated in the metabolic processes category had synthase, transferase and cyclohydrolase activities, among others, highlighting the broad range of functions carried out by proteins in this category.

#### Proteins involved in the adaptive response

The abundance of many proteins with roles in the stress response were reduced in the ΔBCAS0292 mutant, representing 6 % of the total number of downregulated proteins in this group (Figure 4b). Of particular interest was the reduced abundance of 19 proteins encoded on the low-oxygen activated (*lxa*) locus (BCAM0272 to 0323) in the mutant strain, highlighting the possible co-regulation of expression of proteins encoded on this gene cluster (Table 3 and Supplementary Table S2). This locus has been shown to be upregulated under hypoxic conditions and in stationary phase (Sass *et al.*, 2013); and in sequential clinical *B. cenocepacia* isolates over time of chronic infection (Cullen *et al.*, 2018). In particular a two-component system response regulator encoded within the locus, BCAM0288, was undetectable in the mutant strain. Eight *lxa*-encoded proteins that showed reduced abundance proteins have stress response functions. The abundance of a small heat shock/α-crystallin protein (BCAM0278) was dramatically reduced (276-fold) relative to K56-2 WT. There are 6 universal stress proteins (USPs) encoded on the *lxa* locus, and five of these showed reduced abundance in the ΔBCAS0292 mutant strain, with BCAM0292, being the lowest (6.5-fold). Other stress response proteins in this category were reduced in abundance from 1.5- to 2.2-fold. Three other *lxa* locus encoded proteins reduced by 2- to 6.3-fold have roles in phospholipid binding, electron transfer, storage polymer turnover and alcohol turnover. Sass *et al.* (2013) identified four other low-oxygen co-regulated gene loci and five proteins that are encoded by *lxa* co-regulated genes and these were also reduced in abundance: BCAM1480- to- 1482, another USP, BCAM1495, and an uncharacterised protein, BCAM1496. In contrast, only three *lxa*-encoded proteins BCAM0299 (a zinc-binding alcohol dehydrogenase), BCAM0300 (Metallo-β-lactamase superfamily protein) and BCAM0310 (Putative ribonucleotide reductase), were increased 1.9- to 1.6-fold. Further, there were 27 proteins with roles in stress responses showing increased abundance and all but one of these proteins were associated with DNA repair functions (Table 3).

#### Phage or viral proteins

Viral proteins represented 17 % of the total number of proteins that were undetectable in the ΔBCAS0292 mutant, including several hypothetical phage proteins, a putative phage tail tube protein (BCAS0517), a putative phage tail sheath protein (BCAS0518) and a putative DNA-binding phage protein (BCAS0547). Viral proteins also represented 4 % of the proteins reduced in abundance (Figure 4b), including hypothetical phage proteins (BCAM1883, BCAM1919 and BCAM1920) and a putative phage-related regulator (BCAL3205a), which were reduced by 9.5-, 8.5-, 12.5- and 5.4-fold, respectively (Table 3).

#### Virulence proteins

Three T6SS secretion system proteins encoded by BCAL0337, BCAL0341 and BCAL0343 (TssD, Hcp1) were also reduced in abundance by 4.2 to 1.6-fold in the mutant. Other virulence proteins included a zinc metalloprotease (ZmpA) (BCAS0409), a known quorum sensing regulated virulence factor (Kooi *et al.*, 2006); nemtatocidal AidA and a Flp pilus type assembly-related protein (BCAL1529), were also reduced. Flagellar hook-associated protein 1 (HAP1) and flagellar hook-associated protein 3 (HAP3) were both identified in the ΔBCAS0292 mutant while absent from the K56-2 WT strain, indicating that BCAS0292 represses their expression. In contrast, two proteins belonging to the T3SS and T6SS families of proteins, were increased in abundance in the ΔBCAS0292 strain, including a type III secretion system protein (BCAM2040) and a putative type VI secretion system protein TssK (BCAL0338) (Supplementary Table S3).

#### Uncharacterised or hypothetical proteins

BCAS0292 is identified as a hypothetical protein (Winsor *et al.*, 2008) and over a third (35%) of the proteins showing reduced abundance in the ΔBCAS0292 mutant were also classed as uncharacterized proteins or hypothetical proteins, including BCAL3228 and BCAS0636 (59.5- and 29-fold reduced abundance respectively). A putative exported protein (BCAL2401), with unknown function, was also reduced by 25.4-fold in the ΔBCAS0292 mutant strain. There are five PixA containing proteins in the sequenced J2315 genome (Winsor *et al.*, 2008) and all other PixA proteins showed reduced abundance (between 4.1 to 49.8-fold) or were undetectable in the ΔBCAS0292 mutant suggesting a co-regulation or co-dependence of these uncharacterised proteins. Three of these PixA proteins are encoded on chromosome 2, with BCAM1412, being downregulated by 4.1-fold while the proteins encoded on the adjacent genes, BCAM1413 and BCAM1414 were both absent in the ΔBCAS0292 mutant. BCAM1412 and BCAM1414 genes are both orthologs of *aidA*, while BCAM1413 is not.

#### Transcription and Translation

Proteins with roles in transcription represented another 6 % of the proteins reduced in abundance (55 proteins), including a HU DNA-binding protein (BCAS0551) which was reduced by a substantial 60.1-fold in the mutant. Among the 23 proteins unique to the ΔBCAS0292 mutant strain were 4 proteins (19%) with roles in transcriptional regulation and a transcription-repair-coupling factor (BCAL2017). There were only 17 proteins with roles in transcription that were increased in abundance in the mutant strain, including two DNA-directed RNA polymerase subunit proteins (RpoB and RpoC) which were increased by 3.2- and 2.9-fold. A large number of ribosomal proteins were also increased in the ΔBCAS0292 mutant (Supplemental Table S3).

#### Proteins associated with other functions

Membrane proteins with roles in adhesion and possible roles in structure/ transport represented 4 % of total number of proteins reduced in abundance in the mutant strain (Figure 4b) including ten membrane proteins, along with other surface proteins were lost or significantly reduced (by ≥ 1.5-fold) in the ΔBCAS0292 strain (Table 3). Among these were proteins associated with roles in adhesion that were reduced by between 3.5- and 12.5-fold in the ΔBCAS0292 mutant including a maltose-binding protein (BCAL3041), putative lipoproteins (BCAM0834 and BCAM0944) and a putative membrane protein (BCAM1261. In addition, four membrane proteins were undetectable in the mutant strain, including the outer membrane assembly protein, BamE and a putative lipoprotein (BCAL2166), a putative lipoprotein (BCAL2166) and a putative membrane protein (BCAM0264) (Table 2).

### ΔBCAS0292 mutant shows gross morphological changes

Given the vast number of proteins with altered abundance in the mutant strain we compared the morphology of the mutant with the WT and complement strains. Unsurprisingly, the ΔBCAS0292 mutant cells had a grossly altered morphology, with a highly irregular cell surface and complete loss of their rod-like shape (Figure 4c). This was reversed in the complemented strain suggesting that BCAS0292 is involved in maintaining cell structure integrity which may contribute to the reduced adherence of this strain to lung epithelial cells.

### ΔBCAS0292 mutant has limited growth at 6% oxygen

Given that several proteins encoded on the *lxa* locus showed reduced abundance in the ΔBCAS0292 mutant, we compared the growth of WT and ΔBCAS0292 mutant in 6% oxygen in a controlled hypoxia chamber. The growth of the mutant strain was severely reduced under hypoxic conditions (*p* < 0.0001, Figure 4D). When the bacteria from each strain were plated at time intervals and CFU/ml compared, it was apparent that the survival of the mutant strain was also impaired (p = 0.0294) following liquid culture under hypoxic conditions. In contrast, survival recovered to WT levels in the complemented strain.

### Structural analysis of BCAS0292 protein

In order to help elucidate the function of BCAS0292, we analysed its structural features in solution. Following affinity chromatography purification, His-tagged BCAS0292 was purified by SEC to remove any contaminating proteins and protein aggregates and obtain pure fractions (Figure 5a). The secondary structure of rBCAS0292, purified under non-denaturing conditions was determined by Circular Dichroism (CD). A peak minimum was observed at 215 nm indicating that rBCAS0292 has a β-sheet secondary structure (fig. 5b). The purified protein was highly stable with a melting temperature (Tm) of 49 °C, (Figure 5c). Light scattering was used to determine the oligomeric state of the BCAS0292 protein. Rayleigh scattering showed that it has a dimeric organisation, in both reducing and oxidising conditions, with a molecular weight of 39.4 ±0.5 kDa (Figure 5d). Despite several attempts, we were unable to obtain crystals of BCAS0292, therefore molecular modelling was used to predict the 3D structure of the protein using consensus sequence alignment with PixA inclusion body protein BCAM1413 from *B. cenocepacia* described by Nocek et al., (2013) as a template (PDB code 4lzk, E-value- 3.5e^−55^). Consistent with CD analysis (Figure 5b), BCAS0292 homology model presents a structure with mostly β-folds, and only one small α helix in the N-terminal region of the protein (residues 13-21) (Figure 6a). Also, as experimentally proven by light scattering studies (Figure 5d), the molecule has a dimeric organization, where two jelly-roll like monomers are compactly held together (Figure 6a). Electrostatic potential calculations show that the entire surface of the protein has a strong negative charge distribution (Figure 6b), a finding which excludes BCAS0292 regulatory function occurring via interactions with DNA or RNA. Instead, similar electrostatic surface distributions are a distinctive feature of DNA mimicry (Wang *et al.*, 2014).

**Figure 5:**
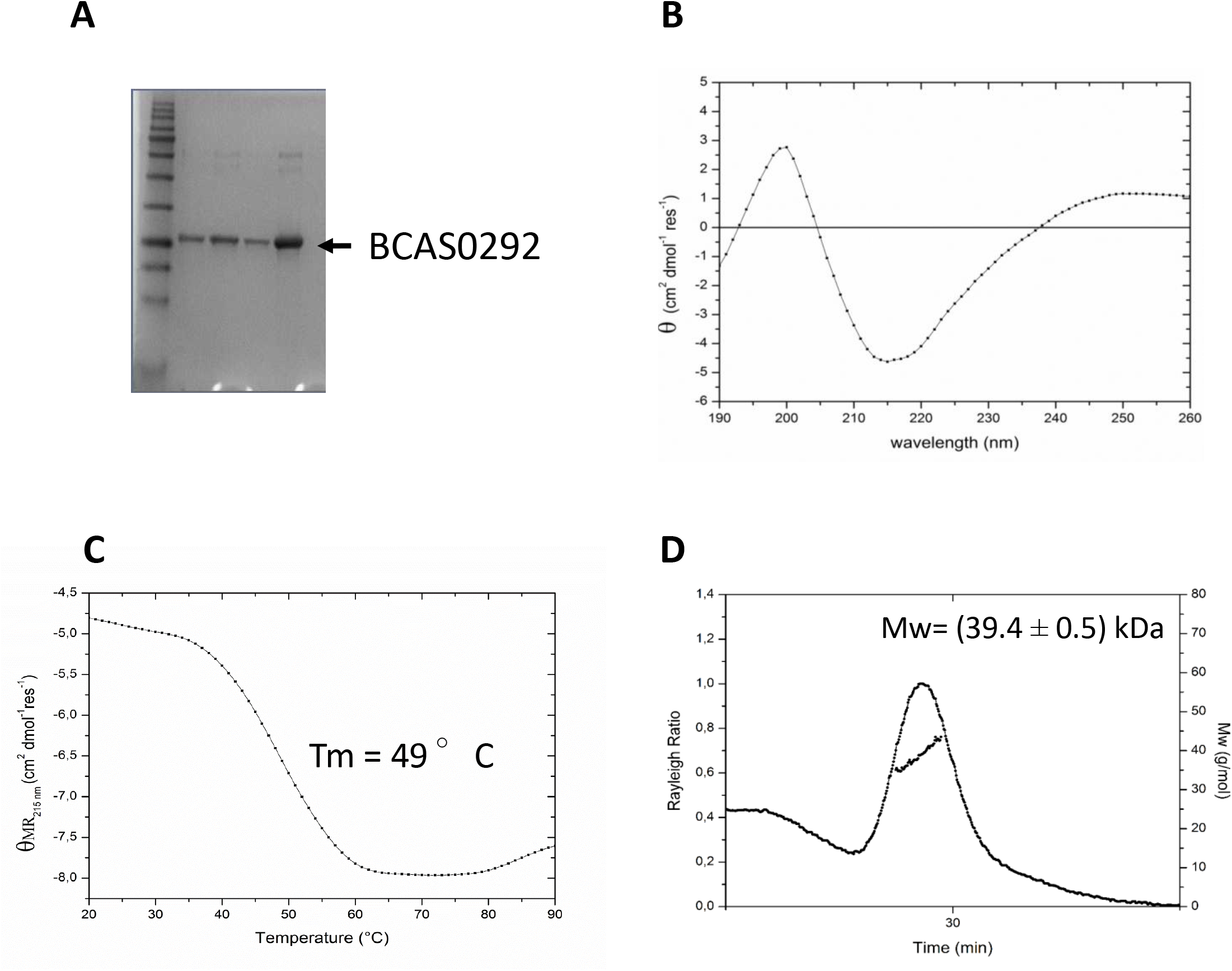
Structural analysis of rBCAS0292. A) Size-exclusion chromatogram and subsequent SDS-PAGE analysis of purified rBCAS0292 protein. B) CD analysis of purified rBCAS0292 protein to determine the secondary structure at 20 °C. A minimum was observed at 215nm, indicative of a β-sheet fold. C) Thermal gradient to determine the Tm of purified rBCAS0292, demonstrating a Tm of 49 °C. D) CD analysis of purified rBCAS0292 protein to examine the ability of the protein to re-fold. Black line: 20 °C, green line: 90 °C, red line: 20 °C following thermal gradient. This indicates that the protein aggregates following heat denaturation and is unable to restore the correct fold at 20°C. E) Light scattering analysis to determine the oligomeric state of rBCAS0292, which indicated that the protein has a dimeric organisation.

**Figure 6:**
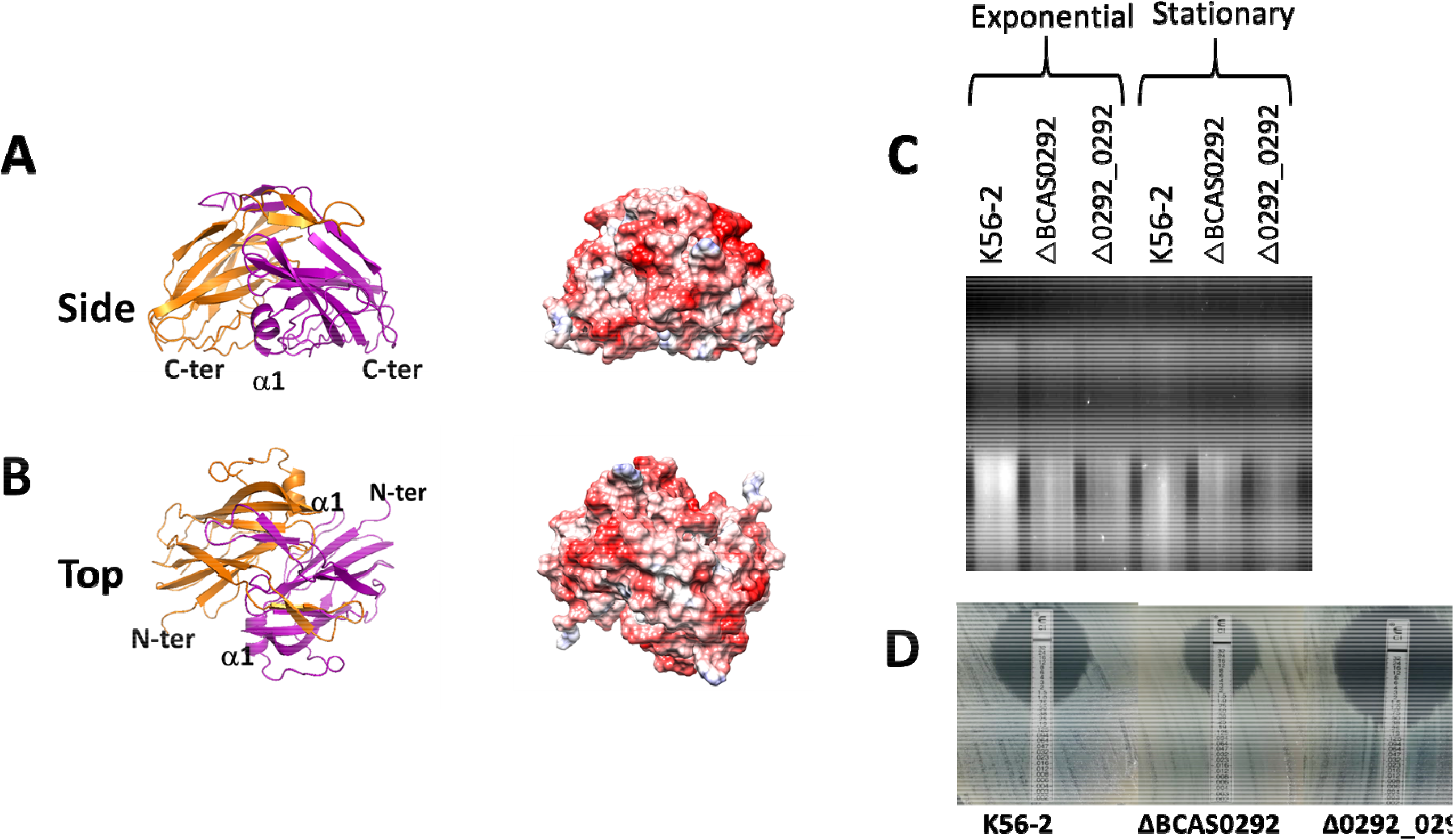
Molecular modelling of BCAS0292 and impact on DNA supercoiling A) Cartoon representation of the predicted 3D structure of the dimer. Consensus based sequence alignment was used to determine the predicted structure using MODELLER software. **B)** Analysis of the electrostatic potential of the hypothetical protein. Side view of the predicted dimer model. C) Comparison of DNA supercoiling of plasmid DNA extracted from *B. cenocepacia* K56-2, ΔBCAS0292 and Δ0292_0292 cells cultured to either exponential or stationary phase. Chloroquine-stained gel image is representative of three independent experiments. D) Ciprofloxacin susceptibility of K56-2, ΔBCAS0292 and Δ0292_0292 strains as determined by E-strips. Images representative of three independent experiments.

### BCAS0292 may function as a DNA mimic protein

Given that BCAS0292 had an overall negative surface charge, we hypothesized that it bound to DNA-binding proteins to mediate its global regulatory activity. To corroborate this hypothesis we examined whether BCAS0292 alters DNA supercoiling, an activity associated with global changes in bacterial gene regulation (Shortt *et al.*, 2016, Scanlan *et al.*, 2017) by investigating the impact of the presence of BCAS0292 on DNA topology. BCAS0292 has been shown to be upregulated in stationary phase (Sass *et al.*, 2013), therefore we compared stationary phase cultures with exponential phase cultures for the WT K56-2, ΔBCAS0292 mutant and complemented strain. DNA topoisomers were not as highly resolved in *B. cenocepacia* (compared with *E. coli*, Supplemental Figure S1) presumably due to the very high levels of restriction modification systems expressed in *Burkholderia*. There were no obvious differences in supercoiling in plasmid DNA extracted from exponential phase cultures of the three strains (Figure 6c). In contrast, DNA extracted from stationary phase cultures of the mutant showed a more relaxed DNA topology with fewer highly supercoiled topoisomers at the top of the chloroquine gel relative to WT and complement strains (Figure 6c), indicating that BCAS0292 expressed in stationary phase alters DNA supercoiling. Fluoroquinolones target DNA gyrase and topisomerase IV, enzymes involved in establishing and or relaxing supercoiling. Consequently, any difference in DNA topology may have impact on the susceptibility to fluoroquinolones between the mutant and WT. In addition, the mutant clearly showed a 2-fold decrease in ciprofloxacin susceptibility, which was reversed in the complemented strain (Figure 6d), further indicating that a lack of BCAS0292 expression impacted on DNA topology.

## Discussion

The adaptation of opportunistic pathogens enhancing their fitness in the host and consequently their successful colonisation requires substantial rearrangement of gene expression. Understanding the processes of bacterial regulation in pathogens that adapt to chronic infection is crucial, particularly those that are virtually impossible to eradicate. Our initial studies on host cell attachment of the *B. cenocepacia* K56-2 ΔBCAS0292 mutant strongly pointed towards a regulatory role for the BCAS0292 protein. The mutant showed reduced attachment to epithelial cells relative to the K56-2 WT strain but induction of BCAS0292 expression had no impact on the adhesion of *E. coli* BL21 cells to CFBE41o^−^ cells in contrast to our earlier work on peptidoglycan-associated lipoprotein (Dennehy *et al.*, 2017). Consistent with this, the dramatic upregulation of BCAS0292 and *aidA* genes in the presence of low oxygen or in stationary phase *B. cenocepacia* cultures (Sass *et al.*, 2013), coupled with the dramatic downregulation of these genes in the quorum sensing mutant strain K56-2 Δ*cepR* (O’Grady and Sokol, 2011) demonstrated that both genes are highly responsive and collectively suggest that BCAS0292 plays a role in the response of *B. cenocepacia* to extracellular stress. Our work now demonstrates that it has a global regulatory role in the expression of over a thousand proteins including those involved in stress responses and metabolism. Specifically, the reduced abundance of five of the six USPs and 14 other *lxa*-encoded proteins in the mutant supports the role for BCAS0292 in the regulation of the hypoxic stress responses. Further, the impaired growth of the mutant when cultured in a tightly controlled environment of 6% oxygen also substantiates a role for BCAS0292 in the adaptation of *B. cenocepacia* to hypoxic conditions.

BCAS0292 may also play an indirect role in antibiotic susceptibility. The increased susceptibility of the ΔBCAS0292 mutant to polymyxin B, an antibiotic that targets LPS was not surprising given the impact that the absence of BCAS0292 had on the abundance over 10 membrane proteins and on cell morphology. Although meropenem is quite resistant to degradation by β-lactamases, the production of metallo-β-lactamases can facilitate meropenem resistance (Yano *et al.*, 2001); thus the increased abundance of metallo-β-lactamase (BCAM0300) likely contributed to meropenem resistance in the mutant. Similarly, the reduced virulence in the *G. mellonella* infection model of the mutant is likely due to the depressed expression of some of the virulence factors in the mutant, including ZmpA; T6SS proteins, and a flp pilus protein.

Comparison of the proteomes of the K56-2 WT and ΔBCAS0292 mutant strains highlight that discrete groups of proteins involved in other specific processes, were also affected by the deletion of BCAS0292. A recent *in silico* analysis demonstrated that *B. cenocepacia* genomes are rich in phage genomes, with 63 prophage regions identified in 16 sequenced *B. cenocepacia* databases (Roszniowski *et al.*, 2018). A total of 18 viral proteins, most of which are encoded within two clusters on chromosomes 2 and 3, were either absent or dramatically reduced in the ΔBCAS0292 mutant, indicating that there is a positive correlation between the abundance of specific viral proteins and BCAS0292 expression. The finding that eight phage proteins were among the 49 proteins (16%) that were undetectable in the mutant strengthens this view. Among these 18 phage proteins are six encoded by the intact prophage Bcep-Mu which is widespread among human pathogenic *B. cenocepacia* strains of the ET12 lineage (Summer *et al.*, 2004) and four are encoded in a newly identified prophage region (BCAM1879 to BCAM1926), not previously reported (Roszniowski *et al.*, 2018). Finally, BCAL3205 is a helix-turn-helix transcriptional regulator associated with the control of lytic/lysogenic cycle of phages and bacterial gene expression (Aravind *et al.*, 2005). Its reduced abundance in the mutant may regulate the expression of the other phage proteins downstream. The extracellular portion of the T6SS and bacteriophage tail-associated protein complexes are both structurally and functionally similar (Zoued *et al.*, 2017). Thus, the shared impact of BCAS0292 gene deletion implicates BCAS0292 protein in the regulation of expression of these phage tail-like proteins and T6SS in *B. cenocepacia* by a common mechanism.

Among the proteins that were unique to the mutant (i.e. undetectable in the WT) at stationary phase was RpfR, the receptor for Burkholderia diffusible signal factor (BDSF). RpfR is involved in the synthesis and degradation of cyclic-di-GMP, a second messenger which regulates biofilm formation, biofilm dispersal, motility, extracellular protease production, antibiotic resistance and other virulence associated phenotypes. Point mutations in different *rpfR* domains resulted in mutants showing increased biofilm formation and impaired motility, consistent with the ΔBCAS0292 mutant (Mhatre *et al.*, 2020). Given that the ΔBCAS0292 mutant showed increased biofilm formation and reduced motility and uniquely expressed RpfR, it is likely that the BCAS0292 protein mediates some of its pleiotropic effects via RpfR suppression.

Our observation that all 5 PixA proteins encoded in *B. cenocepacia* were either reduced by up to 50-fold or absent in the ΔBCAS0292 mutant, indicates that the other PixA proteins require BCAS0292 for their complete expression. A reduction in AidA abundance in the mutant was expected, as the *aidA* gene is located in the same operon and co-regulated with BCAS0292 (O’Grady *et al.*, 2009); however, the three other pixA genes are located on chromosome 2 suggesting their co-regulation with BCAS0292. Studies on PixA and PixB in *X. nematophila* revealed they had no direct role in virulence or colonisation of nematodes (Lucas *et al.*, 2018); however, they may also be involved in regulation of the response to different environmental conditions in this bacterium.

Purified rBCAS0292 protein is dimeric, consistent with many functionally important proteins involved in regulation such as transcription factors (Zhanhua *et al.*, 2005, Ispolatov *et al.*, 2005), which also points towards a regulatory role for the BCAS0292 protein. The unexpected finding that BCAS0292 has a highly negatively charged surface, indicated that BCAS0292 is likely to bind to DNA or RNA binding protein(s) and may act as a DNA mimic protein, which led us to evaluate its impact on DNA topology in stationary phase. DNA supercoiling is considered a crude regulator of gene expression with variable responses to environmental signals and the potential to act right across the bacterial genome (Dorman and Corcoran, 2009). The role of DNA supercoiling in global regulation of SPI1 and SPI2 pathogenicity islands genes in *Salmonella enterica* serovar typhimurium are well described, with negative supercoiling up-regulating SPI1 genes and conferring a more invasive phenotype, while SPI2 genes upregulated by DNA relaxation are essential for survival inside the macrophage (Dorman and Corcoran, 2009, O’ Croinin *et al.*, 2006). Hence DNA supercoiling discriminated between two different intracellular environments and modulated the expression of virulence genes in macrophage cells (O’ Croinin *et al.*, 2006). Alterations in *C. jejuni* supercoiling also resulted in increased cellular invasion and secretion (Scanlan *et al.*, 2017). Differences in DNA supercoiling between the ΔBCAS0292 mutant and WT at stationary phase combined with enhanced ciprofloxacin susceptibility reinforced the view that BCAS0292 can alter DNA topology, suggesting this is the mechanism by which it wields its regulatory effect.

DNA mimic proteins generally have a molecular weight <25kDa and pI values <5.0 and are characterised by having a negative charged distribution on the protein surface, allowing them to mimic the negatively charged phosphate backbone of DNA (Wang *et al.*, 2018) and BCAS0292 matches each of these criteria. Very few (~20) have been discovered to date because, consistent with BCAS0292, they have unique amino acid sequences and structures and are hard to predict using bioinformatic tools. Half of the DNA mimic proteins already identified are bacterial and are involved in many internal response controls (Wang *et al.*, 2014). Two were identified in *Neisseria* spp, namely DMP12 and DMP19 (Wang *et al.*, 2012, Wang *et al.*, 2013). The latter, DMP19, forms a dimer and has a variety of potential binding partners and is suggested to prevent the interaction of Neisseria hypothetical transcription factor from binding to its DNA binding sites (Wang *et al.*, 2012, Huang *et al.*, 2017), while DMP12 is likely to maintain the stability of unbound HU-binding regions (Wang *et al.*, 2014, Wang *et al.*, 2013). Another DNA mimic, MfpA, was identified in fluoroquinolone resistant strains of *Mycobacterium tuberculosis* and functions by protecting the DNA binding site of gyrase from fluoroquinolones (Hegde *et al.*, 2005). Ongoing pull-down analyses involving different approaches to identify possible binding partners of BCAS0292 have been inconclusive to date.

A large number of proteins encoded on gene clusters associated with roles in translation showed increased abundance in the ΔBCAS0292 mutant suggesting that BCAS0292 negatively regulates or represses their expression. Taken together, this led us to propose that BCAS0292, which is dramatically upregulated in response to hypoxic stress and in stationary phase interacts with DNA-binding proteins, altering DNA topology which globally influences transcription and expression of a vast number of genes involved virulence, host cell attachment, antibiotic susceptibility and biofilm formation (Figure 7). Recently, we showed that 20 proteins encoded on the *lxa* locus show increased abundance in late sequential chronic CF isolates relative to the earlier isolates (Cullen *et al.*, 2018). Thirteen of these proteins are also reduced in the ΔBCAS0292 mutant suggesting a common adaptive stress response in the hypoxic CF lung (Supplementary table S4). In contrast, only two proteins encoded on genes within the adjacent 37-gene cci pathogenicity island were significantly reduced in the ΔBCAS0292 mutant indicating that the downregulation of *lxa-*encoded genes in the mutant is specific to the BCAS0292 protein. Subsequent analysis revealed that 109 other proteins that were either absent or reduced in abundance in the mutant strain were also elevated in the late chronic infection *B. cenocepacia* isolates (Suppl Table S5). This represents 64% of the proteins with increased abundance over time of chronic infection (Cullen *et al.*, 2018) and points to a role of BCAS0292 in adaptation to chronic infection. Taken together these data suggest that this protein may contribute to the adaptation and survival of *B. cencocepacia* in the hypoxic environment of the CF lung.

**Figure 7:**
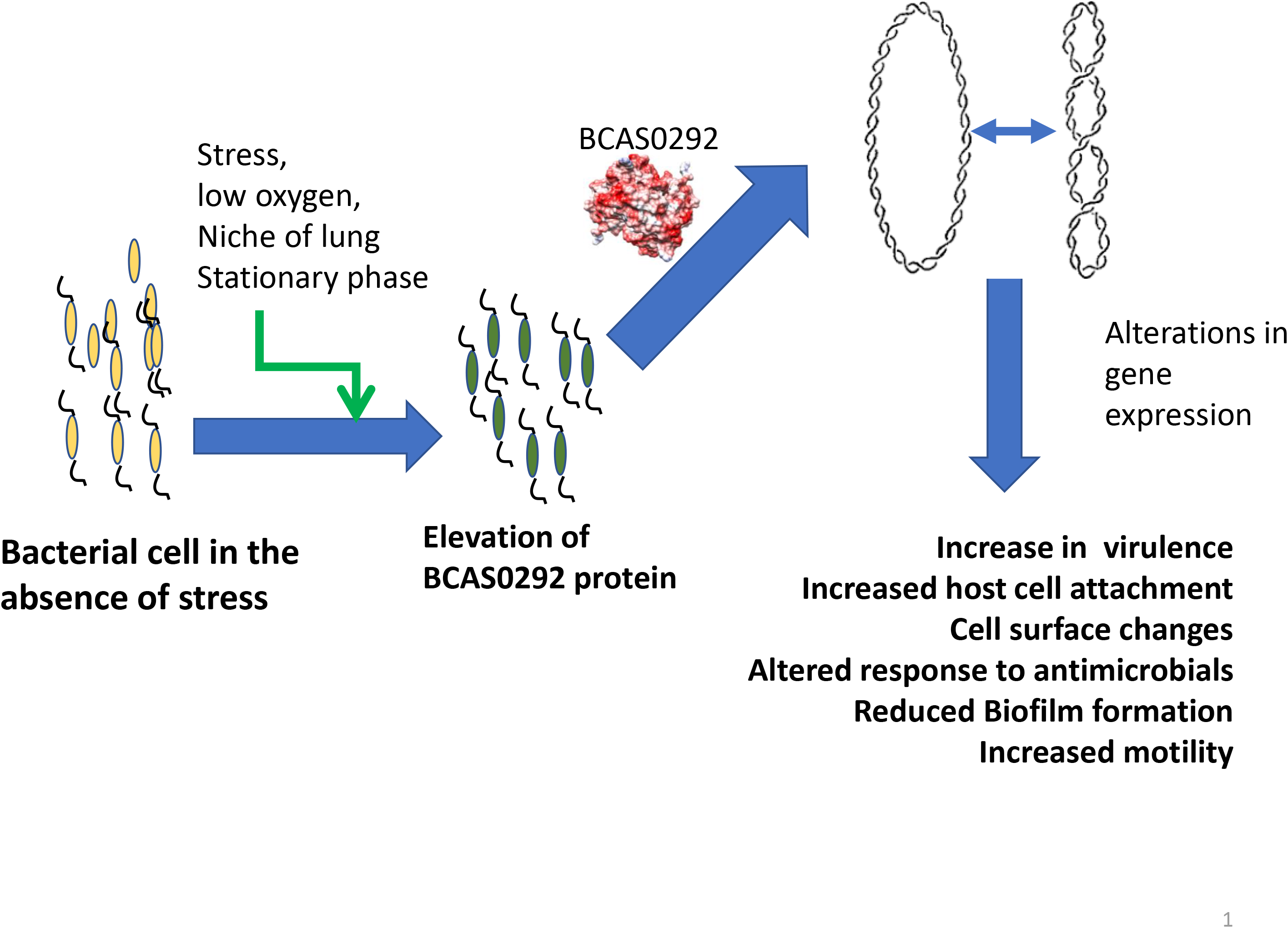
Schematic of the proposed mechanism of regulation by BCAS0292 in response to stress, low oxygen and stationary phase. Previous work has shown that BCAS0292 is elevated in response to stress, low oxygen and stationary phase and elevated in late isolates from chronically colonised patients. The present work shows that absence of BCAS0292 reduces the abundance of ≥ 500 proteins, particularly those including those involved in the stress response, metabolic processes, transcription cell signaling, and virulence. In addition we have shown that the presence of BCAS0292 alters DNA topology, providing a mechanism by which it globally regulates protein expression in response to stress.

In summary, we propose a mechanism whereby BCAS0292 regulates expression of multiple groups of genes which facilitate its adaptation to stress (Figure 7). In this way, it allows an environmental pathogen such as *Burkholderia* which is exquisitely successful at thriving in a wide variety of niches from the rhizosphere, to the CF lung and disinfectants. The finding that two thirds of proteins previously shown to be upregulated in chronic infection isolates suggests that BCAS0292 may also play a role in the adaptation to the hypoxic CF lung. Future work will focus on identifying the probing the mechanism by which it mediates this effect.

## Experimental procedures

### Bacterial and epithelial cell culture maintenance

*B. cenocepacia* strain K56-2 was obtained from the BCCM/LMG, University of Ghent, Belgium and was routinely plated on to *B. cepacia* selective agar (BCSA) (Henry *et al.*, 1997). Bacteria were routinely grown in Luria Bertani (LB broth) at 37 °C with orbital agitation (150 rpm). The CFBE41o^−^ cell line, which is homozygous for the ΔF508 mutation, was maintained in fibronectin-vitrogen coated flasks containing minimal essential medium (MEM) with 10% FBS, 1% penicillin/ streptomycin, 1% L-glutamine and 1% non-essential amino acids and incubated in 5% CO_2_ environment at 37 °C (Wright *et al.*, 2011).

### Mutagenesis of *B. cenocepacia* K56-2 and complementation

Unmarked, non-polar gene deletions were constructed as previously described (Flannagan *et al.*, 2008). The deletion of BCAS0292 was confirmed by DNA sequencing of the PCR product spanning the deletion site. To complement *B. cenocepacia* K56-2_ΔBCAS0292, wild-type BCAS0292 was amplified from *B. cenocepacia* K56-2 (See primer pairs in Supplementary Table S1), digested with the restriction enzymes NdeI and XbaI and ligated into similarly digested pMH447. The complementation plasmid was introduced into the mutant by conjugation. Once transferred into the target mutant strain the complementation vector integrates into the genome at aminoglycoside efflux genes (*BCAL1674-BCAL1675*) (Hamad *et al.*, 2010). As before, pDAISce-I, was then introduced, resulting in the replacement of BCAL1674-1675 by BCAS0292. The complementation of BCAS0292 was confirmed by PCR and then confirmed by phenotype analysis.

### *mellonella* infection model

Log-phase grown bacteria were suspended to an starting OD_600nm_ in 0.1 in sterile PBS and serially diluted to 10^−7^. Each dilution (20 μl) was injected into the hindmost left proleg of *G. mellonella* larvae using a sterile Terumo 0.3 ml syringe (10 larvae per group) (Costello *et al.*, 2011). An equal volume of each dilution was also plated onto LB agar counted at 48 h to accurately quantify bioburden injected. Ten larvae injected with PBS alone served as controls. Larvae were incubated at 37 °C and examined for survival at 24, 48 and 72 h following injection.

### Cloning and expression of recombinant BCAS0292

The forward primer was designed to contain the ‘CACC’ sequence at the beginning of the primer to introduce a 5’ overhang allowing directional cloning of BCAS0292 into the pET100/ D-Topo vector in frame with the N-terminal 6xHis tag. PCR amplification was carried out using HotStar Taq polymerase (Qiagen, Germany) and amplicons cloned into the pET100/ D-Topo vector as previously described (Dennehy *et al.*, 2017). Each transformation reaction was spread on to pre-warmed selective plates containing 50 μg/ml of ampicillin and incubated overnight at 37°C and transformants identified by PCR. *E. coli* BL21 Star™ (DE3) One Shot® cells (Thermo Fisher Scientific) were used for protein expression (Dennehy *et al.*, 2017). Following the confirmation of recombinant BCAS0292 (rBCAS0292) expression in a pilot study, 1 L cultures of BL21 cells transformed with the expression plasmid, were grown in LB-ampicillin (100 μg/ ml) and induced with IPTG at a final concentration of 1 mM overnight at 25 °C. Bacteria were then pelleted using centrifugation at 2,500 g for 10 min. Supernatants were discarded, pellets weighed and stored at −80 °C.

### Recombinant BCAS0292 purification

Purification of rBCAS0292 was carried out as previously described (Dennehy *et al.*, 2017). Ice-cold lysis buffer containing 6 M Guanidine-HCL, 100 mM sodium phosphate, 10 mM Tris-HCL, (pH= 8) and 1X EDTA-free protease inhibitor was added to bacterial pellets at a volume of 30 ml/ 7 g of pellet. Cells were resuspended in lysis buffer and lysed on ice with a cell disruptor (maximum output, two 30-sec pulses). Cell debris and any remaining intact cells were pelleted using centrifugation at 4,800 x*g* for 35 min. Supernatants were filtered through a 0.45 μM filter and incubated with a rotating HisPur Ni-NTA column (ThermoFisher, Ireland), which was pre-equilibrated with buffer A, for 1 h at 4 °C. Unbound proteins were removed by a series of washes and bound protein was subsequently eluted in two steps with 250 mM and 300 mM imidazole. Following purification, the protein concentration of each fraction was determined using the Bradford assay. Protein purity was determined by SDS-PAGE.

### Attachment of bacteria to epithelial cells

To quantify the attachment of Bcc or BCAS0292-expressing *E. coli* strains to lung epithelial cells, CFBE41o^−^ cells were seeded in coated chamber slides 24 h prior to the 30 min incubation of individual bacterial strains at an MOI of 50:1, as previously described (Dennehy *et al.*, 2017). In the case of studies with recombinant *E. coli* strains, rBCAS0292 expression was induced in all strains (*E. coli* BL21, *E. coli* BL21_BCAS0292) with 1 mM IPTG for 3 h prior to incubation with epithelial cells. Unattached bacteria were removed by washing with sterile PBS four times for five min each. The CFBE41o^−^ cells and attached bacteria were fixed in 3 % paraformaldehyde (pH=7.2) for 10 min at room temperature. Cells were gently washed once with PBS and then blocked in PBS containing 5 % BSA for one hour at room temperature. Cells were then incubated overnight at 4 °C in PBS containing 1 % BSA and either a rabbit anti-Bcc antibody (1:800 dilution) or an anti-*E. coli* FITC-conjugated antibody (1:100 dilution). The following day cells were washed three times with PBS before the addition of FITC-conjugated anti-rabbit antibody (1:500 dilution), in the case of Bcc strains, for one h in the dark. The secondary antibody was removed, and cells washed twice for 5 min each with PBS. Cells were then counterstained with DAPI and phalloidin conjugated to Alexa fluor 568 (5U/ ml) for 15 min in the dark at a concentration of 1 μg/ml in PBS. A coverslip was applied to each chamber before bacteria and cells were visualised using confocal microscopy. Bacterial attachment was counted in 20 randomly selected fields for each strain and values were expressed as the number of bacteria/100 cells.

### Whole proteome analysis

To extract whole cell lysates were extracted from stationary phase cultures of three biological replicates of K56-2 WT and the ΔBCAS0292 mutant strains. Bacterial cells were pelleted with centrifugation of 6,000 *g* for ten min. Ice-cold lysis buffer containing 8 M Urea, 25 mM Tris-HCL, 10 mM DTT and 1 X EDTA-free protease inhibitor cocktail (pH=8.6) was added to pellets, using a volume of 8 ml/g bacterial pellet. Pellets were resuspended and cells lysed using a cell-disruptor probe set with an output power of 16, using four 30-sec bursts on ice. Lysates were then centrifuged at 14,000 x*g* for 15 min, supernatants transferred to fresh tubes and a Bradford assay used to determine the protein concentration of each sample. DTT (1M) was added to each supernatant (10 μl/ml lysate) and incubated at 56 °C for 30 min followed by 1M iodoacetamide (55 μl/ml lysate) which was incubated in darkness at RT for 20 min. Lysates were then dialysed in dialysis tubing with a cut-off of 3.5 kDa against 100 mM ammonium bicarbonate overnight with stirring at 4 °C then the ammonium bicarbonate was replaced with fresh solution and incubated for a further six hours. Trypsin (400 ng/μl) was added to dialysed protein (5 μl/100 μl protein) and tubes incubated overnight at 37 °C. Aliquots (20 μl) from each sample were placed in fresh tubes and samples were dried in a speedy vac using medium heat and resuspended in 20 μl resuspension buffer containing 0.5 % TFA in MilliQ water. Tubes were sonicated for two min and centrifuged briefly. ZipTips (Millipore, Germany) were wetted with 10 μl of wetting solution containing 0.1 % TFA in 80 % acetonitrile slowly five times. Equilibration solution (10 μl) containing 0.1 % TFA in MilliQ water was aspirated and dispensed into the ZipTip, five times before 10 μl of the resuspended sample was then aspirated and dispensed slowly into the ZipTip, 15 times. Then 10 μl of the washing solution containing 0.1 % TFA in MilliQ water was aspirated into the ZipTip and dispensed to waste. This step was repeated five times before 10 μl of the elution solution containing 0.1 % TFA in 60 % Acetonitrile was aspirated and dispensed from the ZipTip to a fresh 1.5 ml tube six times. Samples were then dried under medium heat and resuspended in 15 μl of loading buffer containing 0.05 % TFA in 2 % acetonitrile. Tubes were sonicated for 2 min, centrifuged briefly and samples transferred into fresh vials and 3 μl (1 μg protein) used for Q-Exactive analysis. All biological replicate and technical replicate samples were analysed on a Thermo Scientific Q Exactive mass spectrometer connected to a Dionex Ultimate 3000 (RSLCnano) chromatography system. Each sample was loaded onto an EASY-Spray PepMap RSLC C18 Column (50 cm x 75 μm, Thermo Scientific), and was separated by an increasing acetonitrile gradient over 120 min at a flow rate of 250 nL/min. The mass spectrometer was operated in positive ion mode with a capillary temperature of 220 ^☐^C, and with a potential of 2500 V applied to the frit. All data was acquired with the MS operating in automatic data dependent switching mode. A high resolution (140,000) MS scan (300-2000 Dalton) was performed using the Q-Exactive to select the 15 most intense ions prior to MS/MS analysis using HCD (higher-energy collisional dissociation). Protein identification and label free quantitative (LFQ) analysis was conducted using MaxQuant (version 1.2.2.5: http://maxquant.org/) supported by the Andromeda database search engine to correlate MS/MS data against the *B. cenocepacia* strain J2315 Uniprot database and Perseus to organise the data (Version 1.4.1.3). For protein identification the following search parameters were used: precursor-ion mass tolerance of 1.5 Da, fragment ion mass tolerance of 6 ppm with cysteine carbamidomethylation as a fixed modification and a maximum of 2 missed cleavage sites allowed. False Discovery Rates (FDR) were set to 0.01 for both peptides and proteins and only peptides with minimum length of six amino acid length were considered for identification. Proteins were considered identified when a minimum of two peptides for each parent protein was observed. LFQ intensities measured for individual runs were grouped based on their experimental treatment. The data was log2 transformed and an ANOVA (p < 0.05) was performed between the control and individual treatment samples to identify differentially abundant proteins. A log2 fold change ☐≥☐1.5 and log2 fold change☐≤☐−1.5 with adjusted p-value☐<☐0.05 were considered. Proteins with statistically significant differential expression were extracted and these were used to generate maps of expression. To improve visual representation of differentially abundant proteins, mean values were generated for each treatment and used to build the maps. A qualitative assessment was also conducted. This involved the identification of proteins that were completely absent in a specific strain. Those proteins that were completely missing from all replicates of a particular group but present in other groups were determined manually from the data matrix. These proteins are not considered statistically significant as the values for absences are given as NaN (not a number) which is not a valid value for an ANOVA analysis. However, the complete absence of a protein from a group may be biologically significant and these proteins are reported as qualitatively differentially expressed.

### Bacterial cell shape analysis

Cell shape was examined according to published methods (Shiomi *et al.*, 2008). Cells were grown at 37°C in LB overnight before reinoculation and culture to achieve log growth (OD600 of 0.5 – 0.7). Optical sectioning was performed with an Olympus FV-10 ASW confocal microscope using a Plan Apo X60 1.40 oil immersion objective lens and Olympus FLUOVIEW Ver.3.1 software. Section images were captured along the z-axis at 0.2 μm intervals.

### Circular dichroism spectroscopy

Circular dichroism measurements were carried out using a Jasco J-810 spectropolarimeter equipped with a Peltier temperature control system (Model PTC-423-S). The molar ellipticity per mean residue, [θ](deg cm^2^ x dmol^−1^), [θ](degcm2 × dmol^−1^) was calculated from the equation [θ] = [θ]obs x mrw x (10 x l x C)^−1^, [θ] = [θ]obs × mrw × (10 × l × C)^−1^,where [θ]obs is the ellipticity measured in degrees, mrw is the mean residue molecular mass (105.3 Da), C is the protein concentration in g x l^−1^ and l is the optical path length of the cell in cm. Far-UV spectra (190–260 nm) of recombinant BCAS0292 protein were recorded at 293K using a 0.1 cm optical path-length cell, with a protein concentration of 0.2 mg ml^−1^. For thermal denaturation experiments, ellipticity was monitored at 222 nm with a temperature slope of 1°C/min from 20°C to 100°C. Melting temperature (*Tm*) was calculated from the maximum of the first derivative of the unfolding curves.

### Light scattering Analysis

Purified rBCAS0292 was analyzed by size exclusion chromatography (SEC) coupled with light scattering (LS) using a DAWN MALS instrument and an Optilab rEX (Wyatt Technology). Eight hundred micrograms of purified rBCAS0292 was loaded into the column S75 16/60 (GE Healthcare, Italy), equilibrated in 50mM TrisHCl, 150mM NaCl, 5% Glycerol, pH 7.5 and. The on-line measurement of the intensity of the Rayleigh scattering as a function of the angle as well as the differential refractive index of the eluting peak in SEC was used to determine the weight average molar mass (MW) of eluted protein, using the Astra 5.3.4.14 (Wyatt Technologies, USA) software.

### Homology modelling

The homology model structure of BCAS0292 was built after consensus-based sequence comparison using HHPRED. Best model template was identified as the structure of the PixA inclusion body protein from *Burkholderia cenocepacia* (PDB code 4lzk, sequence identity 23.2%, E-value 7.2e-38). Using this alignment, the homology model was built using the program MODELLER (Bitencourt-Ferreira and de Azevedo, 2019). As a validation tool, an independent model was computed using I-Tasser (Yang and Zhang, 2015) and the two models compared. The meta-threading approach LOMETS was used to retrieve template proteins of similar folds from the PDB library. The program SPICKER was used to cluster the decoys based on the pair-wise structure similarity. Best confidence model presented a C-score −0.34 (Yang and Zhang, 2015). The two approaches provided consistent models, with RMSD computed on Cα atoms of 1.5 ◻. Based on the estimated local accuracy of the model computed by i-TASSER, the C-terminal 10 residues were removed from the model. Electrostatic potential surface was computed using the program Chimera (Yang *et al.*, 2012).

### Chloroquine gel electrophoresis

*B. cenocepacia* strains were transformed with pDA-12 reporter plasmid and cultured on plates containing tetracycline (150 μg/ml), ampicillin (100μg/ml) and polymyxin B (50 μg/ml). The reporter plasmids were extracted from either log or stationary phase cultures using a Qiagen Miniprep kit (ThermoFisher, Germany). The plasmids were analyzed on chloroquine gels (1.5% agarose, 10 μg/ml chloroquine) separated for 24 h at 100V, 150mA (Scanlan *et al.*, 2017). The gels were washed 8 to 12 times over 8 h to remove chloroquine, before staining with ethidium bromide (10 μg/ml) for 2 h. The gel was washed for 1 h before imaging under UV light (Scanlan *et al.*, 2017).

### Statistical analysis

Analysis of virulence data was performed using a log rank (Mantel-Cox) test on the survival curves. Statistical analysis of host cell attachment, biofilm formation and hypoxic growth and survival were carried out by two-way ANOVA using Prism software.

## Acknowledgements

RD was funded by Science Foundation Ireland (RFP-BMT-3307); SD was funded by TU Dublin President’s Award; SMcCormack was funded by a UCD SBBS research scholarship; RD acknowledges funding from EU COST Action BM1003: Microbial cell surface determinants of virulence as targets for new therapeutics in cystic fibrosis for Short Term Scientific Missions to the laboratories of MAV and RB. RB acknowledges the Italian MIUR, project 2017SFBFE. The authors acknowledge Aoibhín Dwan for support in proteomic analysis.

## Data Availability

The data that support the findings of this study are available from the corresponding author upon reasonable request.

## References

Altschul, S. F., Madden, T. L., Schaffer, A. A., Zhang, J., Zhang, Z., Miller, W., et al. 1997. Gapped BLAST and PSI-BLAST: a new generation of protein database search programs. Nucleic Acids Res, 25, 3389–402.

Aravind, L., Anantharaman, V., Balaji, S., Babu, M. M. & Iyer, L. M. 2005. The many faces of the helix-turn-helix domain: transcription regulation and beyond. FEMS Microbiol Rev, 29, 231–62.

Bach, E., Sant’anna, F. H., Magrich Dos Passos, J. F., Balsanelli, E., De Baura, V. A., Pedrosa, F. O., et al. 2017. Detection of misidentifications of species from the Burkholderia cepacia complex and description of a new member, the soil bacterium Burkholderia catarinensis sp. nov. Pathog Dis, 75.

Bitencourt-Ferreira, G. & De Azevedo, W. F., Jr. 2019. Homology Modeling of Protein Targets with MODELLER. Methods Mol Biol, 2053, 231–249.

Caraher, E., Duff, C., Mullen, T., Mc Keon, S., Murphy, P., Callaghan, M., et al. 2007. Invasion and biofilm formation of Burkholderia dolosa is comparable with Burkholderia cenocepacia and Burkholderia multivorans. J Cyst Fibros, 6, 49–56.

Costello, A., Herbert, G., Fabunmi, L., Schaffer, K., Kavanagh, K. A., Caraher, E. M., et al. 2011. Virulence of an emerging respiratory pathogen, genus Pandoraea, in vivo and its interactions with lung epithelial cells. J Med Microbiol, 60, 289–99.

Cullen, L., O’connor, A., Drevinek, P., Schaffer, K. & McClean, S. 2017. Sequential Burkholderia cenocepacia Isolates from Siblings with Cystic Fibrosis Show Increased Lung Cell Attachment. Am J Respir Crit Care Med, 195, 832–835.

Cullen, L., O’connor, A., Mccormack, S., Owens, R. A., Holt, G. S., Collins, C., et al. 2018. The involvement of the low-oxygen-activated locus of Burkholderia cenocepacia in adaptation during cystic fibrosis infection. Scientific Reports, 8, 13386.

De Oliveira, D. M. P., Forde, B. M., Kidd, T. J., Harris, P. N. A., Schembri, M. A., Beatson, S. A., et al. 2020. Antimicrobial Resistance in ESKAPE Pathogens. Clin Microbiol Rev, 33.

Dennehy, R., Romano, M., Ruggiero, A., Mohamed, Y. F., Dignam, S. L., Mujica Troncoso, C., et al. 2017. The Burkholderia cenocepacia peptidoglycan-associated lipoprotein is involved in epithelial cell attachment and elicitation of inflammation. Cell Microbiol, 19.

Dorman, C. J. & Corcoran, C. P. 2009. Bacterial DNA topology and infectious disease. Nucleic Acids Res, 37, 672–8.

Flannagan, R. S., Linn, T. & Valvano, M. A. 2008. A system for the construction of targeted unmarked gene deletions in the genus Burkholderia. Environ Microbiol, 10, 1652–60.

Goetsch, M., Owen, H., Goldman, B. & Forst, S. 2006. Analysis of the PixA inclusion body protein of Xenorhabdus nematophila. J Bacteriol, 188, 2706–10.

Hamad, M. A., Skeldon, A. M. & Valvano, M. A. 2010. Construction of aminoglycoside-sensitive Burkholderia cenocepacia strains for use in studies of intracellular bacteria with the gentamicin protection assay. Appl Environ Microbiol, 76, 3170–6.

Hegde, S. S., Vetting, M. W., Roderick, S. L., Mitchenall, L. A., Maxwell, A., Takiff, H., et al. 2005. A fluoroquinolone resistance protein from Mycobacterium tuberculosis that mimics DNA. Science, 308, 1480–3.

Henry, D. A., Campbell, M. E., Lipuma, J. J. & Speert, D. P. 1997. Identification of Burkholderia cepacia isolates from patients with cystic fibrosis and use of a simple new selective medium. J Clin Microbiol, 35, 614–9.

Holden, M. T., Seth-Smith, H. M., Crossman, L. C., Sebaihia, M., Bentley, S. D., Cerdeno-Tarraga, A. M., et al. 2009. The genome of Burkholderia cenocepacia J2315, an epidemic pathogen of cystic fibrosis patients. J Bacteriol, 191, 261–77.

Huang, M. F., Lin, S. J., Ko, T. P., Liao, Y. T., Hsu, K. C. & Wang, H. C. 2017. The monomeric form of Neisseria DNA mimic protein DMP19 prevents DNA from binding to the histone-like HU protein. PLoS One, 12, e0189461.

Huber, B., Feldmann, F., Kothe, M., Vandamme, P., Wopperer, J., Riedel, K., et al. 2004. Identification of a novel virulence factor in Burkholderia cenocepacia H111 required for efficient slow killing of Caenorhabditis elegans. Infect Immun, 72, 7220–30.

Ispolatov, I., Yuryev, A., Mazo, I. & Maslov, S. 2005. Binding properties and evolution of homodimers in protein-protein interaction networks. Nucleic Acids Res, 33, 3629–35.

Kooi, C., Subsin, B., Chen, R., Pohorelic, B. & Sokol, P. A. 2006. Burkholderia cenocepacia ZmpB is a broad-specificity zinc metalloprotease involved in virulence. Infect Immun, 74, 4083–93.

Lucas, J., Goetsch, M., Fischer, M. & Forst, S. 2018. Characterization of the pixB gene in Xenorhabdus nematophila and discovery of a new gene family. Microbiology, 164, 495–508.

Mahenthiralingam, E., Baldwin, A. & Dowson, C. G. 2008. Burkholderia cepacia complex bacteria: opportunistic pathogens with important natural biology. J Appl Microbiol, 104, 1539–51.

Mares, J., Kumaran, S., Gobbo, M. & Zerbe, O. 2009. Interactions of lipopolysaccharide and polymyxin studied by NMR spectroscopy. J Biol Chem, 284, 11498–506.

McClean, S., Healy, M. E., Collins, C., Carberry, S., O’shaughnessy, L., Dennehy, R., et al. 2016. Linocin and OmpW Are Involved in Attachment of the Cystic Fibrosis-Associated Pathogen Burkholderia cepacia Complex to Lung Epithelial Cells and Protect Mice against Infection. Infect Immun, 84, 1424–37.

Mhatre, E., Snyder, D. J., Sileo, E., Turner, C. B., Buskirk, S. W., Fernandez, N. L., et al. 2020. One gene, multiple ecological strategies: a biofilm regulator is a capacitor for sustainable diversity. bioRxiv, 2020.05.02.074534.

O’ Croinin, T., Carroll, R. K., Kelly, A. & Dorman, C. J. 2006. Roles for DNA supercoiling and the Fis protein in modulating expression of virulence genes during intracellular growth of Salmonella enterica serovar Typhimurium. Mol Microbiol, 62, 869–82.

O’grady, E. P. & Sokol, P. A. 2011. Burkholderia cenocepacia differential gene expression during host-pathogen interactions and adaptation to the host environment. Front Cell Infect Microbiol, 1, 15.

O’grady, E. P., Viteri, D. F., Malott, R. J. & Sokol, P. A. 2009. Reciprocal regulation by the CepIR and CciIR quorum sensing systems in Burkholderia cenocepacia. BMC Genomics, 10, 441.

Peeters, C., Zlosnik, J. E., Spilker, T., Hird, T. J., Lipuma, J. J. & Vandamme, P. 2013. Burkholderia pseudomultivorans sp. nov., a novel Burkholderia cepacia complex species from human respiratory samples and the rhizosphere. Syst Appl Microbiol, 36, 483–9.

Roszniowski, B., McClean, S. & Drulis-Kawa, Z. 2018. Burkholderia cenocepacia Prophages-Prevalence, Chromosome Location and Major Genes Involved. Viruses, 10.

Saini, L. S., Galsworthy, S. B., John, M. A. & Valvano, M. A. 1999. Intracellular survival of Burkholderia cepacia complex isolates in the presence of macrophage cell activation. Microbiology, 145 (Pt 12), 3465–75.

Sass, A., Marchbank, A., Tullis, E., Lipuma, J. J. & Mahenthiralingam, E. 2011. Spontaneous and evolutionary changes in the antibiotic resistance of Burkholderia cenocepacia observed by global gene expression analysis. BMC Genomics, 12, 373.

Sass, A. M., Schmerk, C., Agnoli, K., Norville, P. J., Eberl, L., Valvano, M. A., et al. 2013. The unexpected discovery of a novel low-oxygen-activated locus for the anoxic persistence of Burkholderia cenocepacia. Isme J, 7, 1568–81.

Scanlan, E., Ardill, L., Whelan, M. V., Shortt, C., Nally, J. E., Bourke, B., et al. 2017. Relaxation of DNA supercoiling leads to increased invasion of epithelial cells and protein secretion by Campylobacter jejuni. Mol Microbiol, 104, 92–104.

Schmerk, C. L. & Valvano, M. A. 2013. Burkholderia multivorans survival and trafficking within macrophages. J Med Microbiol, 62, 173–184.

Shinoy, M., Dennehy, R., Coleman, L., Carberry, S., Schaffer, K., Callaghan, M., et al. 2013. Immunoproteomic analysis of proteins expressed by two related pathogens, Burkholderia multivorans and Burkholderia cenocepacia, during human infection. PLoS One, 8, e80796.

Shiomi, D., Sakai, M. & Niki, H. 2008. Determination of bacterial rod shape by a novel cytoskeletal membrane protein. EMBO J, 27, 3081–91.

Shortt, C., Scanlan, E., Hilliard, A., Cotroneo, C. E., Bourke, B. & T, O. C. 2016. DNA Supercoiling Regulates the Motility of Campylobacter jejuni and Is Altered by Growth in the Presence of Chicken Mucus. MBio, 7.

Summer, E. J., Gonzalez, C. F., Carlisle, T., Mebane, L. M., Cass, A. M., Savva, C. G., et al. 2004. Burkholderia cenocepacia phage BcepMu and a family of Mu-like phages encoding potential pathogenesis factors. J Mol Biol, 340, 49–65.

Valvano, M. A. 2015. Intracellular survival of Burkholderia cepacia complex in phagocytic cells. Can J Microbiol, 61, 607–15.

Wang, H. C., Chou, C. C., Hsu, K. C., Lee, C. H. & Wang, A. H. 2018. New paradigm of functional regulation by DNA mimic proteins: Recent updates. IUBMB Life.

Wang, H. C., Ho, C. H., Hsu, K. C., Yang, J. M. & Wang, A. H. 2014. DNA mimic proteins: functions, structures, and bioinformatic analysis. Biochemistry, 53, 2865–74.

Wang, H. C., Ko, T. P., Wu, M. L., Ku, S. C., Wu, H. J. & Wang, A. H. 2012. Neisseria conserved protein DMP19 is a DNA mimic protein that prevents DNA binding to a hypothetical nitrogen-response transcription factor. Nucleic Acids Res, 40, 5718–30.

Wang, H. C., Wu, M. L., Ko, T. P. & Wang, A. H. 2013. Neisseria conserved hypothetical protein DMP12 is a DNA mimic that binds to histone-like HU protein. Nucleic Acids Res, 41, 5127–38.

Winsor, G. L., Khaira, B., Van Rossum, T., Lo, R., Whiteside, M. D. & Brinkman, F. S. 2008. The Burkholderia Genome Database: facilitating flexible queries and comparative analyses. Bioinformatics, 24, 2803–4.

Wright, C., Pilkington, R., Callaghan, M. & McClean, S. 2011. Activation of MMP-9 by human lung epithelial cells in response to the cystic fibrosis-associated pathogen Burkholderia cenocepacia reduced wound healing in vitro. Am J Physiol Lung Cell Mol Physiol, 301, L575–86.

Yang, J. & Zhang, Y. 2015. I-TASSER server: new development for protein structure and function predictions. Nucleic Acids Res, 43, W174–81.

Yang, Z., Lasker, K., Schneidman-Duhovny, D., Webb, B., Huang, C. C., Pettersen, E., et al. 2012. UCSF Chimera, MODELLER, and IMP: an integrated modeling system. J Struct Biol, 179, 269–78.

Yano, H., Kuga, A., Okamoto, R., Kitasato, H., Kobayashi, T. & Inoue, M. 2001. Plasmid-encoded metallo-beta-lactamase (IMP-6) conferring resistance to carbapenems, especially meropenem. Antimicrob Agents Chemother, 45, 1343–8.

Zhanhua, C., Gan, J. G., Lei, L., Sakharkar, M. K. & Kangueane, P. 2005. Protein subunit interfaces: heterodimers versus homodimers. Bioinformation, 1, 28–39.

Zhou, J., Chen, Y., Tabibi, S., Alba, L., Garber, E. & Saiman, L. 2007. Antimicrobial susceptibility and synergy studies of Burkholderia cepacia complex isolated from patients with cystic fibrosis. Antimicrob Agents Chemother, 51, 1085–8.

Zoued, A., Durand, E., Santin, Y. G., Journet, L., Roussel, A., Cambillau, C., et al. 2017. TssA: The cap protein of the Type VI secretion system tail. Bioessays, 39.

