## Supplementary material for "The DNA mimic protein BCAS0292 is involved in the regulation of virulence of *Burkholderia cenocepacia*": Suppl tables S1 and S2

**Supplementary tables**

**Table S1: Strains and plasmids used in this study**

| **Strain** | **Characteristics** | **Origin** |
|---|---|---|
| ***B. cenocepacia* strains** | | |
| K56-2 (LMG18863) | ET12 clone | CF clinical isolate, Canada (epidemic strain) |
| Δ*BCAS0292* | Deletion of BCAS0292 in K56-2 | This study |
| Δ*BCAS0292*(*pBCAS0292Bc*) | BCAS0292 integration in Δ*BCAS0292* | This study |
| ***E. coli* strains** | | |
| NCIB9485 | | Laboratory strain |
| GT115 | $F^-$ *mcrA* Δ(*mrr-hsd*RMS-*mcrBC*) Φ80Δ*lac*ZΔM15 Δ*lac*X74 *recA1 rpsL* (StrA) *endA1*Δ*dcm uidA* (ΔMIuI)*::pir-116* Δ*sbcC-sbcD* | Invivogen |
| SY327 | *araD Δ(lac-pro) argE(Am) recA56 nalA λ pir; Rif$^r$* | (Miller and Mekalanos, 1988) |
| DH5α | *$F^-$ ϕ80dlacZΔM15 Δ(lacZYA-argF)U169 endA1 recA1 hsdR17(rK$^-$ mK$^+$) supE44 thi-1 ΔgyrA96 relA1* | Invitrogen |
| Top10 | *$F^-$ mcrA Δ(mrr-hsdRMS-mcrBC) Φ80lacZΔM15 ΔlacX74 recA1 araD139 Δ(ara-leu)7697 galU galK rpsL (Str$^R$) endA1 nupG* | Invitrogen |
| BL21 Star™ (DE3) | $F^-$ *ompT hsdS*$_B$ ($r_B^- m_B^-$) *gal dcm rne131* (DE3) | Invitrogen |
| **Plasmids** | | |
| pGPISceI-2 | *ori*$_{R6K}$, Ω$Tp^r$ , $mob^+$, containing the ISce-I restriction site | (Flannagan *et al.*, 2008) |

| | | |
|---|---|---|
| pRK2013 | *ori*$_{\mathrm{COIEI}}$, RK2 derivative, Kan$^{r}$, mob$^{+}$, tra$^{+}$ | (Figurski and Helinski, 1979) |
| pDAI-SceI | Encodes the ISce-I homing endonuclease, Tet$^{r}$ | (Flannagan *et al.*, 2008) |
| pMH447 | pGPI-SceI derivative used for chromosomal complementation | (Hamad *et al.*, 2012) |
| pET100/D-Topo | Peptide fusion at N-terminal, 6xHis, Amp$^{r}$ | Invitrogen |
| Need reporter plasmid for supercoiling | | |

**Table S2: Primers used in this study**

| **Region:** | **Primer Sequence (5'-3'):** | **Product size (bp)** | **Restriction enzymes:** |
|---|---|---|---|
| **BCAS0292 gene** | | | |
| US | actgacgaattccttcgaaaaccaggtggtgtt<br>agtcagatcgatagaccatcgcaacgaacacca | 392 | *EcoRI*<br>*ClaI* |
| DS | gacactatcgatcgatgaaagaccatttctggc<br>cagagtgctagcacacgatcttcagcgtgaact | 544 | *ClaI*<br>*NheI* |
| Internal | cgccagaaatggtctttcat<br>ggtgaccgtgctgttcatc | 150 | N/A<br>N/A |
| Sequencing | gttccagatcattgatcgcaa<br>atcaatgcgctgatcgcgaac | 534 | N/A<br>N/A |
| Cloning | caccatgtctggtgttcgttgc<br>atccggctacggttcgacgcc | 553 | N/A<br>N/A |
| Complementation | ttttcatatgtctggtgttcgttgcgat<br>gttttctagactacggttcgacgccggg | 560 | *NdeI*<br>*XbaI* |
| **Plasmids** | | | |
| pGPI-Scel-2 | gagtgacacaggaacact | 347 | N/A |

| | | | |
|---|---|---|---|
| | gctcaatcaatcaccggatccc | | N/A |
| pMH447 | ttgatggcgagcgattcttc<br>ccagttcttcagcgtgacga | 347 | N/A<br>N/A |
