## Supplemental Table S3 for "The DNA mimic protein BCAS0292 is involved in the regulation of virulence of *Burkholderia cenocepacia*"

Suppl Table S3: All proteins showing altered abundance by $\geq$ 1.5 fold ($p < 0.05$) in the ΔBCAS0292 mutant strain relative to the K56-2 wild type strain in all replicates.

| | **Protein names** | **UniProt code** | **Fold difference** | **Seq. Len** | **#GOs** |
|---|---|---|---|---|---|
| **Reduced abundance in the mutant strain** | | | | | |
| BCAM0278 | Putative heat shock protein | B4EI89_BURCJ | 275.96 | **144** | 3 |
| pBCA087 | hydrolase family protein | B4EQM2_BURCJ | 62.02 | 237 | 0 |
| BCAS0551 | HU DNA-binding protein | B4ENV0_BURCJ | 60.11 | 90 | 5 |
| BCAL3228 | Uncharacterized protein | B4EDE5_BURCJ | 59.52 | 181 | 0 |
| BCAS0293 aidA | Nematocidal protein AidA | B4EQ59_BURCJ | 49.84 | 167 | 0 |
| BCAM2547a | Glyoxalase/bleomycin resistance protein/dioxygenase superfamily protein | B4EJA1_BURCJ | 33.16 | 148 | 0 |
| BCAS0636 | Uncharacterized protein | B4EP30_BURCJ | 29.03 | 95 | 0 |
| BCAL2401 | Putative exported protein | B4E6C7_BURCJ | 25.39 | 126 | 0 |
| BCAS0191 | Putative endoribonuclease | B4EPV8_BURCJ | 17.40 | 162 | 0 |
| BCAM1261 | Putative membrane protein | B4EHE0_BURCJ | 12.52 | 287 | 0 |
| BCAM1920 | Hypothetical phage protein | B4EMI3_BURCJ | 12.46 | 253 | 0 |
| BCAM0843 | Putative lipoprotein | B4EMP5_BURCJ | 10.48 | 194 | 0 |
| BCAM2752 | NAD dependent epimerase/dehydratase family protein | B4ELF1_BURCJ | 9.53 | 251 | 22 |
| BCAM1883 | Hypothetical phage protein | B4EME7_BURCJ | 9.46 | 300 | 0 |
| BCAM1919 | Hypothetical phage protein | B4EMI2_BURCJ | 8.52 | 138 | 0 |
| BCAM0974 | Glyoxalase/bleomycin resistance protein/dioxygenase superfamily protein | B4EFB9_BURCJ | 8.01 | 129 | 1 |
| BCAM1407 | DJ-1/PfpI family protein | B4EII3_BURCJ | 7.57 | 228 | 2 |
| BCAM0292 | Putative universal stress protein | B4EIA4_BURCJ | 6.51 | 167 | 5 |
| BCAM2129 | 2-amino-3-carboxymuconate 6-semialdehyde decarboxylase | B4EFS6_BURCJ | 6.45 | 344 | 2 |
| BCAM0308 | Uncharacterized protein | B4EIC0_BURCJ | 6.34 | 169 | 0 |
| BCAM0307 | Uncharacterized protein | B4EIB9_BURCJ | 6.28 | 215 | 0 |
| BCAL1529 | Flp pilus type assembly-related protein | B4E825_BURCJ | 6.19 | 412 | 0 |
| BCAM2478 | Serine-carboxyl peptidase | B4EIP4_BURCJ | 6.04 | 529 | 14 |
| BCAM2610 | Putative methyltransferase | B4EK96_BURCJ | 5.99 | 221 | 31 |
| BCAM1547 | Glyoxalase/bleomycin resistance protein/dioxygenase superfamily protein | B4EJN9_BURCJ | 5.99 | 148 | 0 |
| BCAM0042 | Putative aldo/keto reductase | B4EG28_BURCJ | 5.99 | 315 | 2 |

| | | | | | |
|---|---|---|---|---|---|
| BCAM0844 | Uncharacterized protein | \|B4EMP6_BURCJ | 5.73 | 128 | 4 |
| BCAS0742 | Uncharacterized protein | \|B4EPD0_BURCJ | 5.59 | 123 | 0 |
| BCAM0944 | Putative lipoprotein | \|B4EF89_BURCJ | 5.48 | 244 | 7 |
| BCAL3205a | Putative phage-related regulator | \|B4EDC3_BURCJ | 5.43 | 106 | 0 |
| BCAM2625 | Uncharacterized protein | \|B4EKB0_BURCJ | 5.41 | 75 | 0 |
| BCAM1408 | Xenobiotic reductase | \|B4EII4_BURCJ | 5.36 | 369 | 1 |
| BCAL0833 phbB | Putative Acetoacetyl-CoA reductase | \|B4EAU4_BURCJ | **5.28** | 247 | 6 |
| BCAL1978 | Putative pyrazinamidase/nicotinamidase | \|B4EBK5_BURCJ | 5.25 | 210 | 6 |
| BCAL1300 | Uncharacterized protein | \|B4E5I5_BURCJ | 4.98 | 144 | 0 |
| BCAM1824 | Uncharacterized protein | \|B4EM87_BURCJ | 4.38 | 116 | 0 |
| BCAL2703 | Uncharacterized protein | \|B4E8X6_BURCJ | 4.37 | 223 | 0 |
| BCAM0285 | Uncharacterized protein | \|B4EI97_BURCJ | 4.30 | 335 | 0 |
| BCAM2756 cblR | Two-component regulatory system, response regulator protein | \|B4ELF5_BURCJ | 4.30 | 238 | 1 |
| BCAL2962 | Uncharacterized protein | \|B4EB52_BURCJ | 4.20 | 232 | 0 |
| BCAL0337 tssL | Putative type VI secretion system protein TssL | \|B4E6Q9_BURCJ | 4.20 | 260 | 0 |
| BCAL0208 | AnsC family regulatory protein | \|B4E594_BURCJ | 4.18 | 174 | 7 |
| BCAL2295 | Putative chromosome condensation protein | \|B4EEN4_BURCJ | 4.18 | 290 | 18 |
| BCAM2155 | Putative lyase/mutase | \|B4EFV2_BURCJ | 4.15 | 286 | 2 |
| BCAL1985 | Putative exported isomerase | \|B4EBL2_BURCJ | 4.11 | 260 | 2 |
| BCAM1412 | Uncharacterized protein | \|B4EII8_BURCJ | 4.11 | 184 | 0 |
| BCAL3183 | Putative hydrolase | \|B4ECX4_BURCJ | 4.09 | 433 | 5 |
| BCAL2027 | Uncharacterized protein | \|B4EC34_BURCJ | 4.06 | 69 | 1 |
| BCAM0971 | Uncharacterized protein | \|B4EFB6_BURCJ | 4.04 | 89 | 0 |
| BCAL1623 | Uncharacterized protein | \|B4E8P6_BURCJ | 4.01 | 65 | 0 |
| BCAM0679 | Uncharacterized protein | \|B4ELN0_BURCJ | 4.01 | 127 | 0 |
| pBCA017 | Uncharacterized protein | \|B4EQQ1_BURCJ | 3.81 | 470 | 1 |
| BCAL2607 | Putative exported protein | \|B4E7U6_BURCJ | 3.76 | 184 | 0 |
| BCAM173 | Putative oxidoreductase | \|B4ELB3_BURCJ | 3.68 | 539 | 2 |
| BCAM0829 | Putative riboflavin synthase alpha chain | \|B4EMN1_BURCJ | 3.65 | 237 | 4 |
| BCAL0395 | Putative adenylate cyclase | \|B4E7A6_BURCJ | 3.63 | 208 | 2 |
| BCAL1233 | Putative heat shock Hsp20-related protein | \|B4EEH0_BURCJ | 3.61 | 136 | 1 |
| BCAM0315 | Putative exported protein | \|B4EIQ6_BURCJ | 3.58 | 158 | 0 |
| BCAL3041 malE | Maltose-binding protein | \|B4EBS5_BURCJ | 3.53 | 415 | 6 |

| | | | | | |
|---|---|---|---|---|---|
| BCAL0970 sdhB | Succinate dehydrogenase iron-sulfur protein | \|B4EFB5_BURCJ | 3.48 | 233 | 11 |
| BCAM2690 | Putative thioesterase | \|B4EKV9_BURCJ | 3.47 | 147 | 4 |
| BCAM2289 | Uncharacterized protein | \|B4EH05_BURCJ | 3.44 | 74 | 0 |
| BCAM2099 | Putative Pfp1 family protein | \|B4EFG7_BURCJ | 3.44 | 227 | 1 |
| BCAL1508 rbfA | Ribosome-binding factor A | \|RBFA_BURCJ | 3.37 | 132 | 2 |
| BCAM1726 | Putative exported protein | \|B4ELA4_BURCJ | 3.33 | 133 | 0 |
| BCAM0284 | Putative cytochrome c | \|B4EI96_BURCJ | 3.30 | 111 | 2 |
| BCAL0831 | Putative storage protein | \|B4EAU2_BURCJ | 3.29 | 194 | 0 |
| BCAM1871 | Uncharacterized protein | \|B4EMD3_BURCJ | 3.27 | 254 | 0 |
| BCAL1868 | Uncharacterized protein | \|B4EAG7_BURCJ | 3.26 | 71 | 0 |
| BCAS0409 zmpA | Zinc metalloprotease ZmpA | \|B4ENF7_BURCJ | 3.26 | 565 | 7 |
| BCAL0807 | ATP-dependent protease | \|B4EAR9_BURCJ | 3.25 | 211 | 32 |
| BCAL2123 | Uncharacterized protein | \|B4ECR4_BURCJ | 3.25 | 137 | 5 |
| BCAL1973 | Endoribonuclease L-PSP family protein | \|B4EBK1_BURCJ | 3.25 | 131 | 7 |
| BCAM2477 | Serine peptidase, family S10 | \|B4EIP3_BURCJ | 3.23 | 558 | 9 |
| BCAL1648 | Putative stress-related protein | \|B4E8S1_BURCJ | 3.23 | 165 | 7 |
| BCAM1465 | Putative exported protein | \|B4EJ19_BURCJ | 3.23 | 150 | 0 |
| BCAL1910acoB | Acetoin:2,6-dichlorophenolindophenol oxidoreductase beta subunit | \|B4EAZ6_BURCJ | 3.21 | 334 | 3 |
| BCAL3136 apaH | Bis(5-nucleosyl)-tetraphosphatase, symmetrical | \|B4ECS7_BURCJ | 3.21 | 279 | 2 |
| BCAL2669 | Putative exported protein | \|B4E8G2_BURCJ | 3.17 | 246 | 0 |
| BCAL2079 lpxA | Acyl-[acyl-carrier-protein]--UDP-N-acetylglucosamine O-acyltransferase | \|LPXA_BURCJ | 3.12 | 262 | 6 |
| BCAL1399 | OsmC-like protein | \|B4E6B9_BURCJ | 3.08 | 130 | 0 |
| BCAL0702 | Putative ferredoxin | \|B4E9Q2_BURCJ | 3.06 | 105 | 1 |
| BCAM0696 | Putative carboxymuconolactone decarboxylase | \|B4ELP7_BURCJ | 3.06 | 132 | 1 |
| BCAL1241 | Putative malonate decarboxylase delta-subunit | \|B4EEH8_BURCJ | 3.04 | 104 | 5 |
| BCAL1631 | Uncharacterized protein | \|B4E8Q4_BURCJ | 3.02 | 102 | 0 |
| BCAM0176 | AnsC family regulatory protein | \|B4EH77_BURCJ | 3.02 | 162 | 7 |
| BCAL2969a | Uncharacterized protein | \|B4EB66_BURCJ | 3.01 | 85 | 0 |
| BCAM1495 | Universal stress protein | \|B4EJI6_BURCJ | 2.99 | 156 | 4 |
| BCAL0056 | AraC family regulatory protein | \|B4EF09_BURCJ | 2.97 | 293 | 5 |
| BCAL0805 mutM | Formamidopyrimidine-DNA glycosylase | \|FPG_BURCJ | 2.97 | 275 | 6 |
| BCAL0849 | Metallo peptidase, subfamily M48B | \|B4EAW0_BURCJ | 2.95 | 253 | 3 |
| BCAL1798 | Putative exported protein | \|B4E9X9_BURCJ | 2.95 | 187 | 0 |

| | | | | | |
|---|---|---|---|---|---|
| BCAL1298 | Uncharacterized protein | \|B4E5I3_BURCJ | 2.92 | 349 | 0 |
| BCAM1857 | Uncharacterized protein | \|B4EMB9_BURCJ | 2.89 | 159 | 0 |
| BCAL0690 | Uncharacterized protein | \|B4E9P0_BURCJ | 2.87 | 292 | 0 |
| BCAM0953 | Extracellular solute-binding protein | \|B4EF98_BURCJ | 2.85 | 348 | 1 |
| BCAL0378 | Putative hydrolase | \|B4E6V1_BURCJ | 2.85 | 183 | 1 |
| BCAM0729 | Uncharacterized protein | \|B4EM31_BURCJ | 2.85 | 179 | 0 |
| BCAL2642 dcd | Deoxycytidine triphosphate deaminase | \|DCD_BURCJ | 2.83 | 189 | 3 |
| bcas0729 deoA | Thymidine phosphorylase | \|TYPH_BURCJ | 2.83 | 438 | 2 |
| BCAL0319 hisI | Phosphoribosyl-AMP cyclohydrolase | \|HIS3_BURCJ | 2.80 | 138 | 4 |
| BCAS0758 | Uncharacterized protein | \|B4EPE6_BURCJ | 2.78 | 151 | 0 |
| BCAL3204 tolB | Protein TolB | \|TOLB_BURCJ | 2.76 | 431 | 2 |
| BCAL0735 ptsH | Phosphocarrier protein HPr | \|B4EA60_BURCJ | 2.76 | 89 | 7 |
| BCAM0504 | CsbD-like protein | \|B4EK06_BURCJ | 2.75 | 65 | 0 |
| BCAM1482 | Uncharacterized protein | \|B4EJ35_BURCJ | 2.75 | 131 | 0 |
| BCAL0436 | Glyoxalase/bleomycin resistance protein/dioxygenase superfamily protein | \|B4E7E4_BURCJ | 2.75 | 137 | 3 |
| BCAM0623 | Two-component regulatory system, response regulator protein | \|B4EL44_BURCJ | 2.75 | 215 | 5 |
| BCAL2119 | Universal stress protein family protein | \|B4ECR0_BURCJ | 2.74 | 144 | 1 |
| BCAS0646A | Putative DNA-binding protein | \|B4EP39_BURCJ | 2.73 | 116 | 4 |
| BCAM1619 | Putative DNA-binding cold-shock protein | \|B4EK82_BURCJ | 2.70 | 67 | 10 |
| BCAL2418 | Putative exported protein | \|B4E6E4_BURCJ | 2.70 | 390 | 5 |
| BCAS0252 | DJ-1/PfpI family protein | \|B4EQ20_BURCJ | 2.70 | 199 | 0 |
| BCAM0722 | Putative O-methyltransferase | \|B4EM24_BURCJ | 2.70 | 223 | 6 |
| BCAM1472 | Putative glycosyltransferase | \|B4EJ25_BURCJ | 2.69 | 422 | 4 |
| BCAM2159 | Putative exported protein | \|B4EFV6_BURCJ | 2.69 | 117 | 2 |
| BCAM1098 | NUDIX hydrolase | \|B4EG79_BURCJ | 2.69 | 141 | 2 |
| BCAM0165 | Uncharacterized protein | \|B4EH65_BURCJ | 2.69 | 144 | 0 |
| BCAL0669 | Uncharacterized protein | \|B4E9L9_BURCJ | 2.68 | 144 | 0 |
| BCAL3205 | Putative exported protein | \|B4EDC2_BURCJ | 2.68 | 249 | 0 |
| BCAL3364 | Putative gluconokinase | \|B4EEV2_BURCJ | 2.67 | 167 | 6 |
| BCAM0277 | Uncharacterized protein | \|B4EI88_BURCJ | 2.64 | 91 | 0 |
| BCAM1823 | Putative methyltransferase | \|B4EM86_BURCJ | 2.60 | 287 | 3 |
| BCAL0320 hisE | Phosphoribosyl-ATP pyrophosphatase | \|HIS2_BURCJ | 2.59 | 121 | 5 |
| BCAM0380 | Putative exported protein | \|B4EIW8_BURCJ | 2.58 | 167 | 0 |

| | | | | | |
|---|---|---|---|---|---|
| BCAL1715 | Uncharacterized protein | \|B4E9B3_BURCJ | 2.58 | 201 | 0 |
| BCAL1360 | Uncharacterized protein | \|B4E681_BURCJ | 2.57 | 270 | 0 |
| BCAM0847 | Uncharacterized protein | \|B4EMP9_BURCJ | 2.57 | 139 | 0 |
| BCAM0374 | Uncharacterized protein | \|B4EIW2_BURCJ | 2.55 | 110 | 0 |
| BCAL0820 apt | Adenine phosphoribosyltransferase | \|B4EAT1_BURCJ | 2.54 | 201 | 4 |
| BCAM0062 pca( | 4-carboxymuconolactone decarboxylase | \|B4EGI6_BURCJ | 2.54 | 128 | 4 |
| BCAM1053C | Hypothetical phage protein | \|B4EFR2_BURCJ | 2.53 | 80 | 0 |
| BCAM0271 | Uncharacterized protein | \|B4EI81_BURCJ | 2.53 | 89 | 1 |
| BCAL0163 | Putative phospholipid-binding lipoprotein | \|B4EE88_BURCJ | 2.53 | 273 | 3 |
| BCAL0789 | Uncharacterized protein | \|B4EAQ2_BURCJ | 2.52 | 127 | 0 |
| BCAM0064 | Uncharacterized protein | \|B4EGI8_BURCJ | 2.52 | 82 | 0 |
| BCAL1354 | Uncharacterized protein | \|B4E675_BURCJ | 2.52 | 87 | 0 |
| BCAL1792 | Uncharacterized protein | \|B4E9X3_BURCJ | 2.51 | 97 | 0 |
| BCAL3117 galE | UDP-glucose epimerase | \|B4ECD8_BURCJ | 2.51 | 340 | 6 |
| BCAL2087 ugpQ | Glycerophosphoryl diester phosphodiesterase | \|B4E609_BURCJ | 2.50 | 256 | 6 |
| BCAL3076A | Putative DNA-binding protein | \|B4EC97_BURCJ | 2.49 | 75 | 2 |
| BCAL2044 ldcA | Muramoyltetrapeptide carboxypeptidase | \|B4EC51_BURCJ | 2.49 | 312 | 4 |
| BCAM1108 | Putative monooxygenase | \|B4EG88_BURCJ | 2.49 | 111 | 5 |
| BCAL2444 | Uncharacterized protein | \|B4E6H0_BURCJ | 2.48 | 230 | 0 |
| BCAS0750 | Putative exported protein | \|B4EPD8_BURCJ | 2.46 | 180 | 0 |
| BCAM1491 | Putative exported protein | \|B4EJI2_BURCJ | 2.46 | 109 | 0 |
| BCAL2237 | Putative aminotransferase | \|B4EDW6_BURCJ | 2.45 | 464 | 20 |
| BCAM0345 | Uncharacterized protein | \|B4EIT5_BURCJ | 2.45 | 334 | 0 |
| BCAM0833 | Putative OsmC-like protein | \|B4EMN5_BURCJ | 2.44 | 133 | 0 |
| BCAL1030 | Pirin-like protein | \|B4ECJ2_BURCJ | 2.44 | 288 | 8 |
| BCAL0269 yedY | Sulfoxide reductase catalytic subunit YedY | \|B4E5Z3_BURCJ | 2.43 | 331 | 5 |
| BCAL3033 | Probable outer-membrane lipoproteins carrier protein | \|B4EBR7_BURCJ | 2.42 | 229 | 1 |
| BCAM1481 | Uncharacterized protein | \|B4EJ34_BURCJ | 2.42 | 79 | 0 |
| BCAM1443 | Putative exported protein | \|B4EIZ8_BURCJ | 2.41 | 124 | 0 |
| BCAM1727 | UPF0345 protein BceJ2315_51760 | \|Y5176_BURCJ | 2.41 | 106 | 0 |
| BCAS0662 | Uncharacterized protein | \|B4EP56_BURCJ | 2.40 | 174 | 0 |
| BCAL3043 pgl | 6-phosphogluconolactonase | \|B4EBS7_BURCJ | 2.40 | 226 | 1 |
| BCAL3090 prmB | 50S ribosomal protein L3 glutamine methyltransferase | \|B4ECP4_BURCJ | 2.38 | 302 | 6 |

| | | | | | |
|---|---|---|---|---|---|
| BCAL0815 | OstA-like protein | \|B4EAS6_BURCJ | 2.37 | 229 | 8 |
| BCAM0296 phbI | Acetoacetyl-CoA reductase | \|B4EIA8_BURCJ | **2.37** | 248 | 2 |
| BCAL0077 | Putative oxidoreductase | \|B4EF30_BURCJ | 2.36 | 350 | 7 |
| BCAL0942 | Uncharacterized protein | \|B4EBX2_BURCJ | 2.36 | 94 | 0 |
| BCAL1940 | Uncharacterized protein | \|B4EB26_BURCJ | 2.34 | 208 | 0 |
| BCAL0871 | Uncharacterized protein | \|B4EBC2_BURCJ | 2.34 | 147 | 0 |
| BCAL0332 | Putative stringent starvation protein B | \|B4E6Q4_BURCJ | 2.34 | 174 | 5 |
| BCAL3297 | Putative ferritin DPS-family DNA binding protein | \|B4EE19_BURCJ | 2.33 | 162 | 3 |
| BCAL1494 | Putative monooxygenase | \|B4E7J8_BURCJ | 2.33 | 96 | 6 |
| BCAM1480 | Uncharacterized protein | \|B4EJ33_BURCJ | 2.32 | 96 | 0 |
| BCAM2213 | Uncharacterized protein | \|B4EGE5_BURCJ | 2.32 | 164 | 0 |
| BCAL1473 scoB | Succinyl-CoA:3-ketoacid-coenzyme A transferase subunit B | \|B4E7H8_BURCJ | 2.31 | 213 | 16 |
| BCAL3133 rmlC | dTDP-4-keto-6-deoxy-D-glucose 3,5-epimerase | \|B4ECS4_BURCJ | 2.30 | 183 | 10 |
| BCAS0160 | Uncharacterized protein | \|B4EPS6_BURCJ | 2.30 | 138 | 0 |
| BCAL2360 | Putative sigma factor | \|B4E5N8_BURCJ | 2.28 | 187 | 12 |
| BCAM2209 | Uncharacterized protein | \|B4EGE1_BURCJ | 2.28 | 95 | 0 |
| BCAM2656 | Putative Hsp90 family protein | \|B4EKE1_BURCJ | 2.28 | 161 | 0 |
| BCAL3051 ribE | Riboflavin synthase alpha chain | \|B4EBT5_BURCJ | 2.28 | 209 | 1 |
| BCAL2023 | Putative membrane protein | \|B4EC30_BURCJ | 2.28 | 392 | 0 |
| BCAM1014 | Putative 3-demethylubiquinone-9 3-methyltransferase | \|B4EFM2_BURCJ | 2.26 | 157 | 0 |
| BCAL0734 | Sugar transport PTS system IIa component | \|B4EA59_BURCJ | 2.25 | 156 | 6 |
| BCAL0473 queC | 7-cyano-7-deazaguanine synthase | \|QUEC_BURCJ | 2.24 | 244 | 5 |
| BCAL3025 minE | Cell division topological specificity factor | \|MINE_BURCJ | 2.24 | 84 | 4 |
| BCAM2381 | Putative ABC transport system, exported protein | \|B4EHZ8_BURCJ | 2.24 | 307 | 2 |
| Bcal1909acoC | Dihydrolipoyllysine-residue acetyltransferase component of acetoin cleaving sy | \|B4EAZ5_BURCJ | 2.23 | 371 | 3 |
| BCAM0290 | Putative universal stress protein | \|B4EIA2_BURCJ | 2.22 | 156 | 4 |
| BCAM1965 | Putative hydrolase | \|B4EMU7_BURCJ | 2.22 | 236 | 10 |
| BCAM1496 | Uncharacterized protein | \|B4EJI7_BURCJ | 2.19 | 168 | 0 |
| BCAL0207 hppD | 4-hydroxyphenylpyruvic acid dioxygenase | \|B4E593_BURCJ | 2.18 | 365 | 4 |
| BCAM0876 | Uncharacterized protein | \|B4EMZ4_BURCJ | 2.18 | 211 | 0 |
| BCAM1119 | Uncharacterized protein | \|B4EG99_BURCJ | 2.18 | 135 | 0 |
| BCAL0043 | Putative extracellular ligand-binding protein | \|B4EF03_BURCJ | 2.18 | 379 | 1 |
| BCAM1926 | Uncharacterized protein with direct repeat flanking prophage | \|B4EMQ9_BURCJ | 2.18 | 153 | 11 |

| | | | | | |
|---|---|---|---|---|---|
| BCAL2711 | Peptidyl-prolyl cis-trans isomerase | \|B4E8Y5_BURCJ | 2.17 | 151 | 2 |
| BCAL0396 trpC | Indole-3-glycerol phosphate synthase | \|B4E7A7_BURCJ | 2.17 | 261 | 3 |
| pBCA065 | Uncharacterized protein | \|B4EQK3_BURCJ | 2.16 | 73 | 0 |
| BCAL0322 | Uncharacterized protein | \|B4E645_BURCJ | 2.16 | 121 | 10 |
| BCAL2705 | ABC transporter ATP-binding protein | \|B4E8X9_BURCJ | 2.16 | 238 | 6 |
| BCAM2400 | NAD dependent epimerase/dehydratase family protein | \|B4EI17_BURCJ | 2.16 | 323 | 4 |
| BCAM1291 | L-asparaginase | \|B4EHT1_BURCJ | 2.15 | 332 | 5 |
| BCAM1403 | Putative short chain dehydrogenase | \|B4EIH9_BURCJ | 2.15 | 249 | 3 |
| BCAM0626 | Putative DNA-binding protein | \|B4EL47_BURCJ | 2.15 | 109 | 0 |
| BCAL3189 | Uncharacterized protein | \|B4ECY0_BURCJ | 2.14 | 136 | 0 |
| BCAL0435 | Uncharacterized protein | \|B4E7E3_BURCJ | 2.14 | 184 | 0 |
| BCAL0691 | Putative cytidylyltransferase | \|B4E9P1_BURCJ | 2.14 | 161 | 5 |
| BCAL2324 | Uncharacterized protein | \|B4EER2_BURCJ | 2.14 | 158 | 1 |
| BCAL1070 | Putative redoxin protein | \|B4ED06_BURCJ | 2.14 | 182 | 5 |
| BCAM0316 | Uncharacterized protein | \|B4EIQ7_BURCJ | 2.14 | 157 | 0 |
| BCAL2045 tadA | tRNA-specific adenosine deaminase | \|B4EC52_BURCJ | 2.12 | 198 | 3 |
| BCAM1511 | Uncharacterized protein | \|B4EJK1_BURCJ | 2.12 | 82 | 0 |
| BCAL2328 | Uncharacterized protein | \|B4EER6_BURCJ | 2.11 | 140 | 0 |
| BCAM1502 | Uncharacterized protein | \|B4EJJ2_BURCJ | 2.11 | 84 | 0 |
| BCAL1864 | Uncharacterized protein | \|B4EAG3_BURCJ | 2.11 | 279 | 0 |
| BCAM0528 | Putative oxidoreductase/short-chain dehydrogenase | \|B4EKH1_BURCJ | 2.10 | 234 | 8 |
| BCAL1088 | Putative exported protein | \|B4ED24_BURCJ | 2.10 | 213 | 0 |
| BCAL0817 kdsC | Putative 3-deoxy-D-manno-octulosonate 8-phosphate phosphatase | \|B4EAS8_BURCJ | 2.10 | 178 | 4 |
| BCAL3319 | Uncharacterized protein | \|B4EE40_BURCJ | 2.10 | 154 | 0 |
| pBCA09 | Uncharacterized protein | \|B4EQM6_BURCJ | 2.09 | 109 | 0 |
| BCAL3456 | Putative thioredoxin reductase | \|B4E5Y6_BURCJ | 2.09 | 168 | 24 |
| BCAM2609 | Putative exported protein | \|B4EK95_BURCJ | 2.08 | 172 | 0 |
| BCAL3310 | Putative exported protein | \|B4EE31_BURCJ | 2.08 | 193 | 1 |
| BCAL2771 | Cys-tRNA(Pro)/Cys-tRNA(Cys) deacylase | \|B4E9H0_BURCJ | 2.08 | 163 | 3 |
| BCAS0003 | ParA protein | \|B4EQD1_BURCJ | 2.08 | 231 | 1 |
| BCAL1630 | Uncharacterized protein | \|B4E8Q3_BURCJ | 2.07 | 247 | 0 |
| BCAM0900 | Putative exported protein | \|B4EN18_BURCJ | 2.07 | 131 | 0 |
| BCAM2598 | Putative short-chain dehydrogenase | \|B4EJU7_BURCJ | 2.07 | 291 | 2 |

| | | | | | |
|---|---|---|---|---|---|
| BCAL2694 | Putative dehydrogenase | \|B4E8W7_BURCJ | 2.06 | 257 | 3 |
| BCAL1493 | Putative exported protein | \|B4E7J7_BURCJ | 2.06 | 402 | 5 |
| BCAL0349 | Putative outer membrane protein | \|B4E6S1_BURCJ | 2.06 | 316 | 2 |
| BCAL0122 | Histone-like nucleoid-structuring (H-NS) protein | \|B4EDJ6_BURCJ | 2.06 | 98 | 0 |
| BCAL2689 | Putative short-chain type dehydrogenase/reductase | \|B4E8W2_BURCJ | 2.05 | 269 | 6 |
| BCAL0794 | Uncharacterized protein | \|B4EAQ7_BURCJ | 2.05 | 204 | 3 |
| BCAM0015 | Uncharacterized protein | \|B4EG01_BURCJ | 2.05 | 159 | 0 |
| BCAL2828 | Putative exported protein | \|B4EA12_BURCJ | 2.05 | 154 | 0 |
| BCAL0394 ung | Uracil-DNA glycosylase | \|B4E6W7_BURCJ | 2.04 | 286 | 3 |
| BCAL0037 atpC | ATP synthase epsilon chain | \|ATPE_BURCJ | 2.04 | 141 | 10 |
| BCAM1743 | Periplasmic solute-binding protein | \|B4ELC1_BURCJ | 2.04 | 298 | 2 |
| BCAM2238 | Putative short-chain dehydrogenase/reductase | \|B4EGG9_BURCJ | 2.03 | 249 | 9 |
| BCAM0979 | Putative glutathione-S-transferase | \|B4EFC4_BURCJ | 2.03 | 241 | 55 |
| BCAL1962 | Putative deoxyribonuclease | \|B4EBJ0_BURCJ | 2.03 | 259 | 1 |
| BCAM2652 | TetR family regulatory protein | \|B4EKD7_BURCJ | 2.03 | 194 | 3 |
| BCAM0276 | **Universal stress protein** | \|B4EI87_BURCJ | 2.02 | 156 | 3 |
| BCAL2304 | Putative D-beta-hydroxybutyrate dehydrogenase | \|B4EEP3_BURCJ | 2.02 | 262 | 3 |
| BCAL0802 ispE | 4-diphosphocytidyl-2-C-methyl-D-erythritol kinase | \|ISPE_BURCJ | 2.02 | 293 | 7 |
| BCAM0286 | Putative alcohol dehydrogenase | \|B4EI98_BURCJ | 2.01 | 329 | 6 |
| BCAL0009 | Putative pterin-4-alpha-carbinolamine dehydratase | \|PHS_BURCJ | 2.01 | 102 | 2 |
| BCAM 1365 hpa | Putative homoprotocatechuate degradative operon repressor | \|B4EIE1_BURCJ | 2.00 | 146 | 9 |
| BCAM0943 | Uncharacterized protein | \|B4EF88_BURCJ | 2.00 | 106 | 0 |
| BCAL2242 hutI | Imidazolonepropionase | \|HUTI_BURCJ | 2.00 | 407 | 5 |
| BCAM1692 | Putative fumarylpyruvate hydrolase | \|B4EKU2_BURCJ | 1.99 | 232 | 3 |
| BCAM1363 | Uncharacterized protein | \|B4EID9_BURCJ | 1.99 | 115 | 0 |
| BCAL3338 ruvA | Holliday junction ATP-dependent DNA helicase RuvA | \|RUVA_BURCJ | 1.99 | 193 | 11 |
| BCAM0026 | Putative siderophore-interacting protein | \|B4EG12_BURCJ | 1.98 | 273 | 3 |
| BCAL1665 | SpoVR like protein | \|B4E8T8_BURCJ | 1.98 | 507 | 1 |
| BCAL1799 | Uncharacterized protein | \|B4E9Y0_BURCJ | 1.98 | 143 | 0 |
| BCAL1652 sbp | Sulfate-binding protein | \|B4E8S5_BURCJ | 1.98 | 345 | 7 |
| BCAL3349 | Putative OsmC-like protein | \|B4EET8_BURCJ | 1.98 | 154 | 1 |
| BCAL2827 | Uncharacterized protein | \|B4EA11_BURCJ | 1.98 | 139 | 0 |
| BCAS0693 | Putative monooxygenase | \|B4EP85_BURCJ | 1.98 | 95 | 5 |

| | | | | | |
|---|---|---|---|---|---|
| BCAL3272 grpE | Protein GrpE | \|GRPE_BURCJ | 1.97 | 181 | 6 |
| BCAL2325 | Uncharacterized protein | \|B4EER3_BURCJ | 1.97 | 153 | 9 |
| BCAL2434 | Glyoxalase/bleomycin resistance protein/dioxygenase superfamily protein | \|B4E6G0_BURCJ | 1.97 | 151 | 0 |
| BCAL3009greB | Transcription elongation factor GreB | \|B4EBP3_BURCJ | 1.97 | 187 | 4 |
| BCAL3366 eda | KHG/KDPG aldolase | \|B4EEV4_BURCJ | 1.97 | 208 | 8 |
| BCAL3198 | Putative short-chain dehydrogenase | \|B4ECZ0_BURCJ | 1.97 | 248 | 5 |
| BCAL2185 lolD | Lipoprotein releasing system ATP-binding protein | \|B4EDA7_BURCJ | 1.97 | 231 | 7 |
| BCAL1299 | Uncharacterized protein | \|B4E5I4_BURCJ | 1.96 | 148 | 0 |
| BCAM0002 | ArsC family arsenate reductase | \|B4EFY7_BURCJ | 1.96 | 118 | 1 |
| BCAL0405 paaI | Phenylacetic acid degradation protein PaaI | \|B4E7B6_BURCJ | 1.96 | 150 | 1 |
| BCAM1351 | Putative regulatory protein | \|B4EIC7_BURCJ | 1.95 | 130 | 5 |
| BCAM2383 | ABC transporter ATP-binding protein | \|B4EI00_BURCJ | 1.95 | 317 | 5 |
| BCAL0053 | PadR family regulatory protein | \|B4EF06_BURCJ | 1.95 | 190 | 0 |
| BCAL1837 | Uncharacterized protein | \|B4EAD6_BURCJ | 1.95 | 229 | 5 |
| BCAL2307 | Uncharacterized protein | \|B4EEP6_BURCJ | 1.95 | 122 | 0 |
| BCAM0052 | Uncharacterized protein | \|B4EG38_BURCJ | 1.94 | 77 | 0 |
| BCAM2000 | Uncharacterized protein | \|B4EN44_BURCJ | 1.93 | 246 | 3 |
| BCAL0326 | Serine peptidase, subfamily S1B | \|B4E649_BURCJ | 1.93 | 401 | 5 |
| BCAL3382 | GntR family regulatory protein | \|B4EEX0_BURCJ | 1.93 | 240 | 3 |
| BCAL0156 | AnsC family regulatory protein | \|B4EE81_BURCJ | 1.93 | 156 | 3 |
| BCAL1971 | Uncharacterized protein | \|B4EBJ9_BURCJ | 1.93 | 104 | 0 |
| BCAL0060 eutC | Ethanolamine ammonia-lyase light chain | \|B4EF13_BURCJ | 1.93 | 265 | 2 |
| BCAL1651 lexA | LexA repressor | \|LEXA_BURCJ | 1.93 | 215 | 11 |
| BCAL0604 | Putative cheavy metal binding protein | \|B4E927_BURCJ | 1.92 | 66 | 12 |
| BCAL0389 dsbC | Thiol:disulfide interchange protein DsbC | \|B4E6W2_BURCJ | 1.92 | 242 | 7 |
| BCAL3398 | Putative competence-damaged related protein | \|B4E5S8_BURCJ | 1.91 | 166 | 3 |
| BCAL3012 gmk | Guanylate kinase | \|B4EBP6_BURCJ | 1.91 | 227 | 6 |
| BCAS0257 | Putative acetyltransferase | \|B4EQ25_BURCJ | 1.91 | 154 | 9 |
| BCAL0074 gcvH | Glycine cleavage system H protein | \|GCSH_BURCJ | 1.91 | 126 | 4 |
| BCAL2976 fdsG | NAD-dependent formate dehydrogenase gamma subunit | \|B4EB75_BURCJ | 1.90 | 166 | 3 |
| BCAL1297 | Uncharacterized protein | \|B4E5I2_BURCJ | 1.90 | 265 | 6 |
| BCAL0402 apaG | Protein ApaG | \|B4E7B3_BURCJ | 1.90 | 124 | 0 |
| BCAL2240 | HutG protein | \|B4EDW9_BURCJ | 1.90 | 265 | 0 |

| | | | | | |
|---|---|---|---|---|---|
| BCAL0502 dksA | RNA polymerase-binding transcription factor DksA | \|B4E7Z3_BURCJ | 1.90 | 138 | 2 |
| BCAL0305 | Putative exported protein | \|B4E628_BURCJ | 1.89 | 209 | 5 |
| BCAL2095 def | Peptide deformylase | \|B4ECN5_BURCJ | 1.88 | 177 | 6 |
| BCAS0118 | Putative H-NS family DNA-binding protein | \|B4EPN5_BURCJ | 1.88 | 109 | 0 |
| BCAL0008A | Uncharacterized protein | \|B4E565_BURCJ | 1.88 | 76 | 0 |
| BCAM1873 | Glyoxalase/bleomycin resistance/dioxygenase superfamily protein | \|B4EMD5_BURCJ | 1.87 | 128 | 0 |
| BCAL3184 hmgA | Homogentisate 1,2-dioxygenase | \|HGD_BURCJ | 1.87 | 444 | 5 |
| BCAL2296 | Uncharacterized protein | \|B4EEN5_BURCJ | 1.87 | 180 | 0 |
| BCAM2663 | DsbA-like thioredoxin protein | \|B4EKE8_BURCJ | 1.87 | 212 | 17 |
| BCAL0343 bcsL | Putative type VI secretion system protein TssD | \|B4E6R5_BURCJ | 1.87 | 167 | 1 |
| BCAM2594 | Putative alcohol dehydrogenase | \|B4EJU3_BURCJ | 1.87 | 327 | 2 |
| BCAL0476 | HpcH/HpaI aldolase/citrate lyase family protein | \|B4E7W8_BURCJ | 1.87 | 261 | 1 |
| BCAM1818 | Uncharacterized protein | \|B4ELX8_BURCJ | 1.87 | 187 | 0 |
| BCAS0013 | Putative molybdenum transport protein | \|B4EQE1_BURCJ | 1.86 | 141 | 3 |
| BCAL0273 cyaY | Protein CyaY | \|CYAY_BURCJ | 1.86 | 108 | 2 |
| BCAL0765 | Putative exported protein | \|B4EA90_BURCJ | 1.86 | 321 | 0 |
| BCAL3413 aroE | Shikimate dehydrogenase | \|B4E5U3_BURCJ | 1.86 | 288 | 3 |
| BCAM0907 | Sulfurtransferase | \|B4EN25_BURCJ | 1.86 | 289 | 2 |
| BCAL1071 | NAD dependent epimerase/dehydratase family protein | \|B4ED07_BURCJ | 1.86 | 209 | 1 |
| BCAM0050 | Universal stress protein | \|B4EG36_BURCJ | 1.85 | 155 | 7 |
| BCAM1311 ami( | Putative aliphatic amidase expression-regulating protein | \|B4EHV1_BURCJ | 1.85 | 392 | 4 |
| BCAL0814 | ABC transporter ATP-binding protein | \|B4EAS5_BURCJ | 1.85 | 258 | 7 |
| BCAL2466 eco | Ecotin | \|B4E732_BURCJ | 1.85 | 153 | 2 |
| BCAL3199 | Putative thioesterase | \|B4EDB6_BURCJ | 1.85 | 161 | 2 |
| BCAL2102 | ArsC family protein | \|B4ECP2_BURCJ | 1.85 | 122 | 7 |
| BCAL1991 | ABC transporter ATP-binding protein | \|B4EBL8_BURCJ | 1.85 | 237 | 1 |
| BCAM0526 | Pyridoxamine 5-phosphate oxidase family protein | \|B4EKG9_BURCJ | 1.85 | 195 | 8 |
| BCAS0407 | Uncharacterized protein | \|B4ENF6_BURCJ | 1.84 | 104 | 0 |
| BCAL2206 phaP | Phasin-like protein | \|B4EDT5_BURCJ | 1.84 | 188 | 0 |
| BCAS0225 | LysR family regulatory protein | \|B4EPZ1_BURCJ | 1.84 | 330 | 2 |
| BCAL1472 scoA | Succinyl-CoA:3-ketoacid-coenzyme A transferase subunit A | \|B4E7H7_BURCJ | 1.84 | 234 | 16 |
| BCAL2780 | Putative thioredoxin protein | \|B4E9H9_BURCJ | 1.84 | 282 | 3 |
| BCAL0470 | Putative exported protein | \|B4E7W2_BURCJ | 1.84 | 196 | 0 |

| | | | | | |
|---|---|---|---|---|---|
| BCAM0587 | Uncharacterized protein | \|B4EL08_BURCJ | 1.84 | 253 | 10 |
| BCAM0848 | Glyoxalase/bleomycin resistance protein/dioxygenase superfamily protein | \|B4EMQ0_BURCJ | 1.84 | 138 | 0 |
| BCAL0677 dsbA | Thiol:disulfide interchange protein | \|B4E9M7_BURCJ | 1.84 | 212 | 4 |
| BCAM2830 | Uncharacterized protein | \|B4EM08_BURCJ | 1.84 | 235 | 0 |
| BCAL0149 | Putative dienelactone hydrolase | \|B4EE74_BURCJ | 1.84 | 230 | 6 |
| BCAL2766 | Uncharacterized protein | \|B4E9G5_BURCJ | 1.83 | 110 | 0 |
| BCAL0474 queE | 7-carboxy-7-deazaguanine synthase | \|B4E7W6_BURCJ | 1.83 | 210 | 2 |
| BCAL1920 | Putative DNA-binding protein | \|B4EB06_BURCJ | 1.83 | 172 | 1 |
| BCAL0465 def | Peptide deformylase | \|B4E7V7_BURCJ | 1.83 | 167 | 4 |
| BCAS0539 | Cro/cI repressor transcription regulator | \|B4ENT6_BURCJ | 1.82 | 138 | 0 |
| BCAM1211 | Cysteine peptidase/transferase, family C45 | \|B4EH91_BURCJ | 1.82 | 349 | 0 |
| BCAL3147 groS | 10 kDa chaperonin | \|B4ECT8_BURCJ | 1.82 | 97 | 3 |
| BCAM2633 | Streptomycin 3-kinase | \|B4EKB8_BURCJ | 1.82 | 265 | 1 |
| BCAL0989 | Uncharacterized protein | \|B4EAI9_BURCJ | 1.82 | 211 | 0 |
| BCAL0550 | LamB/YcsF family protein | \|B4E8J3_BURCJ | 1.82 | 272 | 2 |
| BCAL1942 rplI | 50S ribosomal protein L9 | \|RL9_BURCJ | 1.81 | 150 | 5 |
| BCAL2232 | Probable Fe(2+)-trafficking protein | \|FETP_BURCJ | 1.81 | 91 | 1 |
| BCAL2464 | Short chain dehydrogenase | \|B4E730_BURCJ | 1.81 | 246 | 11 |
| BCAL2211 | Two-component regulatory system, response regulator protein | \|B4EDU0_BURCJ | 1.81 | 212 | 2 |
| BCAL1861 phbB | Acetoacetyl-CoA reductase | \|B4EAG0_BURCJ | **1.81** | 246 | 8 |
| BCAM2509 | Putative FucU/RbsD family transport protein | \|B4EJ62_BURCJ | 1.81 | 151 | 3 |
| BCAL1898 | Nucleoid-associated protein BceJ2315_18610 | \|Y1861_BURCJ | 1.80 | 108 | 4 |
| BCAL1859 | Uncharacterized protein | \|B4EAF8_BURCJ | 1.80 | 177 | 1 |
| BCAL0221 nusG | Transcription termination/antitermination protein NusG | \|B4E5A7_BURCJ | 1.79 | 185 | 3 |
| BCAL2913 fdx | 2Fe-2S ferredoxin | \|B4EDB5_BURCJ | 1.79 | 113 | 4 |
| BCAL1441 | SirA-like protein | \|B4E707_BURCJ | 1.79 | 75 | 3 |
| BCAL2067 | Uncharacterized protein | \|B4EC74_BURCJ | 1.79 | 238 | 0 |
| BCAL3434 | Putative hypoxanthine phosphoribosyltransferase | \|B4E5W4_BURCJ | 1.79 | 192 | 4 |
| BCAL3378 fur | Ferric uptake regulator | \|B4EEW6_BURCJ | 1.79 | 142 | 8 |
| BCAL2153 ppiB | Peptidyl-prolyl cis-trans isomerase | \|B4ED76_BURCJ | 1.78 | 163 | 11 |
| BCAL2175 | Uncharacterized protein | \|B4ED97_BURCJ | 1.78 | 174 | 14 |
| BCAL1280 | Putative hydrolase | \|B4E5G7_BURCJ | 1.78 | 153 | 1 |
| BCAL0012 | Putative adenylate cyclase | \|B4E569_BURCJ | 1.78 | 177 | 0 |

| | | | | | |
|---|---|---|---|---|---|
| BCAM2278 | GntR family regulatory protein | \|B4EGZ4_BURCJ | 1.78 | 255 | 6 |
| BCAM0023 adc | Probable acetoacetate decarboxylase | \|ADC_BURCJ | 1.78 | 246 | 2 |
| BCAM2742 | Uncharacterized protein | \|B4ELE1_BURCJ | 1.77 | 117 | 4 |
| BCAM2153 | Putative cytochrome P450 oxidoreductase | \|B4EFV0_BURCJ | 1.77 | 386 | 2 |
| BCAL1520 | Putative lipoprotein | \|B4E7M4_BURCJ | 1.77 | 153 | 0 |
| BCAL2956 | Putative exported protein | \|B4EB46_BURCJ | 1.77 | 199 | 0 |
| BCAS0281 | Putative 2-hydroxy-3-oxopropionate reductase | \|B4EQ47_BURCJ | 1.77 | 300 | 9 |
| BCAM1411 | Putative short-chain dehydrogenase | \|B4EII7_BURCJ | 1.76 | 264 | 1 |
| BCAM2644 | Putative glutathione S-transferase | \|B4EKC9_BURCJ | 1.76 | 223 | 2 |
| BCAL1933 | L-arabinose formyltransferase | \|B4EB19_BURCJ | 1.76 | 315 | 3 |
| BCAL3093 | Uncharacterized protein | \|B4ECB4_BURCJ | 1.76 | 105 | 3 |
| BCAL2790 kynB | Kynurenine formamidase | \|KYNB_BURCJ | 1.76 | 213 | 4 |
| BCAL1026 | Uncharacterized protein | \|B4EA30_BURCJ | 1.75 | 167 | 2 |
| BCAL0892 | Putative nucleotidyl transferase | \|B4EBE3_BURCJ | 1.75 | 240 | 12 |
| BCAL0317 hisA | 1-(5-phosphoribosyl)-5-[(5-phosphoribosylamino)methylideneamino] imidazol | \|HIS4_BURCJ | 1.75 | 251 | 4 |
| **BCAM1217 ahp** | **Alkyl hydroperoxide reductase subunit C** | \|B4EH97_BURCJ | 1.75 | 187 | 20 |
| BCAL2106gpo | Glutathione peroxidase | \|B4ECP6_BURCJ | 1.75 | 159 | 3 |
| BCAM2449 | Putative metallophosphoesterase protein | \|B4EIL5_BURCJ | 1.75 | 244 | 2 |
| BCAM0294 | Putative universal stress protein | \|B4EIA6_BURCJ | 1.75 | 279 | 0 |
| BCAL1503 | Putative transcriptional regulator protein | \|B4E7K7_BURCJ | 1.74 | 345 | 3 |
| BCAL2392 rsfS | Ribosomal silencing factor RsfS | \|B4E5S0_BURCJ | 1.74 | 147 | 7 |
| BCAL0984 | Haloacid dehalogenase-like hydrolase | \|B4EAJ4_BURCJ | 1.74 | 219 | 9 |
| BCAL1751 | Glyoxalase/bleomycin resistance protein/dioxygenase superfamily protein | \|B4E9T2_BURCJ | 1.74 | 127 | 7 |
| BCAL1992 | Putative acyl-CoA thioesterase | \|B4EBL9_BURCJ | 1.74 | 222 | 6 |
| BCAM2688 | Putative isomerase | \|B4EKV7_BURCJ | 1.74 | 369 | 4 |
| BCAL1900 trxA | Thioredoxin | \|B4EAY5_BURCJ | 1.74 | 108 | 12 |
| BCAL3145 | Putative kinase | \|B4ECT6_BURCJ | 1.73 | 288 | 1 |
| BCAM1192 | Putative carboxylesterase | \|B4EGU7_BURCJ | 1.73 | 227 | 6 |
| BCAM0906 | Putative dienelactone hydrolase family protein | \|B4EN24_BURCJ | 1.73 | 291 | 1 |
| BCAM1801 | Hypothetical phage protein | \|B4ELW1_BURCJ | 1.73 | 69 | 0 |
| BCAL0878 | Uncharacterized protein | \|B4EBC9_BURCJ | 1.73 | 117 | 0 |
| BCAL3407 | Putative aldolase | \|B4E5T7_BURCJ | 1.73 | 225 | 4 |
| BCAL2870 mucB | Sigma-E factor regulatory protein RseB 1 | \|B4EAH8_BURCJ | 1.72 | 350 | 7 |

| | | | | | |
|---|---|---|---|---|---|
| BCAL0795 coaD | Phosphopantetheine adenylyltransferase | \|COAD_BURCJ | 1.72 | 165 | 6 |
| BCAL2616 glnB2 | Nitrogen regulatory protein P-II 2 | \|B4E8B0_BURCJ | 1.72 | 112 | 7 |
| BCAL0215 paaB | Phenylacetic acid degradation protein PaaB | \|B4E5A1_BURCJ | 1.72 | 94 | 1 |
| BCAL0939 | Uncharacterized protein | \|B4EBW9_BURCJ | 1.71 | 115 | 7 |
| BCAL0422 dnaN | DNA polymerase III subunit beta | \|B4E7D0_BURCJ | 1.71 | 368 | 6 |
| BCAL0706 | Uncharacterized protein | \|B4E9Q6_BURCJ | 1.71 | 102 | 0 |
| BCAL2651 panC | Pantothenate synthetase | \|PANC_BURCJ | 1.71 | 279 | 5 |
| BCAM1538 | Putative dehydrogenase, monooxygenase subunit | \|B4EJM8_BURCJ | 1.71 | 99 | 6 |
| BCAM0001 | Putative sigma factor | \|B3KYE4_BURCJ | 1.71 | 199 | 6 |
| BCAL1999 | MarR family regulatory protein | \|B4EBM6_BURCJ | 1.71 | 149 | 11 |
| BCAM0291 | Putative universal stress protein | \|B4EIA3_BURCJ | 1.71 | 277 | 0 |
| BCAL1869 | Putative exported protein | \|B4EAG8_BURCJ | 1.71 | 224 | 0 |
| BCAL0468 htpX | Protease HtpX homolog | \|HTPX_BURCJ | 1.71 | 285 | 6 |
| BCAL2706 | ABC transporter ATP-binding protein | \|B4E8Y0_BURCJ | 1.71 | 258 | 6 |
| BCAL0962 moaC | Cyclic pyranopterin monophosphate synthase accessory protein | \|MOAC_BURCJ | 1.70 | 162 | 2 |
| BCAL0970 | CreA protein 1 | \|B4EAK8_BURCJ | 1.70 | 161 | 0 |
| BCAS0735 | Metallo peptidase, family M20 unassigned | \|B4EPC3_BURCJ | 1.70 | 426 | 5 |
| BCAL1887 ndk | Nucleoside diphosphate kinase | \|NDK_BURCJ | 1.70 | 141 | 6 |
| BCAM2160 | Two-component regulatory system, response regulator protein | \|B4EFV7_BURCJ | 1.70 | 220 | 9 |
| BCAM0140 | Putative short-chain dehydrogenase | \|B4EH40_BURCJ | 1.70 | 260 | 16 |
| BCAL1936 | AhpC/TSA family protein | \|B4EB22_BURCJ | 1.70 | 153 | 9 |
| BCAM2599 | TetR family regulatory protein | \|B4EJU8_BURCJ | 1.70 | 192 | 0 |
| BCAM0869 fbp | Peptidyl-prolyl cis-trans isomerase | \|B4EMY8_BURCJ | 1.70 | 113 | 6 |
| BCAL1257 gloB | Hydroxyacylglutathione hydrolase | \|B4EEJ4_BURCJ | 1.70 | 226 | 5 |
| BCAM0147 | Uncharacterized protein | \|B4EH47_BURCJ | 1.70 | 252 | 0 |
| BCAL3291 | Uncharacterized protein | \|B4EE13_BURCJ | 1.70 | 232 | 1 |
| BCAL0670 | Putative short chain dehydrogenase | \|B4E9M0_BURCJ | 1.70 | 225 | 5 |
| BCAL2906 | Uncharacterized protein | \|B4EAL4_BURCJ | 1.70 | 72 | 1 |
| BCAM0797 | Uncharacterized protein | \|B4EMJ9_BURCJ | 1.69 | 149 | 5 |
| BCAL2973 | Putative exported protein | \|B4EB72_BURCJ | 1.69 | 117 | 0 |
| BCAL0812 | Sigma-54 modulation protein | \|B4EAS3_BURCJ | 1.69 | 119 | 4 |
| BCAL0506 | Uncharacterized protein | \|B4E7Z7_BURCJ | 1.69 | 249 | 0 |
| BCAL0367 | Putative chaperone protein | \|B4E6T9_BURCJ | 1.69 | 416 | 5 |

| | | | | | |
|---|---|---|---|---|---|
| BCAL3026 minD | \|B4EBR0_BURCJ Site-determining protein | \|B4EBR0_BURCJ | 1.69 | 266 | 7 |
| BCAS0638 groS | 10 kDa chaperonin | \|B4EP32_BURCJ | 1.69 | 105 | 3 |
| BCAL0732 gshB | Glutathione synthetase | \|B4EA58_BURCJ | 1.69 | 318 | 3 |
| pBCA054 | LuxR family regulatory protein | \|B4EQJ3_BURCJ | 1.69 | 342 | 0 |
| BCAL3472 coq7 | 2-nonaprenyl-3-methyl-6-methoxy-1,4-benzoquinol hydroxylase | \|COQ7_BURCJ | 1.69 | 208 | 6 |
| BCAL0069 | Uncharacterized protein | \|B4EF22_BURCJ | 1.69 | 153 | 0 |
| BCAM2431 | Putative enoyl coenzyme A hydratase-like protein | \|B4EIJ7_BURCJ | 1.68 | 261 | 3 |
| BCAL3163 | Putative nucleotidyltransferase | \|B4ECV4_BURCJ | 1.68 | 196 | 0 |
| BCAL2039 | Putative uricase | \|B4EC46_BURCJ | 1.68 | 173 | 2 |
| BCAL0440 | Putative exported protein | \|B4E7E8_BURCJ | 1.68 | 105 | 0 |
| BCAL1498 | Uncharacterized protein | \|B4E7K2_BURCJ | 1.68 | 105 | 0 |
| BCAL2762 adk | Adenylate kinase | \|KAD_BURCJ | 1.68 | 220 | 6 |
| BCAM2450 | Putative hydrolase | \|B4EIL6_BURCJ | 1.68 | 203 | 18 |
| BCAL2622 ppa | Inorganic pyrophosphatase | \|B4E8B6_BURCJ | 1.68 | 175 | 6 |
| BCAL1465 | Uncharacterized protein | \|B4E7H0_BURCJ | 1.67 | 151 | 6 |
| BCAL2743 | Putative aldo/keto reductase family oxidoreductase | \|B4E9E2_BURCJ | 1.67 | 281 | 3 |
| BCAL2114 | Uracil DNA glycosylase superfamily protein | \|B4ECQ4_BURCJ | 1.67 | 343 | 1 |
| BCAL2915 dfrA | Dihydrofolate reductase | \|B4EAM3_BURCJ | 1.67 | 166 | 4 |
| BCAL1982 msrB | Peptide methionine sulfoxide reductase MsrB | \|MSRB_BURCJ | 1.67 | 143 | 4 |
| BCAL2816 | S-formylglutathione hydrolase | \|B4EA00_BURCJ | 1.67 | 282 | 8 |
| BCAL3004 | Chorismate mutase | \|B4EBN8_BURCJ | 1.67 | 197 | 3 |
| BCAL0995 acpP1 | Acyl carrier protein | \|B4EAI3_BURCJ | 1.66 | 79 | 5 |
| BCAM2461 | Putative inosine-uridine preferring nucleoside hydrolase | \|B4EIM7_BURCJ | 1.66 | 350 | 1 |
| BCAM2073 | Putative exported protein | \|B4EFE1_BURCJ | 1.66 | 192 | 0 |
| BCAM1139 | MarR family regulatory protein | \|B4EGB9_BURCJ | 1.66 | 156 | 4 |
| BCAL1250 | Putative glutathione S-transferase | \|B4EEI7_BURCJ | 1.66 | 209 | 34 |
| BCAL0745trmL | tRNA (cytidine(34)-2-O)-methyltransferase | \|B4EA70_BURCJ | 1.66 | 156 | 3 |
| BCAL2826a | Putative membrane protein | \|B4EA10_BURCJ | 1.65 | 235 | 0 |
| BCAL2934 etfA | Electron transfer flavoprotein alpha-subunit | \|B4EAP2_BURCJ | 1.65 | 311 | 6 |
| BCAL2940 | Putative histone deacetylase-family protein | \|B4EB30_BURCJ | 1.65 | 307 | 17 |
| BCAL0316 hisH | Imidazole glycerol phosphate synthase subunit HisH | \|B4E639_BURCJ | 1.65 | 213 | 5 |
| BCAL0120 | Haloacid dehalogenase-like hydrolase | \|B4EDJ4_BURCJ | 1.65 | 274 | 3 |
| BCAM0765 | Aldose epimerase family protein | \|B4EM67_BURCJ | 1.65 | 309 | 10 |

| | | | | | |
|---|---|---|---|---|---|
| BCAL2932 lrp | Leucine-responsive regulatory protein | \|B4EAP0_BURCJ | 1.64 | 162 | 7 |
| BCAL0516 | Uncharacterized protein | \|B4E807_BURCJ | 1.64 | 70 | 0 |
| BCAM1019 fdnH | Formate dehydrogenase, iron-sulfur subunit | \|B4EFM6_BURCJ | 1.64 | 304 | 6 |
| BCAL0557 | Putative glutathione S-transferase | \|B4E8K0_BURCJ | 1.64 | 215 | 2 |
| BCAL3192 | Putative oxidoreductase | \|B4ECY3_BURCJ | 1.64 | 212 | 25 |
| BCAM0990 trpF | N-(5-phosphoribosyl)anthranilate isomerase | \|B4EFJ9_BURCJ | 1.63 | 233 | 3 |
| BCAL2731 clpS | ATP-dependent Clp protease adapter protein ClpS | \|CLPS_BURCJ | 1.63 | 104 | 2 |
| BCAM0884 | Two-component regulatory system, sensor kinase protein | \|B4EN02_BURCJ | 1.63 | 259 | 27 |
| BCAL0341 Tssb | Putative type VI secretion system protein TssB | \|B4E6R3_BURCJ | 1.63 | 171 | 0 |
| BCAM0682 | Putative oxidoreductase | \|B4ELN3_BURCJ | 1.63 | 329 | 8 |
| BCAL0968 | Uncharacterized protein | \|B4EBZ8_BURCJ | 1.63 | 64 | 0 |
| BCAL2630 hemC | Porphobilinogen deaminase | \|HEM3_BURCJ | 1.63 | 334 | 4 |
| BCAL1459 | Calcineurin-like phosphoesterase | \|B4E725_BURCJ | 1.63 | 276 | 1 |
| BCAL1922 moaE | Molybdopterin converting factor subunit 2 | \|B4EB08_BURCJ | 1.62 | 158 | 3 |
| BCAM0322 | Two-component regulatory system, response regulator protein | \|B4EIR3_BURCJ | 1.62 | 220 | 8 |
| BCAL2037 allA | Ureidoglycolate lyase | \|B4EC44_BURCJ | 1.62 | 170 | 3 |
| BCAL0517 | phosphotransferase enzyme family protein | \|B4E808_BURCJ | 1.62 | 340 | 4 |
| BCAL0971 | 4Fe-4S ferredoxin | \|B4EAK7_BURCJ | 1.62 | 107 | 3 |
| BCAM0191 | Putative non-ribosomal peptide synthetase | \|B4EHM3_BURCJ | 1.61 | 453 | 3 |
| BCAL0409 paaF | Putative phenylacetic acid degradation enoyl-CoA hydratase PaaF | \|B4E7C0_BURCJ | 1.61 | 258 | 11 |
| BCAL2372 | Putative alanyl-tRNA synthetase | \|B4E5Q0_BURCJ | 1.61 | 243 | 1 |
| BCAL0988 | Maf-like protein BCAL0988 | \|B4EAJ0_BURCJ | 1.60 | 210 | 1 |
| BCAM2476 nadE | NH(3)-dependent NAD(+) synthetase | \|NADE_BURCJ | 1.60 | 282 | 5 |
| BCAM2642 | Uncharacterized protein | \|B4EKC7_BURCJ | 1.60 | 139 | 2 |
| BCAL2667 | Cell division protein ZapA | \|B4E8G0_BURCJ | 1.60 | 104 | 0 |
| BCAL3096 pcm | Protein-L-isoaspartate O-methyltransferase | \|B4ECB7_BURCJ | 1.60 | 218 | 7 |
| BCAL2668 | Uncharacterized protein | \|B4E8G1_BURCJ | 1.60 | 153 | 1 |
| BCAM1536 | TetR family regulatory protein | \|B4EJM6_BURCJ | 1.60 | 206 | 5 |
| BCAL2779 | Pirin-like protein | \|B4E9H8_BURCJ | 1.59 | 293 | 7 |
| BCAL2695 | Uncharacterized protein | \|B4E8W8_BURCJ | 1.59 | 166 | 4 |
| BCAL1505 rimP | Ribosome maturation factor RimP | \|RIMP_BURCJ | 1.59 | 152 | 2 |
| BCAL2030 | 5-hydroxyisourate hydrolase | \|B4EC37_BURCJ | 1.59 | 117 | 3 |
| BCAL1725 cobH | Precorrin-8X methylmutase | \|B4E9C4_BURCJ | 1.59 | 208 | 3 |

| | | | | | |
|---|---|---|---|---|---|
| BCAS0598 deoC | Deoxyribose-phosphate aldolase | \|DEOC_BURCJ | 1.59 | 226 | 6 |
| BCAL2013 | AhpC/TSA family protein | \|B4EC20_BURCJ | 1.59 | 182 | 11 |
| BCAL2229 | Putative exported protein | \|B4EDV8_BURCJ | 1.59 | 329 | 0 |
| BCAL0896 pdxA | 4-hydroxythreonine-4-phosphate dehydrogenase | \|B4EBE7_BURCJ | 1.58 | 329 | 6 |
| BCAL3140 pyrR | Bifunctional regulator/uracil phosphoribosyltransferase | \|B4ECT1_BURCJ | 1.57 | 174 | 2 |
| BCAL1425 | Putative glucose 1-dehydrogenase | \|B4E6Z1_BURCJ | 1.57 | 248 | 7 |
| BCAL1595 | Putative DNA-binding phage protein | \|B4E891_BURCJ | 1.57 | 156 | 4 |
| BCAL0728 | Uncharacterized protein | \|B4EA54_BURCJ | 1.57 | 83 | 0 |
| BCAL3393 | Uncharacterized protein | \|B4EEY1_BURCJ | 1.57 | 187 | 1 |
| BCAM2399 hyi | Hydroxypyruvate isomerase | \|B4EI16_BURCJ | 1.57 | 258 | 3 |
| BCAS0738 | Putative short-chain dehydrogenase family protein | \|B4EPC6_BURCJ | 1.57 | 255 | 23 |
| BCAL0463 | Putative thioredoxin | \|B4E7V5_BURCJ | 1.56 | 126 | 28 |
| BCAL2428 | Putative cytochrome C-related protein | \|B4E6F4_BURCJ | 1.56 | 119 | 7 |
| BCAL1268 folP | Dihydropteroate synthase | \|B4E5F5_BURCJ | 1.56 | 292 | 5 |
| BCAL2941 | Putative exported transglycosylase | \|B4EB31_BURCJ | 1.56 | 405 | 4 |
| BCAL2666 | Uncharacterized protein | \|B4E8F9_BURCJ | 1.56 | 116 | 0 |
| BCAL2373 | Putative globin | \|B4E5Q1_BURCJ | 1.56 | 136 | 1 |
| BCAM2733 | Acylphosphatase | \|B4EL02_BURCJ | 1.56 | 98 | 6 |
| BCAM2618 argT | Putative periplasmic lysine-arginine-ornithine-binding protein | \|B4EKA4_BURCJ | 1.56 | 260 | 5 |
| BCAL2090 tsf | Elongation factor Ts | \|EFTS_BURCJ | 1.56 | 293 | 4 |
| BCAL2323 | Putative glutathione S-transferase | \|B4EER1_BURCJ | 1.56 | 230 | 3 |
| BCAL2172 | Putative phosphoesterase | \|B4ED94_BURCJ | 1.56 | 279 | 11 |
| BCAM2569 | IclR family regulatory protein | \|B4EJR8_BURCJ | 1.55 | 275 | 3 |
| BCAL3424 tpx | Probable thiol peroxidase | \|B4E5V4_BURCJ | 1.55 | 167 | 8 |
| BCAS0242 | Uncharacterized protein | \|B4EQ09_BURCJ | 1.55 | 244 | 5 |
| BCAM1492 | Putative exported protein | \|B4EJI3_BURCJ | 1.55 | 134 | 0 |
| BCAL3216 cysC | Adenylyl-sulfate kinase | \|B4EDD5_BURCJ | 1.55 | 187 | 6 |
| BCAL0368 cspD | Cold shock-like protein CspD | \|B4E6U0_BURCJ | 1.55 | 67 | 10 |
| BCAM0141 | Putative short-chain dehydrogenase | \|B4EH41_BURCJ | 1.55 | 252 | 2 |
| BCAL3031 | Uncharacterized protein | \|B4EBR5_BURCJ | 1.55 | 88 | 0 |
| BCAL3475 | Putative molybdopterin-containing oxidoreductase | \|B4E6K5_BURCJ | 1.55 | 691 | 4 |
| BCAL3328 | Putative hydrolase | \|B4EE49_BURCJ | 1.55 | 169 | 4 |
| BCAL2171 | Uncharacterized protein | \|B4ED93_BURCJ | 1.55 | 210 | 1 |

| | | | | | |
|---|---|---|---|---|---|
| BCAL1905 rpmE | 50S ribosomal protein L31 type B | \|RL31B_BURCJ | 1.55 | 86 | 4 |
| BCAL1263 greA | Transcription elongation factor GreA | \|B4E5F0_BURCJ | 1.55 | 158 | 6 |
| BCAM0916 | Uncharacterized protein | \|B4EN34_BURCJ | 1.54 | 148 | 3 |
| BCAL1035 otsB | Trehalose 6-phosphate phosphatase | \|B4ECJ7_BURCJ | 1.54 | 250 | 6 |
| BCAL0930 | Putative gamma-glutamyltransferase | \|B4EBW0_BURCJ | 1.54 | 546 | 8 |
| BCAL0431 | Uncharacterized protein | \|B4E7D9_BURCJ | 1.54 | 82 | 0 |
| BCAL3450 | NUDIX hydrolase | \|B4E5Y0_BURCJ | 1.54 | 147 | 5 |
| BCAL0194 | Putative oxidoreductase | \|B4EEC0_BURCJ | 1.54 | 289 | 7 |
| BCAL0575 ycgR | Flagellar brake protein YcgR | \|B4E8L7_BURCJ | 1.54 | 251 | 5 |
| BCAL0391 acpD | FMN-dependent NADH-azoreductase | \|B4E6W4_BURCJ | 1.54 | 198 | 4 |
| BCAM2713 | Putative exported protein | \|B4EKY2_BURCJ | 1.53 | 175 | 0 |
| BCAL0685 | IclR family regulatory protein | \|B4E9N5_BURCJ | 1.53 | 288 | 8 |
| BCAL0334 | Periplasmic solute-binding protein | \|B4E6Q6_BURCJ | 1.53 | 266 | 3 |
| BCAM0635 | AnsC family regulatory protein | \|B4EL56_BURCJ | 1.53 | 151 | 6 |
| BCAL0830 | Putative ParA family protein | \|B4EAU1_BURCJ | 1.53 | 271 | 1 |
| BCAM0280 | utative phospholipid-binding protein | \|B4EI91_BURCJ | 1.52 | 216 | 2 |
| BCAL0118 tag | DNA-3-methyladenine glycosylase I | \|B4EDJ2_BURCJ | 1.52 | 200 | 6 |
| BCAM0994 accD | Acetyl-coenzyme A carboxylase carboxyl transferase subunit beta | \|ACCD_BURCJ | 1.52 | 290 | 8 |
| BCAL2785 msrA | Peptide methionine sulfoxide reductase MsrA | \|B4E9I4_BURCJ | 1.52 | 186 | 2 |
| BCAL2159 | Uncharacterized protein | \|B4ED82_BURCJ | 1.52 | 108 | 0 |
| BCAL0002 | Carboxylate-amine ligase BceJ2315_00010 | \|CAAL_BURCJ | 1.52 | 371 | 6 |
| BCAL0211 | Uncharacterized protein | \|B4E597_BURCJ | 1.52 | 273 | 0 |
| BCAL2776 | Putative hydrolase | \|B4E9H5_BURCJ | 1.52 | 184 | 4 |
| BCAL2112 thiD | Phosphomethylpyrimidine kinase | \|B4ECQ2_BURCJ | 1.52 | 269 | 5 |
| BCAM2473 | DeoR family regulatory protein | \|B4EIN9_BURCJ | 1.52 | 252 | 3 |
| BCAL0985 | Rieske iron-sulphur protein | \|B4EAJ3_BURCJ | 1.52 | 133 | 3 |
| BCAM1800 | Uncharacterized protein | \|B4ELW0_BURCJ | 1.51 | 91 | 0 |
| BCAL3015 | Non-canonical purine NTP pyrophosphatase | \|B4EBP9_BURCJ | 1.51 | 208 | 5 |
| BCAL0875 | Uncharacterized protein | \|B4EBC6_BURCJ | 1.51 | 208 | 0 |
| BCAL1967 | Uncharacterized protein | \|B4EBJ5_BURCJ | 1.50 | 281 | 3 |
| BCAL2761 kdsB | 3-deoxy-manno-octulosonate cytidylyltransferase | \|KDSB_BURCJ | 1.50 | 263 | 6 |
| BCAL2187 | Putative exported protein | \|B4EDA9_BURCJ | 1.50 | 358 | 0 |
| BCAL3142 | UPF0301 protein BceJ2315_30870 | \|Y3087_BURCJ | 1.50 | 192 | 0 |

| | | | | | |
|---|---|---|---|---|---|
| BCAL2983A | Putative lipoprotein | \|B4EB83_BURCJ | 1.50 | 232 | 0 |

| | Protein names | Seq. Description | Actual fold d | Seq. Len | #GOs |
|---|---|---|---|---|---|
| **Increased abundance in the mutant** | | | | | |
| BCAL0438 | Putative DNA-3-methyladenine glycosylase II | \|B4E7E6_BURCJ | 9.82 | 319 | 3 |
| BCAM1704 | 2,3-butanediol dehydrogenase | \|B4EL82_BURCJ | 8.82 | 365 | 5 |
| BCAL1265 | Uncharacterized protein | \|B4E5F2_BURCJ | 6.97 | 175 | 2 |
| BCAL0863 | Cysteine peptidase, family C56 | \|B4EBB4_BURCJ | 6.73 | 193 | 5 |
| BCAL0327 | Putative GTP cyclohydrolase 1 type 2 | \|B4E650_BURCJ | 5.22 | 248 | 4 |
| uvrA | UvrABC system protein A | \|B4EAT6_BURCJ | 4.87 | 961 | 16 |
| rpsU | 30S ribosomal protein S21 | \|B4EN33_BURCJ | 4.77 | 70 | 4 |
| BCAL3429 | Ribonucleoside-diphosphate reductase | \|B4E5V9_BURCJ | 4.43 | 1003 | 20 |
| pyrE | Orotate phosphoribosyltransferase | \|B4E589_BURCJ | 4.33 | 228 | 4 |
| BCAM1012 | Putative histone-like protein | \|B4EFM0_BURCJ | 4.20 | 149 | 0 |
| otsA | Alpha,alpha-trehalose-phosphate synthase | \|B4EK36_BURCJ | 4.13 | 468 | 1 |
| BCAL0884 | Putative acyl-CoA dehydrogenase oxidoreductase protein | \|B4EBD5_BURCJ | 4.11 | 595 | 3 |
| ileS | Isoleucine--tRNA ligase | \|SYI_BURCJ | 4.02 | 945 | 9 |
| BCAL3277 | DNA repair protein RecN | \|B4EDZ9_BURCJ | 3.92 | 549 | 6 |
| rpoD | RNA polymerase sigma factor RpoD | \|B4EN36_BURCJ | 3.84 | 621 | 7 |
| arcB | Ornithine carbamoyltransferase | \|OTC_BURCJ | 3.83 | 309 | 7 |
| hutH | Histidine ammonia-lyase | \|HUTH_BURCJ | 3.83 | 507 | 5 |
| BCAL2721 | NOL1/NOP2/Sun family protein | \|B4E8Z4_BURCJ | 3.81 | 421 | 11 |
| BCAM2739 | MoaA/NifB/PqqE family protein | \|B4ELD7_BURCJ | 3.79 | 386 | 1 |
| codA | Cytosine deaminase | \|B4EJ11_BURCJ | 3.76 | 413 | 5 |
| BCAL2327 | Putative acyl-CoA dehydrogenase family protein | \|B4EER5_BURCJ | 3.59 | 398 | 6 |
| parE | DNA topoisomerase 4 subunit B | \|B4E6I1_BURCJ | 3.58 | 660 | 8 |
| BCAL0047 | Putative acyl-CoA dehydrogenase | \|B4EEZ6_BURCJ | 3.55 | 550 | 13 |
| BCAL086 | YjgF family protein | \|B4EBB8_BURCJ | 3.52 | 128 | 9 |
| ilvG | Acetolactate synthase, large subunit | \|B4EF21_BURCJ | 3.50 | 567 | 3 |
| BCAL1722 | Putative exported chitinase | \|B4E9C1_BURCJ | 3.50 | 451 | 3 |
| cafA | Ribonuclease G | \|B4E5S3_BURCJ | 3.40 | 489 | 8 |
| thiC | Phosphomethylpyrimidine synthase | \|THIC_BURCJ | 3.40 | 643 | 3 |
| hisB | Imidazoleglycerol-phosphate dehydratase | \|HIS7_BURCJ | 3.39 | 195 | 4 |

| | | | | | |
|---|---|---|---|---|---|
| adiA | Biodegradative arginine decarboxylase | \|B4EG92_BURCJ | 3.39 | 754 | 9 |
| rhlE1 | ATP-dependent RNA helicase RhlE | \|B4EBW3_BURCJ | 3.38 | 477 | 11 |
| rne1 | Ribonuclease E | \|B4EAJ6_BURCJ | 3.37 | 1054 | 7 |
| gcvP | Glycine dehydrogenase (decarboxylating) | \|GCSP_BURCJ | 3.37 | 975 | 5 |
| glt1 | Glutamate synthase large subunit | \|B4E612_BURCJ | 3.34 | 1607 | 15 |
| rpml | 50S ribosomal protein L35 | \|RL35_BURCJ | 3.34 | 65 | 4 |
| BCAM0810 | Putative aromatic oxygenase | \|B4EML2_BURCJ | 3.29 | 423 | 4 |
| sucA | 2-oxoglutarate dehydrogenase E1 component | \|B4E7L9_BURCJ | 3.28 | 954 | 9 |
| BCAL0671 | Carbonic anhydrase | \|B4E9M1_BURCJ | 3.27 | 255 | 8 |
| BCAL1240 | Putative malonate decarboxylase alpha-subunit | \|B4EEH7_BURCJ | 3.26 | 548 | 1 |
| BCAL2207 | Putative dihydrolipoamide dehydrogenase | \|B4EDT6_BURCJ | 3.25 | 589 | 15 |
| BCAL0885 | Putative 3-hydroxyacyl-CoA dehydrogenase oxidoreductase | \|B4EBD6_BURCJ | 3.23 | 811 | 11 |
| rpsG | 30S ribosomal protein S7 | \|RS7_BURCJ | 3.20 | 156 | 5 |
| BCAL2107 | ABC transporter ATP-binding protein | \|B4ECP7_BURCJ | 3.19 | 643 | 8 |
| rpoB | DNA-directed RNA polymerase subunit beta | \|RPOB_BURCJ | 3.17 | 1368 | 6 |
| trpS | Tryptophan--tRNA ligase | \|B4ED91_BURCJ | 3.17 | 400 | 6 |
| pnp | Polyribonucleotide nucleotidyltransferase | \|PNP_BURCJ | 3.15 | 713 | 7 |
| mutS | DNA mismatch repair protein MutS | \|MUTS_BURCJ | 3.14 | 885 | 4 |
| rpsL | 30S ribosomal protein S12 | \|RS12_BURCJ | 3.13 | 126 | 5 |
| rplP | 50S ribosomal protein L16 | \|RL16_BURCJ | 3.07 | 138 | 5 |
| uvrB | UvrABC system protein B | \|B4EEP1_BURCJ | 3.06 | 696 | 12 |
| BCAM0901 | Putative AMP nucleosidase | \|B4EN19_BURCJ | 3.05 | 508 | 3 |
| rpsK | 30S ribosomal protein S11 | \|RS11_BURCJ | 3.04 | 133 | 5 |
| mnmG | tRNA uridine 5-carboxymethylaminomethyl modification enzyme MnmG | \|MNMG_BURCJ | 3.04 | 656 | 3 |
| BCAM2619 | Succinylglutamate desuccinylase/aspartoacylase family protein | \|B4EKA5_BURCJ | 3.01 | 371 | 0 |
| BCAM0359 | Putative pyridoxal-dependent decarboxylase | \|B4EIU7_BURCJ | 3.00 | 450 | 1 |
| BCAL0080 | Putative cytochrome | \|B4EF33_BURCJ | 2.98 | 295 | 9 |
| leuS | Leucine--tRNA ligase | \|SYL_BURCJ | 2.98 | 864 | 8 |
| ilvE | Putative branched-chain amino acid aminotransferase IlvE | \|B4EA28_BURCJ | 2.96 | 307 | 3 |
| secB | Protein-export protein SecB | \|SECB_BURCJ | 2.96 | 163 | 5 |
| ilvD | Dihydroxy-acid dehydratase | \|B4EHE1_BURCJ | 2.94 | 619 | 4 |
| secA | Protein translocase subunit SecA | \|SECA_BURCJ | 2.94 | 933 | 5 |
| rpoC | DNA-directed RNA polymerase subunit beta | \|RPOC_BURCJ | 2.93 | 1413 | 5 |

| | | | | | |
|---|---|---|---|---|---|
| BCAS0727 | Putative oxidoredutase | \|B4EPB5_BURCJ | 2.92 | 371 | 8 |
| ffh | Signal recognition particle protein | \|B4E5W3_BURCJ | 2.91 | 455 | 7 |
| proS | Proline--tRNA ligase | \|SYP_BURCJ | 2.90 | 578 | 7 |
| BCAL0437 | O6-methylguanine-DNA methyltransferase | \|B4E7E5_BURCJ | 2.90 | 363 | 5 |
| paaH | 3-hydroxybutyryl-CoA dehydrogenase | \|B4EL90_BURCJ | 2.89 | 518 | 10 |
| cheB1 | Chemotaxis response regulator protein-glutamate methylesterase | \|B4EE59_BURCJ | 2.88 | 363 | 10 |
| clpA | Putative ATP-dependent Clp protease ATP-binding subunit | \|B4E903_BURCJ | 2.86 | 766 | 6 |
| maeB | NADP-dependent malic enzyme | \|B4EEY3_BURCJ | 2.86 | 785 | 8 |
| hutU | Urocanate hydratase | \|B4EDX3_BURCJ | 2.85 | 562 | 4 |
| BCAL0390 | Metallo peptidase, family M61 | \|B4E6W3_BURCJ | 2.82 | 599 | 3 |
| BCAM0964 | Putative lyase | \|B4EFA9_BURCJ | 2.80 | 333 | 1 |
| odhL | Dihydrolipoyl dehydrogenase | \|B4E7M1_BURCJ | 2.79 | 476 | 18 |
| BCAL2938 | ABC transporter ATP-binding protein | \|B4EB28_BURCJ | 2.79 | 344 | 7 |
| moeA3 | Molybdopterin biosynthesis protein MoeA 3 | \|B4EB10_BURCJ | 2.79 | 415 | 12 |
| aspS | Aspartate--tRNA ligase | \|SYD_BURCJ | 2.78 | 600 | 7 |
| pheT | Phenylalanine--tRNA ligase beta subunit | \|B4E7J0_BURCJ | 2.77 | 809 | 10 |
| acoE | Acetyl-coenzyme A synthetase | \|B4EEM3_BURCJ | 2.76 | 660 | 7 |
| ribH | 6,7-dimethyl-8-ribityllumazine synthase | \|RISB_BURCJ | 2.76 | 171 | 5 |
| bioC | Malonyl-[acyl-carrier protein] O-methyltransferase | \|B4EA72_BURCJ | 2.75 | 321 | 2 |
| BCAL1098 | Putative exodeoxyribonuclease V alpha chain | \|B4ED34_BURCJ | 2.74 | 769 | 8 |
| dadA | D-amino acid dehydrogenase small subunit | \|DADA_BURCJ | 2.73 | 428 | 6 |
| hemN | Coproporphyrinogen-III oxidase | \|B4ECB5_BURCJ | 2.71 | 463 | 4 |
| speF | Ornithine decarboxylase | \|B4EG91_BURCJ | 2.70 | 726 | 6 |
| BCAM2561 | Putative 4-aminobutyrate aminotransferase | \|B4EJR0_BURCJ | 2.70 | 427 | 4 |
| BCAM0917 | DNA primase | \|B4EN35_BURCJ | 2.69 | 625 | 8 |
| leuA | 2-isopropylmalate synthase | \|B4EHL9_BURCJ | 2.67 | 570 | 4 |
| BCAL2456 | ABC transporter ATP-binding protein | \|B4E6I2_BURCJ | 2.67 | 649 | 7 |
| paaK | Phenylacetate-coenzyme A ligase | \|B4EL89_BURCJ | 2.67 | 440 | 4 |
| alaS | Alanine--tRNA ligase | \|B4E6Y2_BURCJ | 2.67 | 874 | 8 |
| BCAL0513 | Aldo/keto reductase family protein | \|B4E804_BURCJ | 2.67 | 347 | 9 |
| glnS | Glutamine--tRNA ligase | \|B4E6X8_BURCJ | 2.66 | 569 | 6 |
| parC | DNA topoisomerase 4 subunit A | \|B4E6I0_BURCJ | 2.66 | 773 | 11 |
| kynA | Tryptophan 2,3-dioxygenase | \|T23O_BURCJ | 2.66 | 311 | 6 |

<table>
<tr><td>BCAL0380</td><td>ABC transporter ATP-binding subunit</td><td>|B4E6V3_BURCJ</td><td>2.63</td><td>555</td><td>15</td></tr>
<tr><td>BCAL2641</td><td>Putative ornithine decarboxylase</td><td>|B4E8D5_BURCJ</td><td>2.62</td><td>759</td><td>9</td></tr>
<tr><td>iscS</td><td>Cysteine desulfurase</td><td>|B4EDS7_BURCJ</td><td>2.62</td><td>407</td><td>3</td></tr>
<tr><td>prfA</td><td>Peptide chain release factor 1</td><td>|RF1_BURCJ</td><td>2.61</td><td>360</td><td>7</td></tr>
<tr><td>rpsM</td><td>30S ribosomal protein S13</td><td>|RS13_BURCJ</td><td>2.58</td><td>121</td><td>5</td></tr>
<tr><td>lepA1</td><td>Elongation factor 4</td><td>|B4EAH5_BURCJ</td><td>2.55</td><td>597</td><td>10</td></tr>
<tr><td>BCAL0462</td><td>DNA topoisomerase</td><td>|B4E7V4_BURCJ</td><td>2.55</td><td>865</td><td>9</td></tr>
<tr><td>nuoD</td><td>NADH-quinone oxidoreductase subunit D</td><td>|NUOD_BURCJ</td><td>2.53</td><td>417</td><td>7</td></tr>
<tr><td>BCAL1937</td><td>Putative phosphorous metabolism-related protein</td><td>|B4EB23_BURCJ</td><td>2.52</td><td>549</td><td>1</td></tr>
<tr><td>pdhB</td><td>Dihydrolipoamide acetyltransferase component of pyruvate dehydrogenase co</td><td>|B4EDT7_BURCJ</td><td>2.50</td><td>547</td><td>10</td></tr>
<tr><td>mnmA</td><td>tRNA-specific 2-thiouridylase MnmA</td><td>|MNMA_BURCJ</td><td>2.50</td><td>391</td><td>5</td></tr>
<tr><td>glpD</td><td>Putative glycerol-3-phosphate dehydrogenase</td><td>|B4EBV6_BURCJ</td><td>2.50</td><td>507</td><td>10</td></tr>
<tr><td>guaB</td><td>Inosine-5-monophosphate dehydrogenase</td><td>|B4EC70_BURCJ</td><td>2.50</td><td>486</td><td>10</td></tr>
<tr><td>BCAL2338</td><td>Putative NADH dehydrogenase I chain G</td><td>|B4E5L6_BURCJ</td><td>2.49</td><td>776</td><td>6</td></tr>
<tr><td>prpC</td><td>Citrate synthase</td><td>|B4EKX1_BURCJ</td><td>2.47</td><td>390</td><td>6</td></tr>
<tr><td>mpl</td><td>UDP-N-acetylmuramate:L-alanyl-gamma-D-glutamyl-meso-diaminopimelate lig</td><td>|B4E5U6_BURCJ</td><td>2.47</td><td>461</td><td>3</td></tr>
<tr><td>argA</td><td>Amino-acid acetyltransferase</td><td>|B4EDW0_BURCJ</td><td>2.47</td><td>459</td><td>5</td></tr>
<tr><td>nrdB</td><td>Ribonucleoside-diphosphate reductase subunit beta</td><td>|B4E5V8_BURCJ</td><td>2.46</td><td>403</td><td>6</td></tr>
<tr><td>BCAL0808</td><td>UPF0042 nucleotide-binding protein BceJ2315_08000</td><td>|Y800_BURCJ</td><td>2.46</td><td>302</td><td>1</td></tr>
<tr><td>BCAL0205</td><td>NADP-dependent malic enzyme</td><td>|B4E590_BURCJ</td><td>2.45</td><td>756</td><td>8</td></tr>
<tr><td>BCAL0886</td><td>Putative 3-ketoacyl-CoA thiolase</td><td>|B4EBD7_BURCJ</td><td>2.45</td><td>399</td><td>19</td></tr>
<tr><td>prpD</td><td>2-methylcitrate dehydratase</td><td>|B4EFA7_BURCJ</td><td>2.44</td><td>483</td><td>4</td></tr>
<tr><td>trpB</td><td>Tryptophan synthase beta chain</td><td>|B4EFK0_BURCJ</td><td>2.44</td><td>397</td><td>3</td></tr>
<tr><td>clpX</td><td>ATP-dependent Clp protease ATP-binding subunit ClpX</td><td>|CLPX_BURCJ</td><td>2.44</td><td>423</td><td>5</td></tr>
<tr><td>ansB</td><td>L-asparaginase II</td><td>|B4EBI1_BURCJ</td><td>2.43</td><td>340</td><td>5</td></tr>
<tr><td>BCAL0650</td><td>Putative pyruvate-flavodoxin oxidoreductase</td><td>|B4E972_BURCJ</td><td>2.43</td><td>1190</td><td>11</td></tr>
<tr><td>murG</td><td>UDP-N-acetylglucosamine--N-acetylmuramyl-(pentapeptide) pyrophosphoryl-u</td><td>|MURG_BURCJ</td><td>2.42</td><td>367</td><td>12</td></tr>
<tr><td>BCAM0908</td><td>Putative iron-sulfur protein</td><td>|B4EN26_BURCJ</td><td>2.41</td><td>368</td><td>2</td></tr>
<tr><td>typA</td><td>GTP-binding protein</td><td>|B4E7L8_BURCJ</td><td>2.41</td><td>608</td><td>8</td></tr>
<tr><td>glmS1</td><td>Glutamine--fructose-6-phosphate aminotransferase [isomerizing]</td><td>|B4E934_BURCJ</td><td>2.41</td><td>605</td><td>6</td></tr>
<tr><td>BCAL1449</td><td>Putative helicase</td><td>|B4E715_BURCJ</td><td>2.37</td><td>818</td><td>2</td></tr>
<tr><td>paaK</td><td>Phenylacetate-coenzyme A ligase</td><td>|B4E7B5_BURCJ</td><td>2.37</td><td>432</td><td>4</td></tr>
<tr><td>carB</td><td>Carbamoyl-phosphate synthase large chain</td><td>|B4EEJ9_BURCJ</td><td>2.37</td><td>1084</td><td>9</td></tr>
</table>

| | | | | | |
|---|---|---|---|---|---|
| BCAL1504 | Pseudouridine synthase | \|B4E7K8_BURCJ | 2.36 | 583 | 2 |
| pyrG | CTP synthase | \|PYRG_BURCJ | 2.36 | 552 | 5 |
| purB | Adenylosuccinate lyase | \|B4EEU9_BURCJ | 2.36 | 462 | 6 |
| katG1 | Catalase-peroxidase 1 | \|KATG1_BURCJ | 2.35 | 728 | 7 |
| bceC | UDP-glucose dehydrogenase | \|B4EMQ8_BURCJ | 2.35 | 470 | 3 |
| ppc | Phosphoenolpyruvate carboxylase | \|B4E8C5_BURCJ | 2.33 | 989 | 7 |
| rplB | 50S ribosomal protein L2 | \|RL2_BURCJ | 2.32 | 275 | 6 |
| glyQ | Putative glycyl-tRNA synthetase alpha chain | \|B4EBF5_BURCJ | 2.32 | 334 | 7 |
| paaZ | Putative phenylacetic acid degradation oxidoreductase | \|B4E7B9_BURCJ | 2.32 | 568 | 1 |
| BCAL0281 | Deoxyguanosinetriphosphate triphosphohydrolase-like protein | \|B4E604_BURCJ | 2.31 | 383 | 3 |
| BCAL1249 | Putative PHB depolymerase | \|B4EEI6_BURCJ | 2.31 | 405 | 0 |
| BCAL2437 | Uncharacterized protein | \|B4E6G3_BURCJ | 2.30 | 305 | 11 |
| pncB | Nicotinate phosphoribosyltransferase | \|B4EAK6_BURCJ | 2.29 | 399 | 6 |
| rpsI | 30S ribosomal protein S9 | \|RS9_BURCJ | 2.28 | 130 | 4 |
| rplM | 50S ribosomal protein L13 | \|RL13_BURCJ | 2.27 | 142 | 4 |
| glnA | Glutamine synthetase | \|B4EDV3_BURCJ | 2.26 | 471 | 9 |
| citB | Aconitate hydratase | \|B4EFA6_BURCJ | 2.26 | 905 | 32 |
| purF | Amidophosphoribosyltransferase | \|B4EFK7_BURCJ | 2.26 | 510 | 7 |
| glyS | Glycine--tRNA ligase beta subunit | \|B4EBF4_BURCJ | 2.25 | 699 | 6 |
| fabZ | 3-hydroxyacyl-[acyl-carrier-protein] dehydratase FabZ | \|B4ECM0_BURCJ | 2.24 | 160 | 5 |
| rplT | 50S ribosomal protein L20 | \|RL20_BURCJ | 2.24 | 119 | 5 |
| BCAL2209 | Pyruvate dehydrogenase E1 component | \|B4EDT8_BURCJ | 2.24 | 898 | 12 |
| BCAL2420 | Putative depolymerase/histone-like protein | \|B4E6E6_BURCJ | 2.24 | 496 | 0 |
| ppsA | Phosphoenolpyruvate synthase | \|B4EC81_BURCJ | 2.24 | 800 | 8 |
| thrS | Threonine--tRNA ligase | \|SYT_BURCJ | 2.23 | 635 | 7 |
| smc | Chromosome partition protein Smc | \|B4ECN8_BURCJ | 2.23 | 1170 | 10 |
| dapB | 4-hydroxy-tetrahydrodipicolinate reductase | \|DAPB_BURCJ | 2.22 | 265 | 6 |
| BCAL2783 | Putative cyclopropane-fatty-acyl-phospholipid synthase | \|B4E9I2_BURCJ | 2.22 | 406 | 3 |
| rnr | Ribonuclease R | \|B4E849_BURCJ | 2.21 | 824 | 9 |
| BCAL3359 | Glutamate dehydrogenase | \|B4EEU7_BURCJ | 2.21 | 428 | 4 |
| BCAL2735 | Isocitrate dehydrogenase [NADP] | \|B4E908_BURCJ | 2.21 | 745 | 7 |
| gapA | Glyceraldehyde 3-phosphate dehydrogenase 1 | \|B4EEX6_BURCJ | 2.20 | 336 | 9 |
| BCAM0601 | Putative HSP70 protein | \|B4EL22_BURCJ | 2.20 | 613 | 8 |

| | | | | | |
|---|---|---|---|---|---|
| pckG | Phosphoenolpyruvate carboxykinase [GTP] | \|B4EK44_BURCJ | 2.19 | 616 | 8 |
| rplO | 50S ribosomal protein L15 | \|RL15_BURCJ | 2.19 | 144 | 5 |
| BCAL2772 | Putative AMP-binding enzyme | \|B4E9H1_BURCJ | 2.19 | 494 | 7 |
| fdsB | NAD-dependent formate dehydrogenase beta subunit | \|B4EB76_BURCJ | 2.19 | 525 | 8 |
| rpmC | 50S ribosomal protein L29 | \|RL29_BURCJ | 2.19 | 64 | 4 |
| metG | Methionine--tRNA ligase | \|SYM_BURCJ | 2.18 | 718 | 8 |
| BCAL1013 | Uncharacterized protein | \|B4EA40_BURCJ | 2.17 | 397 | 0 |
| accC | Biotin carboxylase | \|B4E5V1_BURCJ | 2.17 | 455 | 22 |
| dfp | Putative DNA/pantothenate metabolism flavoprotein | \|B4E8Z9_BURCJ | 2.17 | 403 | 5 |
| BCAL3287 | Putative FAD-binding oxidase | \|B4EE09_BURCJ | 2.17 | 469 | 11 |
| rplV | 50S ribosomal protein L22 | \|RL22_BURCJ | 2.17 | 109 | 5 |
| ispG | 4-hydroxy-3-methylbut-2-en-1-yl diphosphate synthase | \|ISPG_BURCJ | 2.16 | 434 | 7 |
| nadB | utative L-aspartate oxidase | \|B4E8Y9_BURCJ | 2.16 | 528 | 9 |
| rpsC | 30S ribosomal protein S3 | \|RS3_BURCJ | 2.16 | 266 | 5 |
| BCAL0348 | Putative type VI secretion system protein TssA | \|B4E6S0_BURCJ | 2.16 | 373 | 0 |
| rpsN | 30S ribosomal protein S14 | \|RS14_BURCJ | 2.16 | 101 | 5 |
| pgi | Glucose-6-phosphate isomerase | \|G6PI_BURCJ | 2.15 | 540 | 8 |
| BCAL1011 | Sigma-54 interacting response regulator protein | \|B4EA42_BURCJ | 2.15 | 461 | 5 |
| BCAM2432 | Putative biotin-dependent carboxyl transferase | \|B4EIJ8_BURCJ | 2.14 | 535 | 19 |
| BCAL2040 | Polysaccharide deacetylase | \|B4EC47_BURCJ | 2.14 | 317 | 4 |
| acnB | Aconitate hydratase 2 | \|B4EM96_BURCJ | 2.14 | 861 | 10 |
| BCAM1442 | Putative methylmalonate-semialdehyde dehydrogenase | \|B4EIZ7_BURCJ | 2.14 | 505 | 6 |
| BCAL3013 | Uncharacterized protein | \|B4EBP7_BURCJ | 2.13 | 307 | 0 |
| cysI | Putative sulfite reductase | \|B4E8V8_BURCJ | 2.12 | 559 | 2 |
| srpH | Serine acetyltransferase | \|B4EG39_BURCJ | 2.12 | 306 | 5 |
| BCAL1994 | Lon protease | \|B4EBM1_BURCJ | 2.10 | 807 | 11 |
| nuoL | NADH-ubiquinone oxidoreductase I chain L | \|B4E5L1_BURCJ | 2.10 | 684 | 4 |
| gyrB | DNA gyrase subunit B | \|B4E7C9_BURCJ | 2.10 | 824 | 12 |
| nadA | Quinolinate synthase A | \|B4E8Z1_BURCJ | 2.09 | 380 | 7 |
| dctD | C4-dicarboxylate transport transcriptional regulatory protein | \|B4EA50_BURCJ | 2.09 | 450 | 9 |
| BCAL1072 | Uncharacterized protein | \|B4ED08_BURCJ | 2.08 | 491 | 0 |
| rho | Transcription termination factor Rho | \|B4EAY7_BURCJ | 2.08 | 420 | 7 |
| aceA | Isocitrate lyase | \|B4ECQ9_BURCJ | 2.08 | 435 | 10 |

| | | | | | |
|---|---|---|---|---|---|
| BCAL1830 | Putative 2-nitropropane dioxygenase | \|B4EAC9_BURCJ | 2.07 | 396 | 5 |
| recA | Protein RecA | \|RECA_BURCJ | 2.07 | 347 | 8 |
| BCAL3451 | Uncharacterized protein | \|B4E5Y1_BURCJ | 2.07 | 289 | 2 |
| guaA | GMP synthase [glutamine-hydrolyzing] | \|GUAA_BURCJ | 2.07 | 539 | 6 |
| BCAL1278 | Putative exopolyphosphatase | \|B4E5G5_BURCJ | 2.06 | 504 | 4 |
| argD | Acetylornithine aminotransferase | \|B4E8X4_BURCJ | 2.05 | 394 | 5 |
| gca | GDP-mannose 4,6-dehydratase | \|B4EFL2_BURCJ | 2.05 | 348 | 19 |
| nagZ1 | Beta-hexosaminidase 1 | \|B4EA43_BURCJ | 2.05 | 342 | 2 |
| acoD | Acetaldehyde dehydrogenase | \|B4EF17_BURCJ | 2.05 | 508 | 10 |
| BCAL1502 | Putative aminotransferase | \|B4E7K6_BURCJ | 2.05 | 400 | 4 |
| cydA | Cytochrome d ubiquinol oxidase subunit I | \|B4EAP8_BURCJ | 2.05 | 526 | 8 |
| BCAL0154 | Histone-like nucleoid-structuring (H-NS) protein | \|B4EE79_BURCJ | 2.04 | 97 | 4 |
| der | GTPase Der | \|DER_BURCJ | 2.04 | 445 | 3 |
| lpxC | UDP-3-O-[3-hydroxymyristoyl] N-acetylglucosamine deacetylase | \|LPXC_BURCJ | 2.04 | 305 | 5 |
| BCAM2168 | Putative amylo-alpha-1,6-glucosidase | \|B4EFW5_BURCJ | 2.03 | 740 | 0 |
| kynU | Kynureninase | \|B4E9J0_BURCJ | 2.02 | 417 | 9 |
| ilvI | Acetolactate synthase isozyme III large subunit | \|B4E5N7_BURCJ | 2.02 | 587 | 6 |
| BCAS0739 | Putative acetyl-CoA synthetase | \|B4EPC7_BURCJ | 2.02 | 555 | 4 |
| rhlE2 | Putative ATP-dependent RNA helicase 2 | \|B4E6D8_BURCJ | 2.02 | 494 | 11 |
| BCAL0301 | ABC transporter ATP-binding protein | \|B4E624_BURCJ | 2.01 | 273 | 10 |
| wbxD | Glycosyltransferase | \|B4ECE5_BURCJ | 2.01 | 655 | 1 |
| valS | Valine--tRNA ligase | \|B4E714_BURCJ | 2.00 | 955 | 8 |
| BCAL0870 | Putative oxidoreductase | \|B4EBC1_BURCJ | 2.00 | 1341 | 4 |
| gatA | Glutamyl-tRNA(Gln) amidotransferase subunit A | \|GATA_BURCJ | 2.00 | 496 | 5 |
| putA | Bifunctional PutA protein [includes: proline dehydrogenase delta-1-pyrroline-5 | \|B4EF02_BURCJ | 2.00 | 1310 | 7 |
| BCAM0684 | Putative oxidoreductase | \|B4ELN5_BURCJ | 2.00 | 459 | 20 |
| BCAM1556 | Putative succinylglutamate desuccinylase/ aspartoacylase | \|B4EJP8_BURCJ | 2.00 | 343 | 12 |
| gmhA | Phosphoheptose isomerase | \|B4EE87_BURCJ | 1.99 | 194 | 1 |
| BCAM1978 | Aminotransferase class-III | \|B4EMW0_BURCJ | 1.99 | 446 | 20 |
| BCAL2320 | Putative aminotransferase | \|B4EEQ8_BURCJ | 1.99 | 384 | 5 |
| BCAL0377 | Metallo peptidase, subfamily M24B 1 | \|B4E6V0_BURCJ | 1.99 | 604 | 4 |
| BCAM0004 | Putative partitioning protein ParB | \|B4EFY9_BURCJ | 1.99 | 353 | 6 |
| pepA | Probable cytosol aminopeptidase | \|AMPA_BURCJ | 1.98 | 503 | 5 |

| | | | | | |
|---|---|---|---|---|---|
| BCAM1423 | Putative AMP-binding enzyme | \|B4EIX8_BURCJ | 1.96 | 550 | 8 |
| tyrB | Aromatic amino acid aminotransferase | \|B4EEP2_BURCJ | 1.96 | 399 | 14 |
| lpdV | Dihydrolipoyl dehydrogenase | \|B4EEF2_BURCJ | 1.96 | 463 | 10 |
| dalK | Putative carbohydrate kinase | \|B4E9J8_BURCJ | 1.96 | 493 | 3 |
| zwf | Glucose-6-phosphate 1-dehydrogenase | \|B4EJU9_BURCJ | 1.95 | 491 | 7 |
| hflK | Protein HflK | \|B4EAW2_BURCJ | 1.95 | 448 | 4 |
| BCAL1658 | Putative ribose ABC transporter ATP-binding protein | \|B4E8T1_BURCJ | 1.93 | 537 | 1 |
| BCAM0299 | Putative zinc-binding alcoholdehydrogenase | \|B4EIB1_BURCJ | 1.93 | 346 | 1 |
| acnA | Aconitate hydratase 1 | \|B4EKX0_BURCJ | 1.93 | 864 | 20 |
| rpoN | RNA polymerase sigma-54 factor | \|B4EAS4_BURCJ | 1.93 | 501 | 11 |
| rpsP | 30S ribosomal protein S16 | \|RS16_BURCJ | 1.93 | 84 | 4 |
| rpsR | 30S ribosomal protein S18 | \|RS18_BURCJ | 1.92 | 91 | 5 |
| BCAL3143 | Uncharacterized protein | \|B4ECT4_BURCJ | 1.92 | 435 | 0 |
| tkrA | 2-ketogluconate reductase | \|B4E9Y4_BURCJ | 1.92 | 321 | 5 |
| ispB | Octaprenyl-diphosphate synthase | \|B4E5X3_BURCJ | 1.91 | 331 | 13 |
| BCAL3333 | Putative ubiquinone biosynthesis-related protein | \|B4EES1_BURCJ | 1.91 | 395 | 2 |
| potG | Putrescine ABC transporter ATP-binding protein | \|B4EAC2_BURCJ | 1.91 | 386 | 7 |
| metH1 | Putative 5-methyltetrahydrofolate--homocysteine methyltransferase | \|B4E9N1_BURCJ | 1.90 | 905 | 5 |
| BCAL2326 | Putative acyl-CoA dehydrogenase family protein | \|B4EER4_BURCJ | 1.90 | 377 | 4 |
| argH | Argininosuccinate lyase | \|ARLY_BURCJ | 1.90 | 469 | 4 |
| BCAL0362 | Uncharacterized protein | \|B4E6T4_BURCJ | 1.90 | 296 | 4 |
| purH | Bifunctional purine biosynthesis protein PurH | \|PUR9_BURCJ | 1.89 | 521 | 6 |
| ligA | DNA ligase | \|DNLJ_BURCJ | 1.89 | 691 | 6 |
| BCAM0817 | Putative acetolactate synthase | \|B4EML9_BURCJ | 1.88 | 546 | 1 |
| lysS | Lysine--tRNA ligase | \|SYK_BURCJ | 1.88 | 508 | 9 |
| gtaB | Putative UTP-glucose-1-phosphate uridylyltransferase | \|B4EFL8_BURCJ | 1.88 | 295 | 3 |
| rpsD | 30S ribosomal protein S4 | \|RS4_BURCJ | 1.88 | 207 | 5 |
| BCAS0610 | Putative acyl-CoA dehydrogenase | \|B4EP04_BURCJ | 1.88 | 596 | 3 |
| BCAL1831 | Putative betaine aldehyde dehydrogenase | \|B4EAD0_BURCJ | 1.87 | 478 | 6 |
| phbA | Acetyl-CoA acetyltransferase | \|B4EAG1_BURCJ | 1.86 | 393 | 10 |
| BCAS0667 | Uncharacterized protein | \|B4EP61_BURCJ | 1.86 | 999 | 0 |
| BCAL1663 | PrkA family serine protein kinase | \|B4E8T6_BURCJ | 1.86 | 640 | 2 |
| hslU | ATP-dependent protease ATPase subunit HslU | \|HSLU_BURCJ | 1.85 | 447 | 9 |

| | | | | | |
|---|---|---|---|---|---|
| BCAL1796 | Putative saccharopine dehydrogenase | \|B4E9X7_BURCJ | 1.84 | 366 | 4 |
| rplQ | 50S ribosomal protein L17 | \|RL17_BURCJ | 1.84 | 131 | 4 |
| rpoS | RNA polymerase sigma factor RpoS | \|B4EAX7_BURCJ | 1.84 | 362 | 7 |
| BCAL2294 | LysR family regulatory protein | \|B4EEN3_BURCJ | 1.84 | 324 | 2 |
| purL | Phosphoribosylformylglycinamidine synthase | \|B4EBL4_BURCJ | 1.84 | 1354 | 7 |
| BCAS0010 | Putative activator of osmoprotectant transporter | \|B4EQD8_BURCJ | 1.83 | 223 | 0 |
| BCAL3056 | Putative aminotransferase | \|B4EBU0_BURCJ | 1.83 | 398 | 1 |
| infB | Translation initiation factor IF-2 | \|IF2_BURCJ | 1.83 | 971 | 8 |
| dxs | \|DXS_BURCJ 1-deoxy-D-xylulose-5-phosphate synthase | \|DXS_BURCJ | 1.83 | 634 | 5 |
| cca | Multifunctional CCA protein | \|CCA_BURCJ | 1.83 | 413 | 7 |
| nuoF | NADH dehydrogenase I chain F | \|B4E5L7_BURCJ | 1.82 | 436 | 2 |
| BCAL3032 | Putative ATPase protein | \|B4EBR6_BURCJ | 1.82 | 441 | 12 |
| BCAM0142 | Putative acyl-CoA dehydrogenase family protein | \|B4EH42_BURCJ | 1.81 | 414 | 6 |
| BCAL2984 | Cysteine peptidase, family C26 | \|B4EB84_BURCJ | 1.81 | 396 | 3 |
| BCAL1829 | Putative outer membrane protein | \|B4EAC8_BURCJ | 1.81 | 193 | 1 |
| guaD | Guanine deaminase | \|B4EB90_BURCJ | 1.80 | 439 | 5 |
| BCAL3318 | Putative RNA-methylase protein | \|B4EE39_BURCJ | 1.80 | 433 | 1 |
| polA | DNA polymerase I | \|B4EN22_BURCJ | 1.80 | 917 | 9 |
| vioA | Nucleotide sugar aminotransferase | \|B4ECE9_BURCJ | 1.79 | 380 | 1 |
| BCAL0537 | Endonuclease/exonuclease/phosphatase family protein | \|B4E8I0_BURCJ | 1.79 | 271 | 1 |
| bkdA1 | 2-oxoisovalerate dehydrogenase alpha subunit | \|B4EEE9_BURCJ | 1.78 | 410 | 10 |
| BCAL2446 | Putative aminotransferase | \|B4E6H2_BURCJ | 1.78 | 393 | 4 |
| mraZ | Protein MraZ | \|MRAZ_BURCJ | 1.78 | 142 | 0 |
| BCAS0051 | Putative glycerol utilisation-related protein | \|B4EPG8_BURCJ | 1.78 | 566 | 6 |
| BCAM0903 | Lysine decarboxylase family protein | \|B4EN21_BURCJ | 1.78 | 247 | 16 |
| purA | Adenylosuccinate synthetase | \|B4EAH2_BURCJ | 1.77 | 448 | 7 |
| phbC | Poly-beta-hydroxybutyrate polymerase | \|B4EAG2_BURCJ | 1.77 | 621 | 1 |
| clpB | Putative type VI secretion system protein TssH | \|B4E6R9_BURCJ | 1.77 | 889 | 2 |
| BCAM2565 | Putative methyltransferase | \|B4EJR4_BURCJ | 1.76 | 285 | 6 |
| BCAL0306 | Uncharacterized protein | \|B4E629_BURCJ | 1.76 | 91 | 0 |
| clpB | ClpB heat-shock protein | \|B4EB05_BURCJ | 1.76 | 865 | 6 |
| rplR | 50S ribosomal protein L18 | \|RL18_BURCJ | 1.76 | 121 | 5 |
| BCAL3193 | Uncharacterized protein | \|B4ECY4_BURCJ | 1.75 | 100 | 3 |

| | | | | | |
|---|---|---|---|---|---|
| BCAL2980 | Putative oxygenase | \|B4EB79_BURCJ | 1.75 | 548 | 2 |
| rplD | 50S ribosomal protein L4 | \|RL4_BURCJ | 1.74 | 206 | 5 |
| glcB | Malate synthase G | \|B4ELZ9_BURCJ | 1.74 | 724 | 6 |
| BCAL0946 | LysR family regulatory protein | \|B4EBX6_BURCJ | 1.74 | 324 | 3 |
| hctB | Histone H1-like protein | \|B4E5V7_BURCJ | 1.73 | 216 | 0 |
| glt2 | Glutamate synthase small subunit | \|B4E613_BURCJ | 1.73 | 488 | 13 |
| waaF | Putative ADP-heptose--LPS heptosyltransferase II | \|B4EBZ7_BURCJ | 1.73 | 345 | 4 |
| rplX | 50S ribosomal protein L24 | \|RL24_BURCJ | 1.72 | 102 | 5 |
| rpsO | 30S ribosomal protein S15 | \|RS15_BURCJ | 1.72 | 89 | 5 |
| ackA | Acetate kinase | \|B4EIA5_BURCJ | 1.71 | 399 | 3 |
| gatB | Aspartyl/glutamyl-tRNA(Asn/Gln) amidotransferase subunit B | \|GATB_BURCJ | 1.71 | 491 | 3 |
| BCAL2958/omp/ | Putative ompA family protein | \|B4EB48_BURCJ | 1.70 | 222 | 13 |
| BCAM2813 | Uncharacterized protein | \|B4ELZ1_BURCJ | 1.69 | 134 | 0 |
| dnaX | DNA polymerase III subunit gamma | \|B4EAY4_BURCJ | 1.69 | 787 | 11 |
| astD | N-succinylglutamate 5-semialdehyde dehydrogenase | \|ASTD_BURCJ | 1.69 | 487 | 4 |
| BCAM0300 | Metallo-beta-lactamase superfamily protein | \|B4EIB2_BURCJ | 1.69 | 467 | 12 |
| BCAL0893 | Phosphotransferase enzyme family protein | \|B4EBE4_BURCJ | 1.68 | 349 | 1 |
| BCAM1537 | Putative dehydrogenase, zinc-binding subunit | \|B4EJM7_BURCJ | 1.68 | 352 | 8 |
| BCAL0736 | Phosphoenolpyruvate-protein phosphotransferase | \|B4EA61_BURCJ | 1.68 | 590 | 8 |
| bceK | Glycosyltransferase | \|B4EMY3_BURCJ | 1.68 | 388 | 7 |
| paaA | Phenylacetic acid degradation protein PaaA | \|B4E5A2_BURCJ | 1.68 | 332 | 3 |
| BCAL3332 | Aminopeptidase P | \|B4EE53_BURCJ | 1.68 | 461 | 3 |
| panB | 3-methyl-2-oxobutanoate hydroxymethyltransferase | \|B4EDI3_BURCJ | 1.67 | 271 | 5 |
| dnaJ | Chaperone protein DnaJ | \|DNAJ_BURCJ | 1.66 | 378 | 9 |
| BCAL1054 | Putative lipoprotein | \|B4ECL6_BURCJ | 1.66 | 173 | 0 |
| BCAM1352 | Putative phosphoesterase | \|B4EIC8_BURCJ | 1.66 | 592 | 6 |
| BCAL1966 | tRNA-modifying protein YgfZ | \|B4EBJ4_BURCJ | 1.65 | 344 | 6 |
| leuC1 | 3-isopropylmalate dehydratase large subunit | \|B4EFJ1_BURCJ | 1.65 | 469 | 6 |
| radA | DNA repair protein radA | \|B4ECP9_BURCJ | 1.65 | 458 | 4 |
| BCAL2986 | Putative amidotransferase | \|B4EB86_BURCJ | 1.65 | 458 | 1 |
| BCAL1979 | Putative fatty acid degradation protein | \|B4EBK6_BURCJ | 1.64 | 700 | 21 |
| ilvA | Threonine dehydratase biosynthetic | \|B4EBC0_BURCJ | 1.64 | 507 | 9 |
| BCAL0038 | Putative long-chain-fatty-acid--CoA ligase | \|B4EEZ1_BURCJ | 1.63 | 575 | 4 |

| | | | | | |
|---|---|---|---|---|---|
| BCAL2745 | Putative CoA transferase family protein | \|B4E9E4_BURCJ | 1.63 | 465 | 6 |
| yidC | Membrane protein insertase YidC | \|YIDC_BURCJ | 1.63 | 552 | 4 |
| fumC | Fumarate hydratase class II | \|B4EAL6_BURCJ | 1.62 | 464 | 7 |
| pBCA001 | Putative partition protein | \|B3KYE6_BURCJ | 1.62 | 223 | 0 |
| BCAM0310 | Putative ribonucleotide reductase | \|B4EIQ1_BURCJ | 1.62 | 608 | 2 |
| thyA | Thymidylate synthase | \|B4EAM0_BURCJ | 1.62 | 323 | 12 |
| BCAL2410 | UPF0176 protein BceJ2315_23700 | \|Y2370_BURCJ | 1.62 | 284 | 0 |
| BCAL0698 | ABC transporter ATP-binding protein | \|B4E9P8_BURCJ | 1.61 | 304 | 2 |
| BCAL2271 | Putative toxic anion resistance protein | \|B4EEL0_BURCJ | 1.61 | 365 | 0 |
| manC | Putative GDP-mannose pyrophosphorylase | \|B4EDG4_BURCJ | 1.61 | 475 | 9 |
| trpA | Tryptophan synthase alpha chain | \|TRPA_BURCJ | 1.61 | 271 | 3 |
| BCAL1927 | Putative aminotransferase | \|B4EB13_BURCJ | 1.61 | 412 | 8 |
| cysN | Putative sulfate adenylyltransferase subunit 1 | \|B4E8V4_BURCJ | 1.61 | 438 | 5 |
| BCAL2644 | Putative ATP-binding protein | \|B4E8D8_BURCJ | 1.60 | 363 | 5 |
| hisD | Histidinol dehydrogenase | \|B4E635_BURCJ | 1.60 | 438 | 6 |
| atpG | ATP synthase gamma chain | \|ATPG_BURCJ | 1.60 | 291 | 10 |
| BCAL0307 | ABC transporter ATP-binding protein | \|B4E630_BURCJ | 1.60 | 308 | 11 |
| BCAL0123 | Putative glycosyltransferase | \|B4EDJ7_BURCJ | 1.60 | 354 | 4 |
| tex | Putative transcription accessory protein | \|B4EAG9_BURCJ | 1.60 | 774 | 3 |
| BCAL1468 | \|B4E7H3_BURCJ Putative electron transport protein GN=BCAL1468 PE=4 SV=1; | \|B4E7H3_BURCJ | 1.59 | 557 | 12 |
| BCAL3286 | Cobalamin adenosyltransferase protein | \|B4EE08_BURCJ | 1.59 | 183 | 2 |
| BCAL1331 | Putative aldehyde dehydrogenase | \|B4E652_BURCJ | 1.59 | 774 | 4 |
| BCAL1843 | ABC transporter ATP-binding protein | \|B4EAE2_BURCJ | 1.59 | 530 | 22 |
| pykA | Pyruvate kinase | \|B4EA24_BURCJ | 1.59 | 478 | 4 |
| rpmA | 50S ribosomal protein L27 | \|RL27_BURCJ | 1.59 | 87 | 4 |
| fusA | Elongation factor G | \|B4E912_BURCJ | 1.58 | 703 | 8 |
| pyrB | Putative aspartate carbomyltransferase | \|B4EN51_BURCJ | 1.58 | 431 | 3 |
| fusA | Elongation factor G | \|B4E5B7_BURCJ | 1.58 | 700 | 8 |
| atpA | ATP synthase subunit alpha | \|ATPA_BURCJ | 1.58 | 513 | 10 |
| BCAL0027 | Chromosome partitioning protein ParB | \|B4E584_BURCJ | 1.58 | 297 | 1 |
| bfr | Bacterioferritin | \|B4EEM7_BURCJ | 1.58 | 158 | 7 |
| BCAM2468 | Putative aldehyde dehydrogenase family protein | \|B4EIN4_BURCJ | 1.58 | 503 | 15 |
| gor | Glutathione reductase | \|B4E8M8_BURCJ | 1.58 | 451 | 14 |

| | | | | | |
|---|---|---|---|---|---|
| BCAL1885 | Putative membrane protein | \|B4EAX0_BURCJ | 1.57 | 338 | 6 |
| dnaA | Chromosomal replication initiator protein DnaA | \|DNAA_BURCJ | 1.57 | 525 | 8 |
| tdh | L-threonine 3-dehydrogenase | \|TDH_BURCJ | 1.57 | 342 | 6 |
| BCAL0858 | Putative exported protein | \|B4EBA9_BURCJ | 1.56 | 144 | 0 |
| serC | Phosphoserine aminotransferase | \|SERC_BURCJ | 1.56 | 360 | 5 |
| rplU | 50S ribosomal protein L21 | \|RL21_BURCJ | 1.56 | 103 | 5 |
| BCAL2018 | Putative acetylornithine deacetylase | \|B4EC25_BURCJ | 1.56 | 406 | 11 |
| rplE | 50S ribosomal protein L5 | \|RL5_BURCJ | 1.56 | 179 | 5 |
| BCAL2287 | Putative fumarate hydratase | \|B4EEM6_BURCJ | 1.56 | 507 | 3 |
| rimO | Ribosomal protein S12 methylthiotransferase RimO | \|RIMO_BURCJ | 1.56 | 453 | 5 |
| BCAL1856 | Putative thiamine biosynthesis oxidoreductase ThiO | \|B4E620_BURCJ | 1.55 | 378 | 5 |
| murC | UDP-N-acetylmuramate--L-alanine ligase | \|MURC_BURCJ | 1.55 | 465 | 11 |
| hemL | Glutamate-1-semialdehyde 2,1-aminomutase | \|GSA_BURCJ | 1.54 | 427 | 6 |
| hisC | Histidinol-phosphate aminotransferase | \|B4EAR0_BURCJ | 1.54 | 355 | 6 |
| nuoI | NADH-quinone oxidoreductase subunit I | \|NUOI_BURCJ | 1.54 | 162 | 6 |
| BCAL0704 | D-alanyl-D-alanine carboxypeptidase (Penicillin-binding protein) | \|B4E9Q4_BURCJ | 1.53 | 436 | 9 |
| BCAL221 | Oligopeptidase A | \|B4EDU2_BURCJ | 1.53 | 695 | 9 |
| BCAL1660 | Putative ribose operon repressor | \|B4E8T3_BURCJ | 1.53 | 343 | 2 |
| sdaA | L-serine dehydratase I | \|B4EF24_BURCJ | 1.52 | 462 | 6 |
| pheA | P-protein [bifunctional includes: chorismate mutase and prephenate dehydrata | \|B4EB44_BURCJ | 1.52 | 360 | 3 |
| argM | Acetylornithine aminotransferase | \|B4ECZ5_BURCJ | 1.52 | 397 | 6 |
| BCAL1049 | Luciferase-like monooxygenase | \|B4ECL1_BURCJ | 1.52 | 332 | 2 |
| BCAL1061 | Putative arginine N-succinyltransferase, beta chain | \|B4ECZ7_BURCJ | 1.52 | 343 | 3 |
| BCAL1733 | Putative glutathione S-transferase | \|B4E9D2_BURCJ | 1.52 | 256 | 2 |
| BCAL0883 | TetR family regulatory protein | \|B4EBD4_BURCJ | 1.52 | 199 | 0 |
| dnaB | Replicative DNA helicase | \|B4EB27_BURCJ | 1.52 | 460 | 6 |
| BCAL2993 | Aminopeptidase N | \|B4EB93_BURCJ | 1.51 | 897 | 8 |
| lipB | Octanoyltransferase | \|B4E9Q9_BURCJ | 1.51 | 220 | 4 |
| BCAL1957 | Putative sugar transferase | \|B4EBI5_BURCJ | 1.51 | 354 | 4 |
| BCAL1819 | Uncharacterized protein | \|B4EAB8_BURCJ | 1.51 | 556 | 4 |
| hemB | Delta-aminolevulinic acid dehydratase | \|B4E5Y8_BURCJ | 1.50 | 332 | 5 |
| BCAL1065 | Periplasmic solute-binding protein | \|B4ED01_BURCJ | 1.50 | 264 | 4 |
