## supplemental Table S4 for "The DNA mimic protein BCAS0292 is involved in the regulation of virulence of *Burkholderia cenocepacia*"

**Suppl Table S4: Proteins encoded by Lxa- locus genes absent in the BCAS0292 mutant and also altered in chronic infection B. cenocepacia isolates.**

| Protein name | Gene ID | UniProt Code | Sequence length |
|---|---|---|---|
| Universal stress protein | BCAM0276 | \|B4EI87_BURCJ | 156 |
| Uncharacterized protein | BCAM0277 | \|B4EI88_BURCJ | 91 |
| Putative heat shock protein | BCAM0278 | \|B4EI89_BURCJ | 144 |
| Putative phospholipid-binding protein | BCAM0280 | \|B4EI91_BURCJ | 216 |
| Putative cytochrome c | BCAM0284 | \|B4EI96_BURCJ | 111 |
| Uncharacterized protein | BCAM0285 | \|B4EI97_BURCJ | 335 |
| Putative universal stress protein | BCAM0290 | \|B4EIA2_BURCJ | 156 |
| Putative universal stress protein | BCAM0291 | \|B4EIA3_BURCJ | 277 |
| Putative universal stress protein | BCAM0292 | \|B4EIA4_BURCJ | 167 |
| Putative universal stress protein | BCAM0294 | \|B4EIA6_BURCJ | 279 |
| Uncharacterized protein | BCAM0308 | \|B4EIC0_BURCJ | 169 |
| Uncharacterized protein | BCAM0316 | \|B4EIQ7_BURCJ | 157 |
| Acetoacetyl-CoA reductase | BCAM0296 phbB | \|B4EIA8_BURCJ | 248 |
