## supplemental table S5 for "The DNA mimic protein BCAS0292 is involved in the regulation of virulence of *Burkholderia cenocepacia*"

Supplemental Table 5: Proteins that were reduced in abundance or undetectable in the ΔBCAS0292 mutant and increased in abundance in late chronic infe

| Actual fold difference | t-test Significant | t-test p value | BCA ID | Protein names |
|---|---|---|---|---|
| 2.2 | + | 1.60E-05 | L0043 | \|B4EF03_BURCJ Putative extracellular ligand-binding protein OS=Burkholderia |
| 1.5 | + | 0.00179054 | L0118 tag | \|B4EDJ2_BURCJ DNA-3-methyladenine glycosylase I OS=Burkholderia cenocep |
| 1.6 | + | 0.0194004 | L0120 | \|B4EDJ4_BURCJ Haloacid dehalogenase-like hydrolase OS=Burkholderia cenoc |
| 1.5 | + | 0.000286742 | L0211 | \|B4E597_BURCJ Uncharacterized protein OS=Burkholderia cenocepacia (strair |
| 1.9 | + | 2.17E-05 | L0343 bcsL | \|B4E6R5_BURCJ Putative type VI secretion system protein TssD OS=Burkholde |
| 2.8 | + | 4.42E-06 | L0378 | \|B4E6V1_BURCJ Putative hydrolase OS=Burkholderia cenocepacia (strain ATC( |
| 1.5 | + | 0.00580938 | L0391 acpD | \|B4E6W4_BURCJ FMN-dependent NADH-azoreductase OS=Burkholderia ceno |
| 2.0 | + | 0.00331672 | L0394 ung | \|B4E6W7_BURCJ Uracil-DNA glycosylase OS=Burkholderia cenocepacia (strain |
| 3.6 | + | 0.000666689 | L0395 | \|B4E7A6_BURCJ Putative adenylate cyclase OS=Burkholderia cenocepacia (str |
| 2.2 | + | 3.70E-06 | L0396 trpC | \|B4E7A7_BURCJ Indole-3-glycerol phosphate synthase OS=Burkholderia cenoc |
| 1.6 | + | 0.00704941 | L0463 | \|B4E7V5_BURCJ Putative thioredoxin OS=Burkholderia cenocepacia (strain AT |
| 1.6 | + | 0.000291238 | L0557 | \|B4E8K0_BURCJ Putative glutathione S-transferase OS=Burkholderia cenocepa |
| 2.9 | + | 4.10E-05 | L0690 | \|B4E9P0_BURCJ Uncharacterized protein OS=Burkholderia cenocepacia (strair |
| 2.1 | + | 0.0222858 | L0691 | \|B4E9P1_BURCJ Putative cytidylyltransferase OS=Burkholderia cenocepacia (s |
| 3.1 | + | 0.000126083 | L0702 | \|B4E9Q2_BURCJ Putative ferredoxin OS=Burkholderia cenocepacia (strain ATC |
| 1.6 | + | 0.00299335 | L0728 | \|B4EA54_BURCJ Uncharacterized protein OS=Burkholderia cenocepacia (strair |
| 2.3 | + | 1.17E-05 | L0734 | \|B4EA59_BURCJ Sugar transport PTS system IIa component OS=Burkholderia c |
| 2.5 | + | 2.05E-05 | L0789 | \|B4EAQ2_BURCJ Uncharacterized protein OS=Burkholderia cenocepacia (strai |
| 1.7 | + | 9.37E-05 | L0812 | \|B4EAS3_BURCJ Sigma-54 modulation protein OS=Burkholderia cenocepacia (s |
| 1.8 | + | 0.00470324 | L0814 | \|B4EAS5_BURCJ ABC transporter ATP-binding protein OS=Burkholderia cenoce |
| 1.5 | + | 0.00476182 | L0830 | \|B4EAU1_BURCJ Putative ParA family protein OS=Burkholderia cenocepacia (s |
| 3.3 | + | 4.53E-05 | L0831 | \|B4EAU2_BURCJ Putative storage protein OS=Burkholderia cenocepacia (strair |
| 3.0 | + | 0.0121219 | L0849 | \|B4EAW0_BURCJ Metallo peptidase, subfamily M48B OS=Burkholderia cenoce |
| 2.3 | + | 0.000905646 | L0871 | \|B4EBC2_BURCJ Uncharacterized protein OS=Burkholderia cenocepacia (strair |
| 1.7 | + | 0.000220708 | L0970 | \|B4EAK8_BURCJ CreA protein 1 OS=Burkholderia cenocepacia (strain ATCC BA |
| 1.6 | + | 0.00789761 | L0971 | \|B4EAK7_BURCJ 4Fe-4S ferredoxin OS=Burkholderia cenocepacia (strain ATCC |
| 1.9 | + | 2.27E-05 | L1071 | \|B4ED07_BURCJ NAD dependent epimerase/dehydratase family protein OS=B |
| 3.1 | + | 0.000485775 | L1399 | \|B4E6B9_BURCJ OsmC-like protein OS=Burkholderia cenocepacia (strain ATCC |
| 1.7 | + | 0.000569597 | L1465 | \|B4E7H0_BURCJ Uncharacterized protein OS=Burkholderia cenocepacia (strair |
| 2.1 | + | 0.00122252 | L1864 | \|B4EAG3_BURCJ Uncharacterized protein OS=Burkholderia cenocepacia (strair |

| | | | | | | |
|---|---|---|---|---|---|---|
| | 3.3 | + | | 9.01E-06 | L1868 | \|B4EAG7_BURCJ Uncharacterized protein OS=Burkholderia cenocepacia (strai |
| | 3.2 | + | | 0.00180116 | L1910acoB | \|B4EAZ6_BURCJ Acetoin:2,6-dichlorophenolindophenol oxidoreductase beta s |
| | 1.8 | + | | 0.00134802 | L1920 | \|B4EB06_BURCJ Putative DNA-binding protein OS=Burkholderia cenocepacia ( |
| | 1.5 | + | | 0.00480682 | L1967 | \|B4EBJ5_BURCJ Uncharacterized protein OS=Burkholderia cenocepacia (strain |
| | 1.6 | + | | 0.00173613 | L2037 allA | \|B4EC44_BURCJ Ureidoglycolate lyase OS=Burkholderia cenocepacia (strain A |
| | 1.7 | + | | 0.00214961 | L2039 | \|B4EC46_BURCJ Putative uricase OS=Burkholderia cenocepacia (strain ATCC B |
| | 3.2 | + | | 1.23E-05 | L2123 | \|B4ECR4_BURCJ Uncharacterized protein OS=Burkholderia cenocepacia (strai |
| | 1.8 | + | | 0.00101254 | L2153 ppiB | \|B4ED76_BURCJ Peptidyl-prolyl cis-trans isomerase OS=Burkholderia cenocep |
| Not detected | | | 1 | 1 | L2166 | \|B4ED88_BURCJ Putative lipoprotein OS=Burkholderia cenocepacia (strain AT |
| | 1.8 | + | | 0.0209345 | L2175 | \|B4ED97_BURCJ Uncharacterized protein OS=Burkholderia cenocepacia (strai |
| | 2.5 | + | | 0.000162973 | L2237 | \|B4EDW6_BURCJ Putative aminotransferase OS=Burkholderia cenocepacia (st |
| | 2.1 | + | | 0.00119508 | L2324 | \|B4EER2_BURCJ Uncharacterized protein OS=Burkholderia cenocepacia (strai |
| | 2.0 | + | | 0.000148219 | L2325 | \|B4EER3_BURCJ Uncharacterized protein OS=Burkholderia cenocepacia (strai |
| | 2.1 | + | | 0.0122724 | L2328 | \|B4EER6_BURCJ Uncharacterized protein OS=Burkholderia cenocepacia (strai |
| | 1.6 | + | | 0.00611635 | L2373 | \|B4E5Q1_BURCJ Putative globin OS=Burkholderia cenocepacia (strain ATCC B |
| | 1.6 | + | | 0.000736016 | L2428 | \|B4E6F4_BURCJ Putative cytochrome C-related protein OS=Burkholderia ceno |
| | 1.8 | + | | 0.00229316 | L2464 | \|B4E730_BURCJ Short chain dehydrogenase OS=Burkholderia cenocepacia (st |
| | 2.1 | + | | 7.99E-05 | L2694 | \|B4E8W7_BURCJ Putative dehydrogenase OS=Burkholderia cenocepacia (strai |
| | 2.2 | + | | 0.00972957 | L2705 | \|B4E8X9_BURCJ ABC transporter ATP-binding protein OS=Burkholderia cenoce |
| | 2.2 | + | | 9.80E-05 | L2711 | \|B4E8Y5_BURCJ Peptidyl-prolyl cis-trans isomerase OS=Burkholderia cenocep |
| | 1.7 | + | | 0.000488227 | L2743 | \|B4E9E2_BURCJ Putative aldo/keto reductase family oxidoreductase OS=Burk |
| | 1.8 | + | | 0.000162405 | L3147 groS | \|B4ECT8_BURCJ 10 kDa chaperonin OS=Burkholderia cenocepacia (strain ATC |
| | 2.1 | + | | 6.33E-05 | L3319 | \|B4EE40_BURCJ Uncharacterized protein OS=Burkholderia cenocepacia (strai |
| | 1.5 | + | | 0.0428484 | L3328 | \|B4EE49_BURCJ Putative hydrolase OS=Burkholderia cenocepacia (strain ATC |
| | 1.9 | + | | 0.00409385 | L3398 | \|B4E5S8_BURCJ Putative competence-damaged related protein OS=Burkholde |
| | 1.9 | + | | 0.00103657 | L3413 aroE | \|B4E5U3_BURCJ Shikimate dehydrogenase OS=Burkholderia cenocepacia (stra |
| | 1.6 | + | | 0.000749459 | L3424 tpx | \|B4E5V4_BURCJ Probable thiol peroxidase OS=Burkholderia cenocepacia (stra |
| | 1.7 | + | | 0.00269053 | M0001 | \|B3KYE4_BURCJ Putative sigma factor OS=Burkholderia cenocepacia (strain AT |
| Not detected | | | 1 | 1 | M0016 | \|B4EG02_BURCJ Tartrate dehydrogenase OS=Burkholderia cenocepacia (strai |
| | 1.9 | + | | 1.09E-05 | M0050 | \|B4EG36_BURCJ Universal stress protein OS=Burkholderia cenocepacia (strain |
| | 2.7 | + | | 0.00179811 | M0165 | \|B4EH65_BURCJ Uncharacterized protein OS=Burkholderia cenocepacia (strai |
| | 1.6 | + | | 0.0675938 | M0191 | \|B4EHM3_BURCJ Putative non-ribosomal peptide synthetase OS=Burkholderi |
| Not detected | | | 1 | 1 | M0264 | \|B4EI75_BURCJ Putative membrane protein OS=Burkholderia cenocepacia (st |

| | | | | | |
|---|---|---|---|---|---|
| 2.0 | + | 2.42E-05 | M0276 | | \|B4EI87_BURCJ Universal stress protein OS=Burkholderia cenocepacia (strain |
| 2.6 | + | 0.000142127 | M0277 | | \|B4EI88_BURCJ Uncharacterized protein OS=Burkholderia cenocepacia (strain |
| 276.0 | + | 4.41E-09 | M0278 | | \|B4EI89_BURCJ Putative heat shock protein OS=Burkholderia cenocepacia (str |
| 1.5 | + | 0.0123741 | M0280 | | \|B4EI91_BURCJ Putative phospholipid-binding protein OS=Burkholderia cenoc |
| 3.3 | + | 0.00501421 | M0284 | | \|B4EI96_BURCJ Putative cytochrome c OS=Burkholderia cenocepacia (strain A |
| 4.3 | + | 7.86E-08 | M0285 | | \|B4EI97_BURCJ Uncharacterized protein OS=Burkholderia cenocepacia (strain |
| 2.2 | + | 0.00021621 | M0290 | | \|B4EIA2_BURCJ Putative universal stress protein OS=Burkholderia cenocepaci |
| 1.7 | + | 6.96E-05 | M0291 | | \|B4EIA3_BURCJ Putative universal stress protein OS=Burkholderia cenocepaci |
| 6.5 | + | 6.36E-08 | M0292 | | \|B4EIA4_BURCJ Putative universal stress protein OS=Burkholderia cenocepaci |
| 1.7 | + | 7.01E-05 | M0294 | | \|B4EIA6_BURCJ Putative universal stress protein OS=Burkholderia cenocepaci |
| 2.4 | + | 9.35E-06 | M0296 | phbB | \|B4EIA8_BURCJ Acetoacetyl-CoA reductase OS=Burkholderia cenocepacia (stra |
| 6.3 | + | 5.03E-08 | M0308 | | \|B4EIC0_BURCJ Uncharacterized protein OS=Burkholderia cenocepacia (strain |
| 2.1 | + | 0.00353838 | M0316 | | \|B4EIQ7_BURCJ Uncharacterized protein OS=Burkholderia cenocepacia (strain |
| 4.0 | + | 2.99E-05 | M0679 | | \|B4ELN0_BURCJ Uncharacterized protein OS=Burkholderia cenocepacia (strain |
| 5.5 | + | 1.73E-05 | M0944 | | \|B4EF89_BURCJ Putative lipoprotein OS=Burkholderia cenocepacia (strain ATC |
| 2.0 | + | 0.00106757 | M0979 | | \|B4EFC4_BURCJ Putative glutathione-S-transferase OS=Burkholderia cenocepa |
| 1.8 | + | 0.000335702 | M1217 | ahpC | \|B4EH97_BURCJ Alkyl hydroperoxide reductase subunit C OS=Burkholderia ce |
| 12.5 | + | 7.51E-07 | M1261 | | \|B4EHE0_BURCJ Putative membrane protein OS=Burkholderia cenocepacia (st |
| 2.2 | + | 0.00731098 | M1291 | | \|B4EHT1_BURCJ L-asparaginase OS=Burkholderia cenocepacia (strain ATCC BA |
| 2.0 | + | 0.000460699 | M1351 | | \|B4EIC7_BURCJ Putative regulatory protein OS=Burkholderia cenocepacia (stra |
| 2.0 | + | 0.000154496 | M1363 | | \|B4EID9_BURCJ Uncharacterized protein OS=Burkholderia cenocepacia (strain |
| 4.1 | + | 0.000487859 | M1412 | | \|B4EII8_BURCJ Uncharacterized protein OS=Burkholderia cenocepacia (strain |
| Not detected | 1 | 1 | M1414 | | \|B4EIJ0_BURCJ Uncharacterized protein OS=Burkholderia cenocepacia (strain |
| 3.2 | + | 4.81E-05 | M1465 | | \|B4EJ19_BURCJ Putative exported protein OS=Burkholderia cenocepacia (strai |
| 2.3 | + | 0.000191023 | M1480 | | \|B4EJ33_BURCJ Uncharacterized protein OS=Burkholderia cenocepacia (strain |
| 2.4 | + | 0.000375423 | M1481 | | \|B4EJ34_BURCJ Uncharacterized protein OS=Burkholderia cenocepacia (strain |
| 2.8 | + | 1.29E-05 | M1482 | | \|B4EJ35_BURCJ Uncharacterized protein OS=Burkholderia cenocepacia (strain |
| 3.0 | + | 5.26E-07 | M1495 | | \|B4EJI6_BURCJ Universal stress protein OS=Burkholderia cenocepacia (strain |
| 2.2 | + | 0.001934 | M1496 | | \|B4EJI7_BURCJ Uncharacterized protein OS=Burkholderia cenocepacia (strain |
| 2.1 | + | 0.0770267 | M1502 | | \|B4EJJ2_BURCJ Uncharacterized protein OS=Burkholderia cenocepacia (strain |
| 2.1 | + | 0.00399262 | M1511 | | \|B4EJK1_BURCJ Uncharacterized protein OS=Burkholderia cenocepacia (strain |
| 2.6 | + | 6.30E-06 | M1823 | | \|B4EM86_BURCJ Putative methyltransferase OS=Burkholderia cenocepacia (st |
| 1.7 | + | 7.92E-05 | M2160 | | \|B4EFV7_BURCJ Two-component regulatory system, response regulator prote |

| | | | | | |
|---|---|---|---|---|---|
| 1.7 | + | 0.000255677 | M2461 | | \|B4EIM7_BURCJ Putative inosine-uridine preferring nucleoside hydrolase OS= |
| 1.5 | + | 0.0418282 | M2473 | | \|B4EIN9_BURCJ DeoR family regulatory protein OS=Burkholderia cenocepacia |
| 3.2 | + | 2.77E-06 | M2477 | | \|B4EIP3_BURCJ Serine peptidase, family S10 OS=Burkholderia cenocepacia (st |
| 1.8 | + | 0.000194311 | M2509 | | \|B4EJ62_BURCJ Putative FucU/RbsD family transport protein OS=Burkholderia |
| 1.6 | + | 0.0617829 | M2569 | | \|B4EJR8_BURCJ IclR family regulatory protein OS=Burkholderia cenocepacia (s |
| 1.8 | + | 0.0036961 | M2742 | | \|B4ELE1_BURCJ Uncharacterized protein OS=Burkholderia cenocepacia (strain |
| 9.5 | + | 8.29E-07 | M2752 | | \|B4ELF1_BURCJ NAD dependent epimerase/dehydratase family protein OS=Bu |
| 1.9 | + | 0.000202769 | S0013 | | \|B4EQE1_BURCJ Putative molybdenum transport protein OS=Burkholderia cer |
| 1.9 | + | 8.99E-06 | S0257 | | \|B4EQ25_BURCJ Putative acetyltransferase OS=Burkholderia cenocepacia (stra |
| 1.8 | + | 0.000122424 | S0281 | | \|B4EQ47_BURCJ Putative 2-hydroxy-3-oxopropionate reductase OS=Burkholde |
| 49.8 | + | 7.06E-10 | S0293 | aidA | \|B4EQ59_BURCJ Nematocidal protein AidA OS=Burkholderia cenocepacia (stra |
| 29.0 | + | 2.73E-06 | S0636 | | \|B4EP30_BURCJ Uncharacterized protein OS=Burkholderia cenocepacia (strain |
| 1.6 | + | 0.000380517 | S0738 | | \|B4EPC6_BURCJ Putative short-chain dehydrogenase family protein OS=Burkh |

| |
|---|
| ı ATCC BAA-245 / DSM 16553 / LMG 16656 / NCTC 13227 / J2315 / CF5610) GN=L0690 PE=4 SV=1;>tr\|B4EPZ0\|B |
| urkholderia cenocepacia (strain ATCC BAA-245 / DSM 16553 / LMG 16656 / NCTC 13227 / J2315 / CF5610) GN=I |

subunit OS=Burkholderia cenocepacia (strain ATCC BAA-245 / DSM 16553 / LMG 16656 / NCTC 13227 / J2315 / C

rain ATCC BAA-245 / DSM 16553 / LMG 16656 / NCTC 13227 / J2315 / CF5610) GN=L2237 PE=3 SV=1;>tr|B4E92

holderia cenocepacia (strain ATCC BAA-245 / DSM 16553 / LMG 16656 / NCTC 13227 / J2315 / CF5610) GN=L274 ein OS=Burkholderia cenocepacia (strain ATCC BAA-245 / DSM 16553 / LMG 16656 / NCTC 13227 / J2315 / CF561

Burkholderia cenocepacia (strain ATCC BAA-245 / DSM 16553 / LMG 16656 / NCTC 13227 / J2315 / CF5610) GN=

urkholderia cenocepacia (strain ATCC BAA-245 / DSM 16553 / LMG 16656 / NCTC 13227 / J2315 / CF5610) GN=N

olderia cenocepacia (strain ATCC BAA-245 / DSM 16553 / LMG 16656 / NCTC 13227 / J2315 / CF5610) GN=S073
